# Unveiling the taxonomic diversity and unprecedented biosynthetic treasure of the phylum *Myxococcota*

**DOI:** 10.64898/2026.02.12.705523

**Authors:** Amay Ajaykumar Agrawal, Ronald Garcia, Guangyi Chen, Chantal D. Bader, Peter Sullivan, Alexander Popoff, Daniel Krug, Maja Hunter, Sebastian Walesch, Emilia Oueis, Christine Frank, Haowen Zhao, Joachim J. Hug, Sebastian Keller, Pascal Hirsch, Azat Tagirdzhanov, Tatiana Malygina, Mary Victory E. Gutierrez, Alexey Gurevich, Judith Boldt, Boyke Bunk, Jörg Overmann, Ulrich Nübel, Andreas Keller, Olga V. Kalinina, Rolf Müller

## Abstract

Natural products remain vital sources of therapeutics, particularly anti-infectives, and members of the phylum *Myxococcota* constitute an especially rich reservoir for their discovery. Based on decades of microbiological efforts, we present 154 new *Myxococcota* genomes and propose a revised taxonomy expanding the number of described families from 11 to 28 and genera from 32 to 90. Comparison with an equivalent set from the prime source *Actinomycetota* shows that *Myxococcota* possess a comparable biosynthetic diversity, underscoring their promise for large-scale isolation and sequencing efforts. The vast untapped potential reflected in 2,387 uncharacterized gene cluster families is highlighted by genome mining efforts, yielding four validated compounds exhibiting novel chemistry, including myxolutamids and myxopentacins. We show that many *Myxococcota*-derived natural products, such as myxolutamid A and two new sorangicin derivatives, are conserved within taxonomic lineages. New described families thus bear high biosynthetic potential underpinning the importance of precise taxonomic classification guiding targeted drug discovery.

## INTRODUCTION

*Myxococcota* is a phylum of Gram-negative bacteria ubiquitously found in soil and distributed in various other habitats around the globe. They are best known for their swarming, microbial predatory behavior, and formation of multicellular, complex tree-like fruiting bodies^1^. In the last few decades, bacteria from *Myxococcota* have increasingly gained attention because of their capacity for producing diverse and novel secondary metabolites, also known as natural products (NPs)^2–4^. *Myxococcota* are among the most challenging microorganisms to cultivate because of their slow growth and poorly understood metabolism^5–7^. Owing to this, the currently validly described *Myxococcota* phylum only consists of 11 families and 32 genera, distributed into three suborders: *Cystobacterineae, Nannocystineae*, and *Sorangiineae*^8^ (accessed on 01.06.2024). Most of the currently isolated *Myxococcota* belong to the genera *Myxococcus, Corallococcus*, and *Pyxidicoccus*^9–11^, while the other taxa remain largely only identified by metagenomics without cultured representatives, and are understudied^12^. Thus, only the “tip of the iceberg” of the enormous *Myxococcota* diversity has been explored until now^13,14^.

For decades, NPs have been instrumental in drug discovery by providing biologically active chemical scaffolds, with most of the clinically used antibiotics being their derivatives or featuring a NP pharmacophore^15^. With the growing menace of antimicrobial resistance^16^, it becomes of paramount importance to fill the drug discovery pipeline with novel candidate molecules, preferably featuring new mechanisms of action^17^. While the majority of marketed antibiotics have been developed from NPs belonging to the genus *Streptomyces* of the phylum *Actinomycetota*^18^, *Myxococcota* present an appealing alternative source of biologically active compounds. *Myxococcota* NPs frequently exhibit novel chemical scaffolds and, consequently, unique modes of action^4^. The search for alternative NP producers has become increasingly important, as the high rediscovery rate in *Actinomycetota* has emerged as a major bottleneck for drug discovery, whereas *Myxococcota* predominantly yield chemically distinct metabolites^3^. Some striking examples of NPs from *Myxococcota* with pharmaceutical importance are the antibiotics cystobactamids^19–21^ (isolated from *Cystobacter velatus* and acting as topoisomerase inhibitors) and corallopyronin^22,23^ (isolated from *Corallococus coralloides* and acting as RNA polymerase inhibitors), while semisynthetic derivatives of epothilone^24^ (isolated from *Sorangium cellulosum*) have been approved by the FDA to treat breast cancer in 2007^25^.

Most microbial NPs are produced by the expression of genomic loci called biosynthetic gene clusters (BGCs) encoding the respective complex biosynthetic machineries for each NP. Discovering these loci and connecting them to their small-molecule products, referred to as genome mining, is a key step in drug discovery from microorganisms^26^. Over the past years, genome mining has both played a crucial role in uncovering new NPs from *Myxococcota* and enabled the assignment of BGCs to known metabolites, improving insights into their evolution and supporting structure elucidation (Figure 1A). These efforts led to the discovery of the antifungal myxoquaterins, antiviral sandacrabins, new sorangibactin siderophores, antibacterial and cytotoxic sandarazols, and the pyxidicyclines as new topoisomerase inhibitors, to name a few^27–30^. However, although *Myxococcota* harbour the largest bacterial genomes known to date, with a substantial proportion of them dedicated to BGCs, the genome-encoded biosynthetic potential of most *Myxococcota* strains remains largely unexplored^31,32^. Currently, the RefSeq database^33^ only contains 218 *Myxococcota* genomes (accessed on 31.08.2024), most of which are heavily fragmented. This poses a significant challenge for connecting NPs with the corresponding BGCs in these strains, since, due to their genomic make-up and sheer size, BGCs often occur at contig edges and are incomplete in such assemblies. Thus, even though past discoveries of *Myxococcota* NPs often proceeded without complete genomes, obtaining high-quality genome sequences has become an important prerequisite for reliably connecting metabolites to their BGCs and for expanding genome-guided drug discovery.

**Figure 1.**
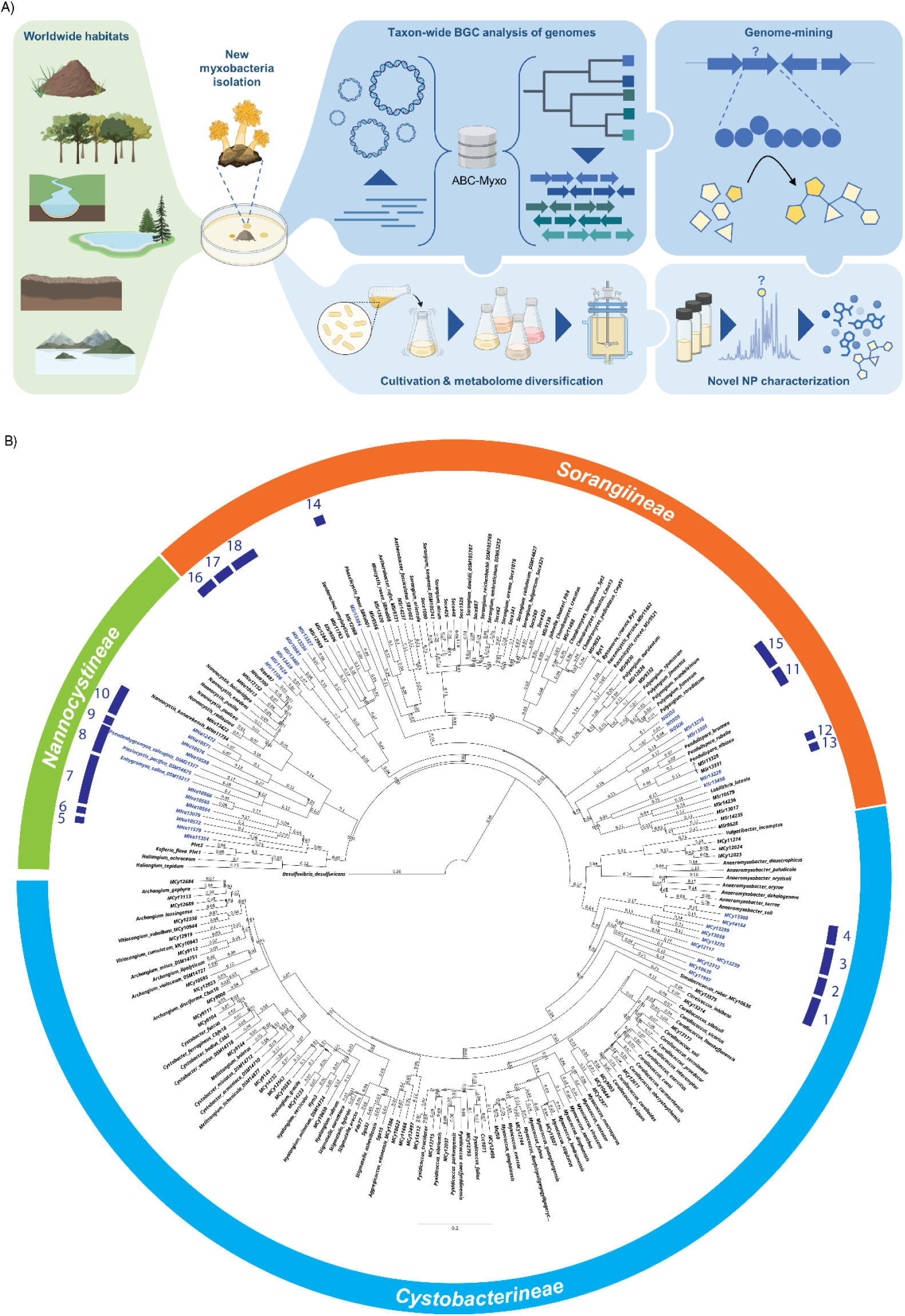
NP discovery framework and taxonomic refinement for the *Myxococcota* phylum. **(A)** Overview of the key steps and their integration in the NP discovery workflows pursued in this study. A curated collection of biosynthetic gene clusters predicted in the high-quality genomes presented in this study forms the foundation for exploring the diverse secondary metabolomes of newly isolated *Myxococcota*. ABC-Myxo = database of genomes and BGCs presented in this study, available online at https://tools.helmholtz-hips.de/abc_myxo/. **(B)** Phylogenomic reconstruction and taxonomic overhaul of the *Myxococcota* phylum by 37 type strains and 117 newly isolated strains are reported in this study. Genomes corresponding to newly introduced families are highlighted in blue, with type strains labeled by species names. Leaves annotated with both species names and strain identifiers denote novel genomes proposed as type strain representatives. Order-level taxonomy is indicated in the outer circle, new families in the inner blue circle, with numbers corresponding to family indices. *Desulfovibrio desulfuricans* serves as the outgroup.

While the biosynthetic potential of *Myxococcota* has not yet been comprehensively characterized at the genomic level, our previous study has tackled this question from a metabolomics perspective^34^. In this study, high-resolution mass spectrometry datasets from ∼2300 *Myxococcota* strains were subjected to a statistical analysis pipeline that revealed a clear correlation between taxonomic distance of strains (assigned in this case based on morphological features and 16S rRNA sequences) and the diversity of their metabolome profiles. While this study painted a clear picture confirming previous empirical observations, its informative value is limited by a bias towards NPs expressed under laboratory conditions.

Genome-level analyses will be key to uncovering the hidden biosynthetic diversity beyond these conditions; however, they are particularly challenging in *Myxococcota* not only due to their large genome sizes and high GC content, requiring, for example, special biochemical procedures for DNA isolation. Additionally, *Myxococcota* isolates are often extremely slow-growing and require particular conditions for isolation and cultivation^6,9,35–37^.

In this study, based on decades of sample collection and isolation efforts, we present 154 newly sequenced genomes of *Myxococcota* strains that were carefully picked to represent a maximum of morphological and taxonomic diversity, substantially expanding the available high-quality genomic resources and cultured biodiversity. We explore how refined taxonomic frameworks can inform NP research and assess the conservation of secondary metabolites across different lineages. The potential biosynthetic capacity of the newly sequenced strains is comprehensively evaluated, with its capacity for uncovering novel chemistry illustrated by the identification of two NP classes, including a new non-ribosomal peptide (NRP) family. Finally, we compare the overall biosynthetic diversity of *Myxococcota* with other previously extensively studied bacterial phyla that currently suffer from high rediscovery rates. We thus provide a broad perspective on the great potential of *Myxococcota* as a source of chemically diverse starting points for drug discovery.

### Assignment of the new 154 strains overhauls the *Myxococcota* taxonomy

During our continuous efforts to explore the phylum *Myxococcota* from a taxonomic, biosynthetic, and chemical perspective, 154 novel strains were isolated and subjected to high-quality genome sequencing (Figure 1B, Supplementary Table 1 for assembly statistics and genome quality assessment). To provide an accurate taxonomic assignment, we retrieved a list of 109 *Myxococcota* type strains from the LPSN database and the corresponding genomes from public data sources^8^. A total of 37 type strains were sequenced as a subset of 154 strains in this study (data sources provided in Supplementary Table 2). The inter-species distance was measured using POCP (percentage of conserved proteins), whereby we identified POCP cutoffs of 76 % to assign strains to the same genus, and of 50 % to assign strains to the same family (Supplementary Figure 1 and 2), which deviates from the values commonly accepted in the literature (50 % for the genus level, and to our knowledge POCP has not previously been used for a family-level classification^38^). This choice of cutoffs is supported by a recent study that suggests a similar genus-level separation cutoff (75 %) within the family *Myxococcaceae*^39^.

While in most cases our POCP values agree well with established taxonomic ranks, we observed some taxonomic units where assignments need to be adjusted (Supplementary Figure 3). In particular, we propose merging *Myxococcaceae* and *Archangiaceae* into one family, since their inter-group mean POCP is above 50 % (Supplementary Figure 4). *Enhygromyxa, Plesiocystis*, and *Pseudenhygromyxa* belong to a new family separated from *Nannocystaceae* (Supplementary Figure 5). Further, we confirm that *Labilitrichaceae, Phaselicystaceae* (*Aetherobacter* and *Minicystis* are reclassified into this family), and *Polyangiaceae* are three separate families (Supplementary Figure 6), a matter that could not be resolved so far in the literature due to the small number of genomes available^40^. On the genus level, we propose splitting *Anaeromyxobacter, Haliangium, Myxococcus, Nannocystis, Pendulispora, Polyangium*, and *Sorangium* into multiple genera, respectively, including reclassifying *Polyangium aurulentum* and *Sorangium orientale* into a separate new genus each (Supplementary Figure 7-13). Additionally, we suggest changes in the genus assignment among some species of the genera *Archangium* and *Vitiosangium* and of the genera *Cystobacter* and *Melittangium* (Supplementary Figures 14 and 15), including reclassifying *Archangium disciforme* into *Angiococcus* as it has been originally suggested^41,42^. We further confirm the genus assignment within *Myxococcus, Pyxidicoccus*, and *Corallococcus*, which has recently been a matter of debate^9,42^. The full list of reclassification suggestions is provided as Supplementary Note 1.

Our set of 154 novel genomes revolutionizes the cultured taxonomic diversity of *Myxococcota*. Together with the reclassification suggestions from the type strains, we propose 18 new (and fusing two existing) families and 58 new genera, extending the total number of families and genera in *Myxococcota* to 28 and 90 (154.5 % and 181.3 % increase), respectively (Supplementary Table 3). Phylogenetically, our 154 new genomes are distributed throughout the whole *Myxococcota* phylum, with most new families situated within the suborders *Nannocystineae* and *Sorangiineae* (Figure 1B, Supplementary Table 4).

### *Myxococcota* taxonomy guides the identification of novel NPs

The presented taxonomic diversity is particularly intriguing, as surveys of the *Myxococcota* chemical diversity indicated that the NP chemistry is mostly conserved within closely related taxa^25^. A comprehensive analysis of the distribution of orthologous genes among *Myxococcota* genomes revealed that several clusters of orthologous genes (COG) categories were enriched among family-specific genes (Supplementary Figure 16). Notably, genes coding for the biosynthesis and transport of NP (COG category Q) were enriched among family-specific genes and >100-fold more likely family-specific than universally shared, suggesting that they often represent taxon-specific, specialized traits. Indeed, for selected characterized *Myxococcota* BGCs encoding for NPs with narrow-spectrum functions often desired for pharmaceutical applications (Figure 2A), we observe that they mostly also display a narrow distribution at the genus level and are often confined to specific families. Examples of such taxonomy-conserved NPs include antibiotics like cystobactamid, corallopyronin, and chlorotonil, while cytotoxic NPs such as epothilone, tubulysin, and myxothiazol display somewhat wider taxonomic distributions by trend^4^. Notable cases with widespread occurrence across taxonomic ranks are the myxochelin and DKxanthen BGCs, two classes known for their rather universal producer-specific functions in iron sequestration and sporulation, respectively^43,44^ (Figure 2A).

**Figure 2.**
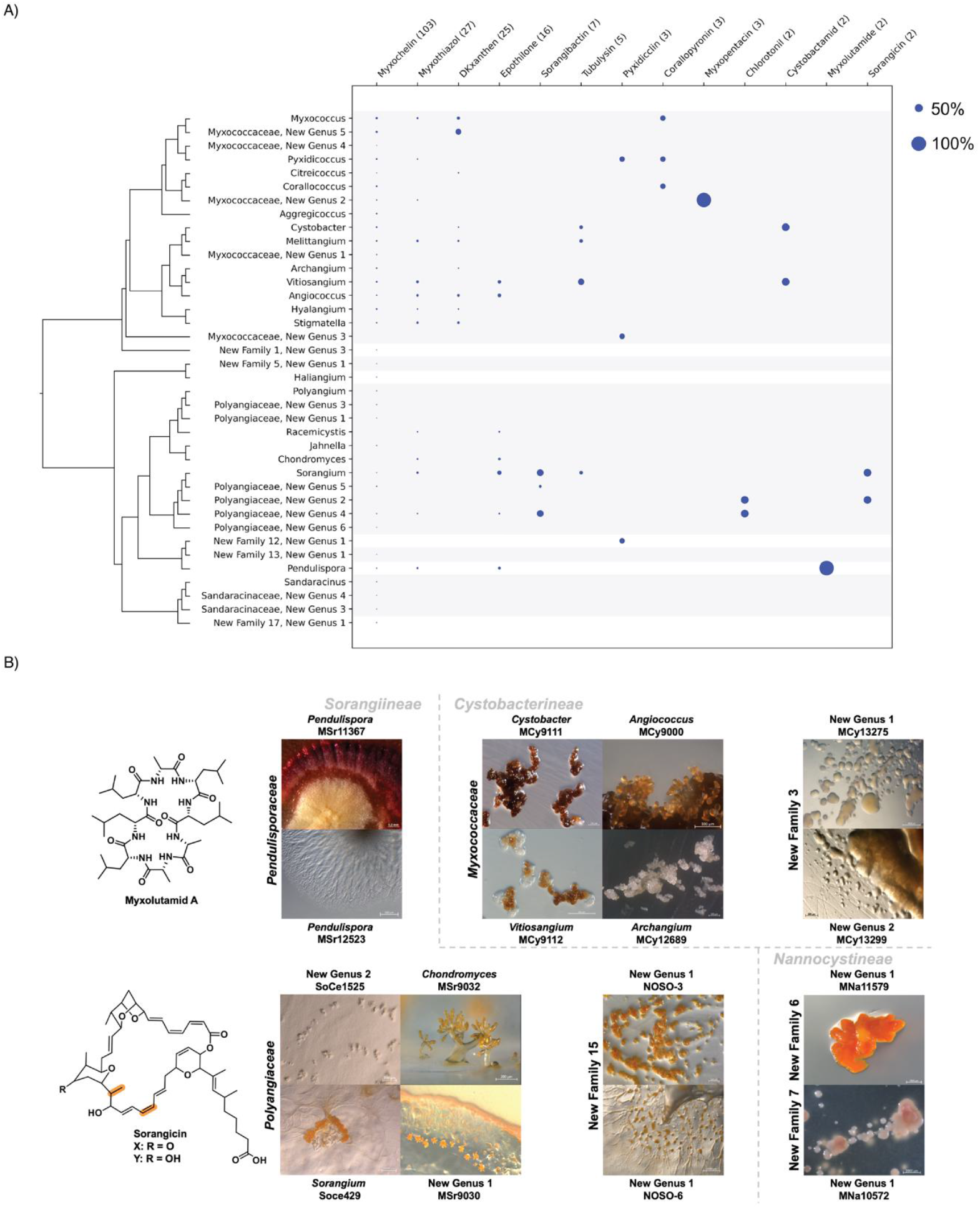
Taxonomic classification provides valuable metadata for strain prioritization in NP discovery. **(A)** Distribution of characterized *Myxococcota* BGCs across taxonomic groups. Dot size indicates the relative abundance of each BGC, with numerical values denoting absolute occurrences in previously reported *Myxococcota* genomes and in those presented in this study. The accompanying phylogenetic tree is resolved at the genus level. **(B)** Expression of characteristic morphological features—such as fruiting bodies, swarms, and fruiting body–like aggregates—is often conserved at the genus or family level within the *Myxococcota*. This morphological conservation frequently parallels chemical diversity, as illustrated by the myxolutamids, a genus-specific NP family conserved within *Pendulispora*, and the sorangicins, a family-specific NP family conserved within *Polyangiaceae*. Images were captured using a stereomicroscope; structural variations between new isolates and sorangicin A are highlighted in orange.

The observed trends corroborate our previous finding that increasing taxonomic distance is associated with greater chemical diversity at the genomic level. This underlines the value of exact taxonomic classification not only from a microbiological but also a chemical perspective^34^. We argue that this ‘taxonomy paradigm’ translates into increased chances for discovering structurally novel NPs from strains belonging to new genera and families, guided by the taxonomic distribution patterns of the matching BGCs. This concept was exemplified by the discovery of taxon-specific *Myxococcota* NPs from two distinct compound families, both structurally elucidated in this study: As a genus-specific example, we describe the myxolutamids —a new family of cyclic peptides isolated from *Pendulispora rubella* MSr11367 (structure elucidation and identification of the putative *mxI*BGC described in Supplementary Note 2). The isolated congener myxolutamid A (Figure 2B) features a unique all-D Leu-Leu-Leu-Ala-Ala-Leu-Leu-Ala sequence with the characteristic tandem mass spectrometry (MS^2^) fragmentation pattern of further family members identified with GNPS^45^ being consistently rich in Leu-Ala motifs (Supplementary Figures 17-26 and Table 5). In this study, we expand the family *Pendulisporaceae*^27^ by two additional members (Supplementary Table 4), enabling the proposal of a candidate myxolutamid BGC (Supplementary Figure 27, Table 6) showing conservation within *Pendulispora* (Figure 2A). As a family-specific example, we present two representatives of the sorangicin family—sorangicin X and Y (Figure 2B, structure elucidation described in Supplementary Note 3)—isolated from *Sorangium cellulosum* Soce429. Both previously unknown derivatives (Supplementary Figures 28-42 and Tables 7-10) have potent antibacterial activities similar to sorangicin A (Supplementary Table 12). Interestingly, some of their substitution patterns (highlighted in Figure 2B) match bioinformatics predictions for domain functionalities in the sorangicin BGC, where previous modifications observed in the sorangicin A structure could not be fully explained (Supplementary Figure 43, Table 11, BGC assignments described in Supplementary Note 4). Noteworthy, the sorangicin A producer *Sorangium cellulosum* SoCe1525 does not produce sorangicin X and Y or vice versa (Supplementary Figure 43). The sorangicin family shows conservation within the *Polyangiaceae* family (Supplementary Figure 44) with the sorangicin A producer SoCe1525 being reclassified from *Sorangium cellulosum* to *Polyangiaceae*, New Genus 2 (Figure 2B).

### Comparative genomics reveals untapped biosynthetic potential of new *Myxococcota*

To survey the biosynthetic potential comprised in our new genomes, BGCs predicted in our 153 *Myxococcota* genomes (one genome was excluded because of substandard assembly) were compared with BGCs in publicly available *Myxococcota & Actinomycetota* genomes (retrieved from the antiSMASH database^46^). In total, our dataset contains 4,780 BGCs, with an average of 31.2 BGCs per genome, compared to 27.1 BGCs in published *Myxococcota* genomes (inlet in Figure 3A). In contrast, although *Actinomycetota* harbor a much larger total number of BGCs due to extensive sampling (Figure 3A), they encode on average only 14.0 BGCs per genome (inlet in Figure 3A), which is considerably fewer than both our newly sequenced strains and previously published *Myxococcota* genomes. The higher average number of BGCs per genome in our dataset is likely a result of the characteristically larger genome sizes of *Myxococcota* strains (Figure 4A).

**Figure 3.**
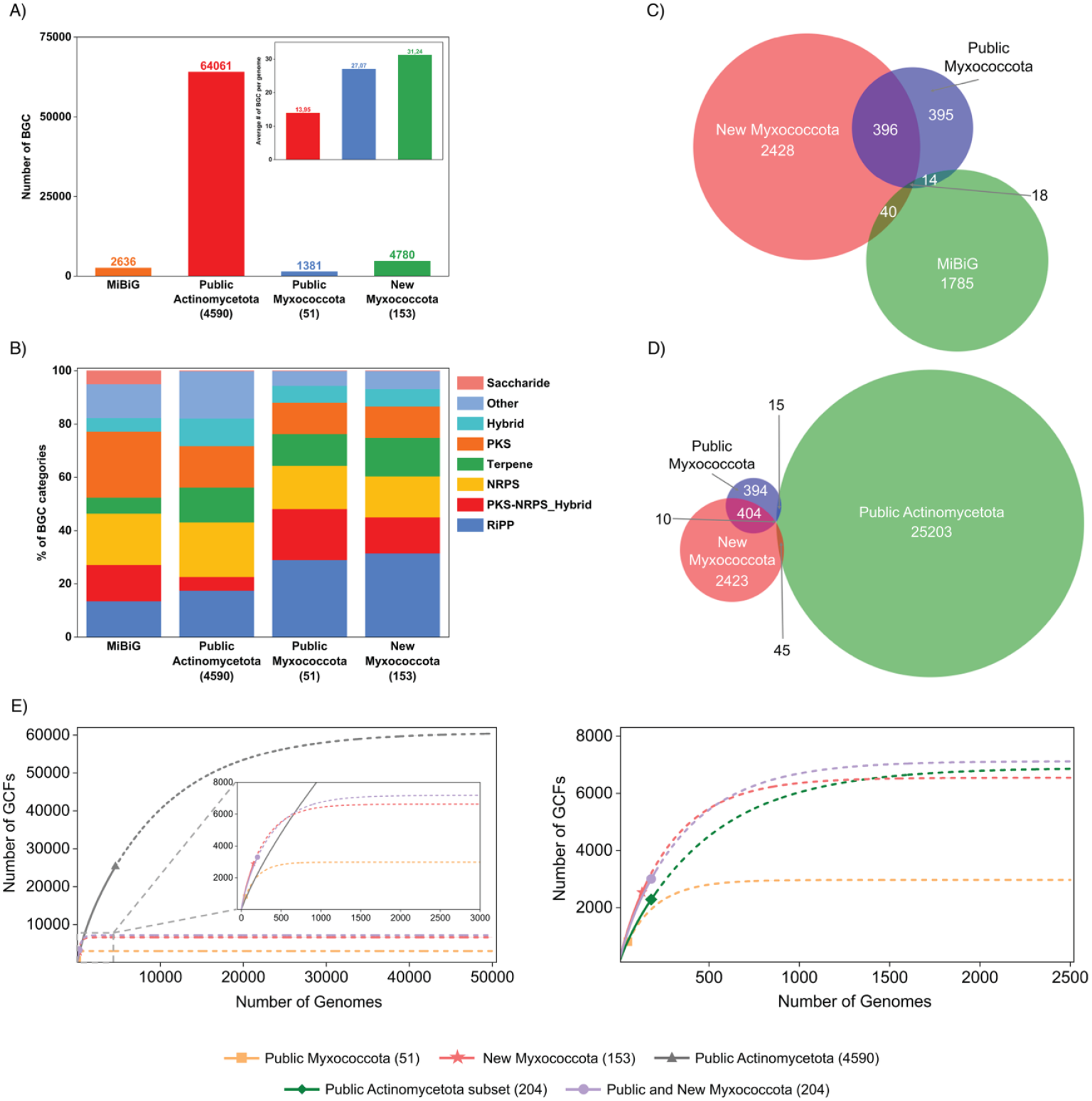
*Myxococcota* comprises substantial untapped biosynthetic potential. **(A)** Absolute numbers of BGCs detected in public *Actinomycetota* and *Myxococcota* genomes and the new dataset in comparison to experimentally characterized BGCs listed in MIBiG. Average number of BGCs per genome presented in inlet for reflecting oversampling in *Actinomycetota*. **(B)** Distribution of BGCs into major biosynthetic classes. Number of strains for each set in (A-B) given in parentheses; PKS = Polyketide Synthase, NRPS = Non-Ribosomal Peptide Synthethase, RiPPs = Ribosomally synthesized and Posttranslationally modified Peptide. **(C-D)** Venn diagram of unique and shared GCFs (identified using BiG-SLiCE, T = 0.4) of the *Myxococcota* genomes presented in this study with publicly available *Myxococcota* genomes and the MIBiG database (C), as well as *Actinomycetota* (D). **(E)** Rarefaction analysis of previously described public *Myxococcota* (orange), public *Actinomycetota* (black) & new *Myxococcota* genomes (red). All public *Actinomycetota* genomes are considered for the left graph, while for the right graph, we selected a subset of public *Actinomycetota* genomes (green) using 16S rRNA distance. Number of strains for each set given in parentheses. Solid lines in the curves represent observed values, while dotted lines represent extrapolated values. The inlet in left graph presents a magnified view of the bottom left axis of the left graph.

**Figure 4.**
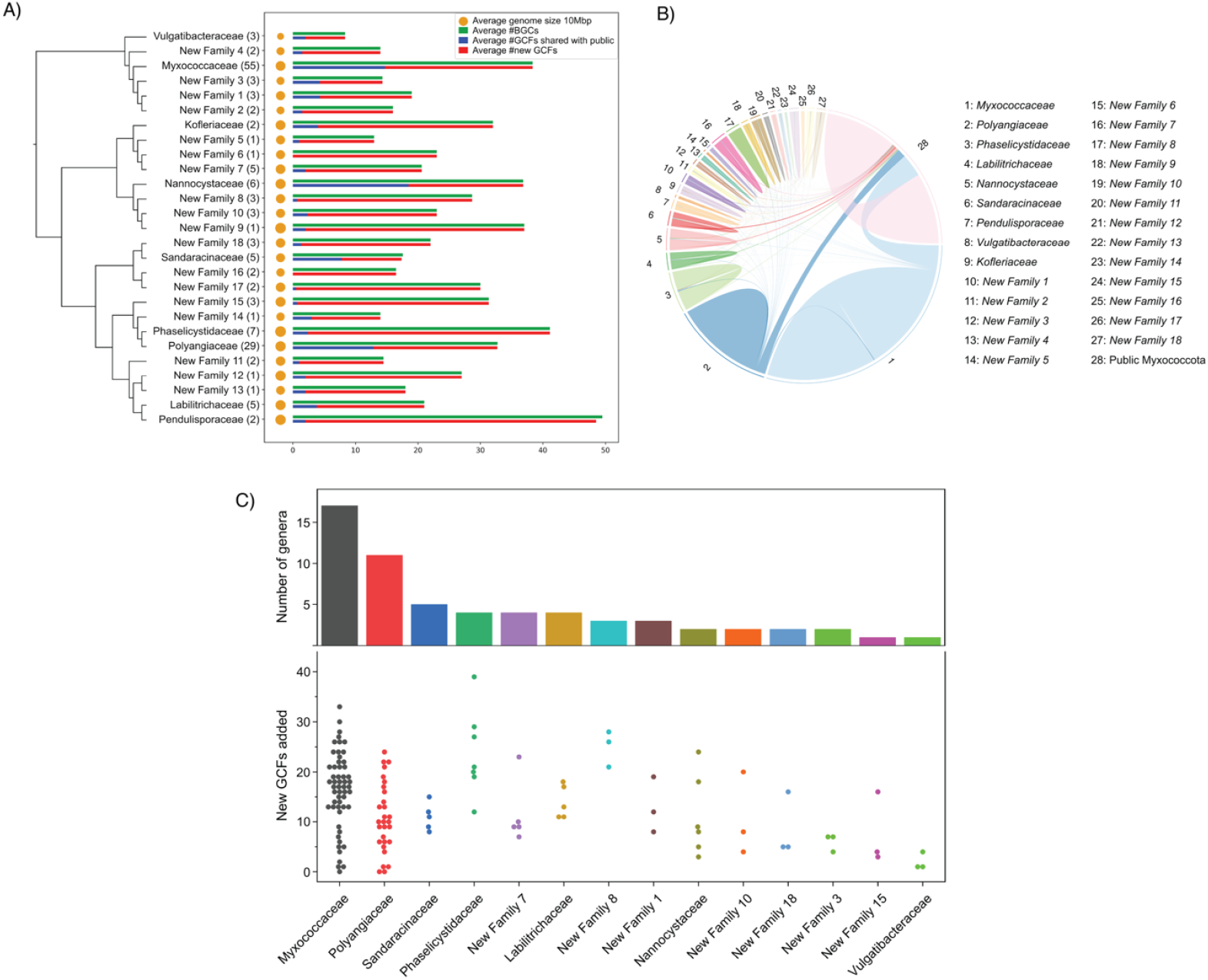
Family-level contributions and estimation of biosynthetic diversity within *Myxococcota*. **(A)** Family-level phylogenetic tree with accompanying barplots showing the numbers of BGCs and GCFs. Dots represent average genome size within the family. GCFs represented in public data are shown in blue, and GCFs only present in this dataset are shown in red. Number of strains for each family given in parentheses. **(B)** Ribbon plot showing the overlap of GCFs between different families in *Myxococcota* and public *Myxococcota* genomes. **(C)** Swarm plot showing the number of new GCFs added by individual strains in each family, with the number of genera present in each family. Only families with more than three strains are considered.

The distribution of biosynthetic classes reveals notable trends within the *Myxococcota* phylum, deviating from those observed for chemically characterized NPs. Among chemically characterized NPs, non-ribosomal peptides (NRPs) and polyketides (PKs) are reported as the dominant classes in *Myxococcota*^47^. However, our analysis with state-of-the-art tools indicates that the most abundant biosynthetic machinery detected in *Myxococcota* is the ribosomally synthesized and post-translationally modified peptides (RiPPs). In our new dataset, 31.6% of all BGCs correspond to RiPPs—an increase from the 29.0% observed in public *Myxococcota* genomes and notably higher than the 17.5% detected in *Actinomycetota*. Additionally, a substantial proportion of nonribosomal peptide synthetase–polyketide synthase (NRPS–PKS) hybrid BGCs were detected in both our new (13.5%) and publicly available (19.1%) *Myxococcota* genomes, markedly higher than the 5.1% observed in *Actinomycetota* (Figure 3B).

To assess their relatedness and diversity, BGCs in our dataset were compared with those from publicly available *Myxococcota & Actinomycetota* genomes and the MIBiG database^48^, a gold-standard database for experimentally characterized microbial BGCs. All collected BGCs were clustered into gene cluster families (GCFs) following standard procedures (selection of the clustering thresholds described in Supplementary Note 5, Supplementary Table 13, 14). Previously available *Myxococcota* BGCs cluster into 823 GCFs (∼1.7 BGC per GCF), of which 414 (∼50.3 %) are shared with GCFs from the new genome dataset. In total, our new set of *Myxococcota* genomes contains 2,882 GCFs (∼1.7 BGC per GCF), of which ∼83% are unique and not found in any other publicly available strains from *Myxococcota, Actinomycetota* or MIBiG (Figure 3C, D, Supplementary Figure 45), demonstrating the untapped biosynthetic potential of *Myxococcota*, to which our new set of 153 genomes substantially contributes. Importantly, there are very few shared GCFs between *Myxococcota* and *Actinomycetota* (Figure 3D), emphasizing the promise of *Myxococcota* as a source of chemically distinct NPs.

Clustering these BGCs into GCFs further underscores the disparity in sampling: *Actinomycetota* average 2.5 BGCs per GCF, whereas *Myxococcota* exhibit 1.7 BGCs per GCF, underpinning that the *Actinomycetota* phylum is much better sampled. Nevertheless, such comparisons are inherently limited by sampling depth, as even extensive genome sequencing captures only a fraction of the total diversity within a bacterial clade. To address this limitation, rarefaction analyses have been introduced in ecology^49,50^ to computationally estimate the total diversity from a given sample. Here, we leveraged it to assess the contribution of our dataset to the total biosynthetic potential of *Myxococcota* and *Actinomycetota*^51,52^ (Figure 3E), corroborating the well-accepted notion that *Actinomycetota* are a prime source of NPs (Figure 3E, left plot).

However, we argue that this comparison is heavily biased, since there are approximately 23-fold more strains from *Actinomycetota* available covering a much broader taxonomic space. To enable a more unbiased comparison, we selected a subset of 204 *Actinomycetota* strains (equal to the total number of publicly available and new *Myxococcota* strains) based on 16S rRNA sequence distance, to match the phylogenetic diversity to the set of *Myxococcota* strains (selection of Actinomycetota strains described in Supplementary Note 6). This reveals a revised picture: our new dataset exhibits a considerable biosynthetic potential approaching to that of *Actinomycetota* for a smaller number of strains, and the combined biosynthetic potential of public and new *Myxococcota* strains is estimated even slightly higher than that of *Actinomycetota* for a comparable number of strains (Figure 3E, right plot, selection of *Actinomycetota* strains described in Supplementary Note 6).

### *Myxococcota* families unevenly contribute to the detected biosynthetic potential

To explore the distribution of this untapped biosynthetic potential across the *Myxococcota* phylum, we conducted a comprehensive analysis to see which families among our new 153 *Myxococcota* strains contribute most in terms of enlarging the genetically encoded diversity of biosynthetic pathways. *Pendulisporaceae*^27^, *Phaselicystidaceae, Myxococcaceae* (which are now merged with *Archangiaceae*), New Family 9, and *Nannocystaceae* not only show the largest average number of BGCs and GCFs (>35 BGCs & GCFs in each family), but they also carry the largest genomes (>12.3 Mbp, Figure 4A). However, genome size does not necessarily correlate with the number of BGCs/GCFs, as exemplified by *Labilitrichaceae*, which have larger genomes (12.45 Mbp) but proportionally fewer BGCs/GCFs (21 average BGCs & GCFs, Figure 4A). Importantly, different families of *Myxococcota* share relatively few GCFs among each other. Nevertheless, novel strains from the previously known families like *Myxococcaceae, Polyangiaceae, Nannocystaceae*, and *Sandaracinaceae* share a non-negligible proportion of GCFs with previously described *Myxococcota* genomes. In contrast, the new families introduced in this study share relatively very few GCFs with previously published genomes (Figure 4B). As similar results were obtained for different *Myxococcota* genera (Supplementary Figure 46), it can be concluded that the new genera and families bear the most promise for the identification of new and chemically diverse NPs.

To guide targeted strain isolation for NP discovery, we asked which *Myxococcota* families or genera are most likely to expand biosynthetic diversity with additional sampling. We quantified the number of unique GCFs contributed by each strain within families or genera represented by at least three isolates (Figure 4C and Supplementary Figure 47). The number of GCFs added per strain varied substantially across families, indicating pronounced differences in the biosynthetic potential even among closely related strains. Notably, New Family 8 and *Phaselicystidaceae* consistently contribute high levels of novel biosynthetic diversity, with each strain harboring at least 12 and up to 39 unique GCFs. However, both families are relatively small, comprising only three and seven strains, respectively. Their high diversity may stem from the inclusion of phylogenetically diverse genera: *Phaselicystidaceae* has seven strains spanning four genera, while New Family 8 includes three strains from three distinct genera. However, we also observe contrasting examples: *Sandaracinaceae*, though composed of five strains from five different genera, shows lower biosynthetic potential, with all strains encoding 15 or fewer unique GCFs. The larger families, such as *Myxococcaceae* and *Polyangiaceae*, have a very large variance, comprising strains both poor and rich in unique GCFs. It should be emphasized that this representation of the genetic potential does not account for GCFs that can currently not be annotated using the available *in silico* tools.

### ABC-Myxo facilitates motif-guided NP discovery

To promote exploration of our new genomes and their biosynthetic potential, we provide an interactive web-based resource, ABC-Myxo (Atlas of Biosynthetic Gene Clusters in *Myxococcota*, https://tools.helmholtz-hips.de/abc_myxo/). Its functionality is based on our previous resource ABC-HuMi^53^ and encompasses 7,577 BGCs identified from the genomes of both publicly available and new *Myxococcota* strains, as well as from the MIBiG database^48^, together with their corresponding clustering in GCFs. The ABC-Myxo database provides browse and search functionality, as well as visualizations of GCF similarity networks. The information about BGCs and GCFs is stored in interactive tables. The BGCs are complemented with antiSMASH-predicted properties and linked to the complete antiSMASH reports. The database incorporates BLAST+^54^ for sequence similarity search and cblaster^55^ for identifying homologs of user-provided BGCs in the database.

To exemplify a motif-guided genome-mining workflow for our *Myxococcota* genomes, we used cblaster on our genome set to search for BGCs comprising genes encoding the seven-enzyme cassette responsible for cispentacin (amino-cyclopentyl-carboxylic acid) formation, resembling a recent discovery of rare building block biosynthesis not known from *Myxococcota* and derived from the nucleoside antibiotic amipurimycin^56,57^ (Figure 5). Mapping the distribution of similar operons across our extended genome collection revealed ten occurrences encoding for a cispentacin-cassette comprising BGC, nine of which are near NRPS-encoding genes (Figure 5), with all strains belonging to the family of *Myxoccocaceae*. Targeted inactivation of the cispentacin-cassette containing BGC in *Myxoccocaceae* New Genus 2 MCy9003 (*mxp* BGC) strain followed by LC-MS profiling of mutant extracts (Supplementary Figure 48), validated its involvement in the biosynthesis of the novel class of NRPS-PKS hybrids termed myxopentacins (structure elucidation described in Supplementary Note 7, Supplementary Figures 49-57 and Table 15). Interestingly, the isolated congener myxopentacin A was found to incorporate not only one cispentacin building block, but five units (Figure 5). Notably, we find three strains harboring an *mxp* BGC in our genome set, with all of them belonging to *Myxococcaceae* New Genus 2 (Figure 2A). While the *mxpB-H* genes show homology to *amcB-F* of the amipurimycin BGC, enabling the generation of the cispentacin building block, the adjacent NRPS domains encoded downstream to the cispentacin locus are plausibly involved in the peptide chain assembly (Figure 5, description of the *mxpBGC* in Supplementary Note 8, Supplementary Figure 58, Table 16). Feeding experiments with isotopically labelled precursors confirm aminobutyric-acid, acetate and arginine to be incorporated in the structure of the hybrid NRP-PK natural product (Figure 5). This is the first report of any NP from myxobacteria containing the respective unusual moiety as predicted by our phylogeny-guided genome mining efforts. The details of multiple cispentacin incorporation, as well as potential bioactivity and functional significance of mature myxopentacins for the producing strain, are currently the subject of additional investigations.

**Figure 5.**
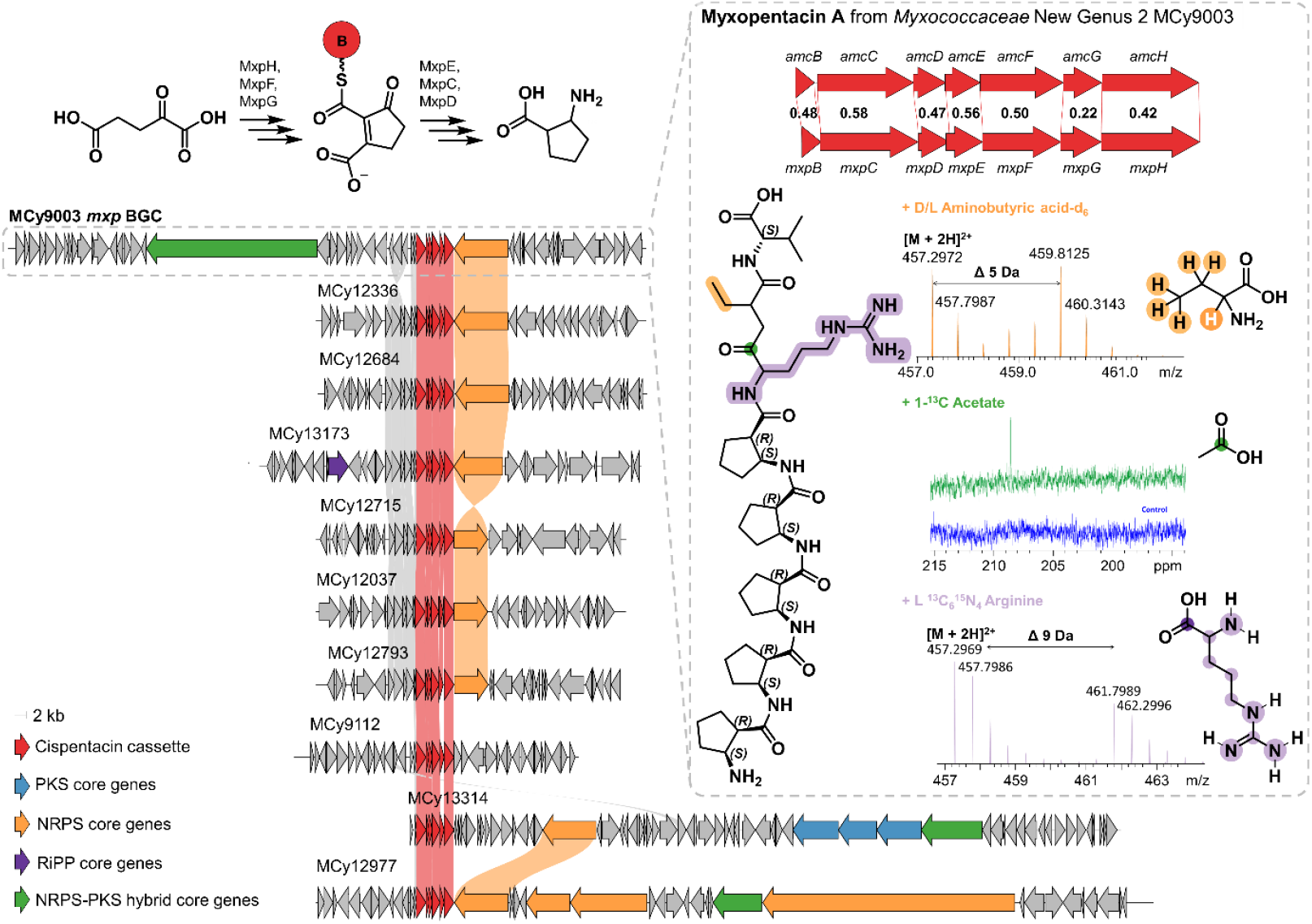
Genome mining unveils the family *Myxococcaceae* to encode for distinct cispentacin-containing NPs. A web-based search in ABC-Myxo identifies ten BGCs with a cispentacin cassette, of which one was assigned to their products – the myxopentacins. Cblaster-based search for the *amcB-H* operon (red) required for cispentacin formation yielded BGC hits from ten *Myxococcota* strains, all belonging to the family *Myxococcaceae*. Besides incorporation of valine and five cispentacin building blocks, feeding experiments in MCy9003 confirmed incorporation of aminobutyric acid (orange), acetate (green) and L-arginine (purple) as a hybrid NRP-PK building block.

## DISCUSSION

Recently, large-scale genomic and metagenomic analysis across diverse bacterial phyla^58^ and environments^59–61^ have attempted to uncover the biosynthetic potential of particular taxa, ecological niches, or microbiomes. In these analyses, *Myxococcota* were identified as having relatively high biosynthetic potential, although they were phylogenetically not broadly represented. In this study, we isolate and culture 154 new *Myxococcota* strains and establish high-quality genomes, quadrupling the number of available full *Myxococcota* genomes, greatly expanding the available cultured diversity and enabling a refined taxonomic framework for the phylum, which is extended by 58 genera and 18 families. The substantial inconsistencies observed with existing classifications highlight the need for clade-specific metrics; accordingly, we propose POCP thresholds of 50% for families and 76% for genera. This emphasizes that taxonomic boundaries cannot be universally defined across bacteria.

The expanded genomic dataset reveals a strong correlation between taxonomic distance and biosynthetic diversity. Most GCFs are family-specific, and only a single GCF spans all three suborders (Supplementary Figure 60), underscoring the importance of focusing on underexplored lineages to maximize chances for chemical novelty. This genomic view provides a more balanced picture than metabolomics-based surveys^34^, which are biased toward strains that grow or produce NPs readily under laboratory conditions.

Our experimental results demonstrate that both rare and well-studied lineages harbor substantial biosynthetic richness. We report three NP families—myxolutamids, sorangicin X/Y, and myxopentacins, two of which present new chemical scaffolds. *Phaselicystidaceae* emerges as an exceptionally productive lineage, whereas genome mining within *Myxococcaceae* exposes untapped biosynthetic potential even in densely sampled families. These findings validate the combined use of precise taxonomy and targeted genome mining for NP discovery.

To support systematic exploration, we developed ABC-Myxo as an integrated platform for navigating *Myxococcota* genomes and BGC diversity. The resource facilitates dereplication, motif-guided searches, and large-scale survey analyses, accelerating discovery across both familiar and newly defined taxa.

Finally, when accounting for phylogenetic diversity rather than sheer strain counts, *Myxococcota* exhibit genome-encoded biosynthetic potential comparable to *Actinomycetota*. However, the near-linear rarefaction curves of both phyla indicate that neither of them is close to saturation. Given the high re-discovery rates in classical natural product-producing taxa, we argue that *Myxococcota* are an emerging source of novel compounds. Continued sampling of underexplored taxa remains essential, and the genomic framework presented here provides a guide for prioritizing *Myxococcota* lineages with the greatest promise for not only medically relevant NPs.

## METHODS

### Strain cultivation, genomic DNA isolation, and whole-genome sequencing

Myxobacterial strains were isolated using baiting techniques with *Escherichia coli* and filter paper on Mineral Salt MS21 medium^35^. Samples from the marine environment were also isolated using baiting techniques on commercial sea-salt agar with pH and salinity adjusted accordingly. Isolation of myxobacteria was based on observable swarming, fruiting bodies, and fruiting body-like aggregates on the medium and sample inoculum. Purification was performed by cutting the outermost swarm edge or by washing the fruiting bodies in sterile water or sea salt solution. Axenic cultures were maintained on standard buffered VY/2 agar^35^ or on the corresponding buffered VY/2/SWS agar^36^. Unless specified, the *Myxococcota* strains were grown in a 300 mL triple-vent baffled-flask containing 5 0mL medium and cultivated for 3-5 days under dark environment, 26% humidity, 30°C, and 180 rpm. shaking conditions. The strains were grown in suitable media including modified Amb (Soluble starch 5 g/L, Bacto Casitone 2.5 g/L, MgSO_4_ x 7 H_2_O 0.5 g/L, K_2_HPO_4_ 0.25 g/L, HEPES 10mM, pH adjusted to 7.0 with KOH before autoclaving), CYH (Soy meal (Hensel) 2 g/L, Bacto Casitone 3 g/L, Glucose 2 g/L, Soluble starch (Roth) 8 g/L, Bacto Yeast extract 1.5 g/L, CaCl_2_ x 2 H_2_O 1 g/L, MgSO_4_ x 7 H_2_O 1 g/L, HEPES 50mM, Fe-EDTA 8 mg/L, pH adjusted to 7.2 with 10N KOH), and buffered VY/2 broth^35^. Strains MCy9330 and MSr10579 were cultured in MD1G-5gC (Bacto Casitone 5 g/L, CaCl_2_ x 2 H_2_O 0.5 g/L, MgSO_4_ x 7 H_2_O 2 g/L, Glucose 3.5 g/L, pH adjusted to 7.0 with KOH before autoclaving) at 26°C. Cultivation of strains DSM 15217^T^, DSM 21377^T^, MNa10571, MNa10576, MNa12472, MSr9506, MSr10681, MSr13296, MSr13327 included seawater salt solutions in the medium as described in previous studies^37,62–64^.

Huge cell aggregates were broken into finer flakes and tiny aggregates using a sterile glass tissue grinder, while cells growing as a homogeneous suspension were directly used for genomic DNA isolation. The extraction of a high-quality genomic DNA was prepared using a Qiagen DNeasy UltraClean® Microbial and a Monarch® Nucleic Acid Purification kit (New England BioLabs® Inc.) following the manufacturer’s instructions. The cells were lysed with a minimal bead-beating condition (4.0 m/sec, 5 sec) using an MP Biomedicals Fastprep-24TM 5G bead-beating machine. Quality check of the genomic DNA was performed by measurement of a 1 µL aliquot using a Nanodrop OneC (Thermo Scientific) and gel electrophoresis (Bio-Rad) using the following set-up condition: Biozyme agarose 8 g/L, Tris-acetate-EDTA buffer 1 x, 60V, 1000mA, 150W, 1 h run time.

For the generation of barcoded libraries for DNA sequencing, an aliquot of 1 µg genomic DNA was sheared, end-repaired, and ligated to barcoded adapters using the SMRTbell® Express Template Prep Kit 2.0 (Pacific Biosciences). Multiplexed microbial libraries (10-kb SMRTbell) were prepared according to the instructions in the Pacific Biosciences manual. Size selection was performed using AMPure Beads, as instructed in the PacBio manual. The libraries were pooled and then sequenced on a Sequel II machine (Pacific Biosciences). The PacBio sequencing was performed by the NGS Competence Center Tübingen (NCCT), Germany, in collaboration with Universitätsklinikum, Tübingen, Germany.

### Genome assembly

The genome assembly for 154 strains was done using three different pipelines, depending on the type of sequencing reads, Pacbio CLR or Pacbio HiFi reads. For the CLR reads, first, the assembly was performed using Flye^65^ (v2.9.2) with default parameters. If the assembly using default parameters was not satisfactory, it was repeated by changing the min-overlap (-m) parameter in the range {1000, 2000, 3000, …, 10000} and the best assembly was chosen among all based on the size and the coverage of the longest contig. For HiFi reads, the assembly was performed using three different assemblers (Flye (v2.9.2), HiFiasm^66^ (v0.19.8), and IPA^67^ (v1.8.0) (all with default parameters)), and the best assembly was chosen among the three different assemblies similarly. After performing the assembly, contamination and completeness were checked using CheckM2^68^ (v1.1.0). Assembly statistics are given in Supplementary Table 1.

### BGC identification & clustering

antiSMASH^69^ (v7.1.0) was used to identify biosynthetic gene clusters (BGCs) in the new *Myxococcota* strains. Further, BGCs were also collected from the antiSMASH database^46^ (v4.0) belonging to *Myxococcota* and *Actinomycetota* phylum, as well as from MIBiG^48^ (v3.0) database. All these BGCs were clustered using BiG-SLiCE^70^ (v2.0.0, default parameters) to identify the GCFs. We have compared clustering produced with BiG-SLiCE^70^ and BiG-SCAPE^71^ with different threshold values, see Supplementary Note 5.

### Rarefaction analysis

The rarefaction and extrapolation analyses were done using the iNEXT^72^ R package. In the rarefaction analysis, the number of GCFs in a series of subsamples of available strains is calculated, and these numbers are used in the extrapolation analysis to predict the projected number of GCFs if a higher number of strains were available. Two different types of rarefaction analysis were done in the study: First, using all the public *Actinomycetota* strains, and second, using a subset of public *Actinomycetota* strains that have the same phylogenetic diversity as that of *Myxococcota* (public and new) strains established using the 16S identity (see Supplementary Note 6). For both types of rarefaction analysis, BGCs from the respective *Actinomycetota* strains were mixed with the BGCs from *Myxococcota* (public and from the new strains), as well as MIBiG, and clustered using BiG-SLiCE to identify GCFs. Next, for each of the identified GCFs, the number of strains from individual data sources was extracted and used as ‘incidence-freq’ data for the iNEXT main function. For the rarefaction analysis using all *Actinomycetota* strains, the number of knots was set to 100,000, and the endpoint was set to 50,000 strains, while for the other analysis, the number of knots was set to 5,000, and the endpoint was set to 2,500 strains. By default, the number of bootstrap replications was 50 for both rarefaction analyses.

### Type strain dataset

We extracted the list of 109 type strains from LPSN^8^ which represents all the valid *Myxococcota* species (except for *Melittangium alboraceum* as there is no cultured type strain available at the moment) known from the phylum *Myxococcota* as of June 2024. This list also incorporates type strains from species that are currently not included, or not yet validly published from LPSN, but have been validly proposed in the literature^10,11,73,74^. To prepare the genome dataset, we downloaded 72 publicly available genomes, 71 from NCBI genomes and 1 from SRA^75^. For the rest of the type strains whose reference genomes are not publicly available, 37 species are proposed from the set of 154 new strains. The genomes of type strains *Hyalangium minutum* DSM 14724^T^ (NCBI RefSeq assembly: GCF_000737315.1) and *Byssovorax cruenta* By c2 (Submitted GenBank assembly: GCA_001312805.1) are publicly available. However, both assemblies are of low quality. Accordingly, we included their improved genomes in our list of 37 genomes. The full list of used type strains is provided in the Supplementary Table 2.

### POCP cutoffs identification

We performed the pairwise calculation of POCP using the nextflow pipeline POCP-nf^76^ (v2.2.0) for the genomes of all the *Myxococcota* type strains collected. The output species-level matrix was converted into genus- and family-level matrices by averaging the corresponding pairwise values for the inter- and intra-genus/family POCP (the values on the diagonal were ignored as they represent self-comparison). We compared the inter-group averages against the intra-group averages to infer the POCP cutoffs for genus and family level.

### Phylogenetic reconstruction and visualization

Phylogenetic trees were constructed using the BIONJ algorithm through FastME (v2.1.6.3)^77,78^ given the distance matrix based on POCP as input. Visualization of the phylogenetic trees was generated by Geneious^79^ (v2024.0.5).

### Taxonomic re-classification

We evaluated the topologies of the phylogenetic tree of the type strains (with *Desulfovibrio desulfuricans* DSM_642, NCBI RefSeq assembly GCF_000420465.1 as the outgroup member), and proposed revised genus and family groupings of species by applying the corresponding POCP cutoffs for genus and family level described above. We made re-classification suggestions when discrepancies were observed between our inferred species groupings and the established genus and family-level classifications. The complete list of reclassification suggestions is provided in Supplementary Note 1.

### New taxa suggestion

We evaluated the topologies of the phylogenetic tree of the type strains together with the 154 new strains (with *Desulfovibrio desulfuricans* DSM_642, NCBI RefSeq assembly GCF_000420465.1 as the outgroup member) and applied the corresponding POCP cutoffs described above and grouped the strains that were found to reside within the genus and family cutoffs. The groups were merged until reaching the best concordance with the phylogenetic tree. We proposed that the groups of strains containing no representative type strains as new *Myxococcota* genera and families.

### UHPLC-MS measurements

High resolution measurements were acquired on a Bruker Daltonics maXis 4G UHR-TOF device (Bruker Daltonics, Billerica, MA, USA), which is an ultrahigh resolution time of flight mass spectrometer utilizing an ESI source and following MS settings: capillary voltage 4000 V, end plate off-set −500 V, nebulizer gas pressure 1 bar, dry gas flow rate 5 L/min, dry gas temperature 200 °C, mass scan range m/z 150–2500. Calibration was performed on the masses of the first isotope signal of sodium formate clusters in Quadratic + HPC mode. For MS/MS measurements a collision energy of 5 eV was used in CID fragmentation mode. LC-MS data were analyzed with DataAnalysis 4.4 (Bruker Daltonics, Billerica, MA, USA).

### NMR analysis

All NMR spectra were recorded on a Bruker Avance III (Ultrashield) 500 MHz spectrometer equipped with a 5 mm TCI cryoprobe and or a Bruker Avance III (Ascend) 700 MHz spectrometer. All measurements were carried out using standard pulse programs. Observed chemical shifts (δ) are specified in ppm and coupling constants (*J*) in Hz. Spectra were calibrated on the intrinsic chemical shifts of the remaining undeuterated solvent.

### Isolation of myxolutamid A

*Pendulispora* MSr11367 was fermented in 2-SWT medium (Soluble starch 2 g/L, Bacto Tryptone 3 g/L, Bacto Soytone 1 g/L, Glucose 2 g/L, Maltose monohydrate 1 g/L, Cellobiose 2 g/L, MgSO_4_ x 7 H_2_O 1 g/L, CaCl_2_ x 2 H_2_O 0.5 g/L, HEPES 2.38 g/L, pH adjusted to 7.0 with KOH before autoclaving), under supplementation of 2% XAD-16 adsorber resin and soil extract. Soil extract was obtained through lyophilization of a soil sample (pond in Schiffweiler, Germany) and super critical fluid extraction using a Waters MV-10 ASFE system in 25 mL extraction vessels. Approximately 36 mg of dried soil extract were dissolved in 4 mL of water/methanol 1:1, sterile filtrated and 1 mL thereof was added to 100 mL liquid culture. After 14 days of incubation at 30 °C, cells and resin were harvested and extracted with MeOH and acetone. The obtained extract was filtrated and dried before resuspension in MeOH followed by liquid-liquid partitioning between MeOH and hexanes. The MeOH phase was dried and redissolved in water and extracted three times with chloroform and ethyl acetate each. The obtained MeOH extract was fractionated using a gravitation column with 3.5 cm diameter and 104 cm length (packing height around 75 cm) filled with Sephadex LH 20. MeOH was used as mobile phase and the flow rate adapted to approximately 246 µL/min assuming a volume per drop of 61.5 µL. Elution was followed by LC-MS and myxolutamid containing fractions subjected to a final separation step using semipreparative HPLC. Separation was achieved on a Waters XBridge Peptide BEH C18 OBD prep column (250 x 10 mm, 5 µm) applying a gradient from 50 to 95% acetonitrile in water with 0.1 % formic acid for 29.5 min at a flow rate of 5 mL/min. For each injection the column was equilibrated for 5 min at 5% and flushed with 95% acetonitrile in water with 0.1 % formic acid for 2 min at 45 °C. Single ion monitoring was utilized for detection of the target mass at m/z 779.5 with 10 V CID voltage on a Thermo Scientific ISQ EM system.

### Marfey’s analysis

0.1 mg of myxolutamid A was dissolved in 100 µL of 3 M HCl for 45 °C at 110 °C. The dry residue was split in two 50 µL aliquots in *dd*H_2_O. To both samples 60 µL of 1 M NaHCO_3_ as well as 20 µL of the derivatization agent (1 % 1-fluroro-2,4-dinitrophenyl-5-leucine-amide solution in acetone [D-FDLA and L-FDLA]) were added and incubation was carried out at 40 °C and 700 rpm for 2 h. The reaction was quenched by addition of 60 µL 3 M HCl and diluted with 150 µL of both MeOH and acetonitrile to be measured on the maXis 4G UHR-TOF (Bruker Daltonics, Billerica, MA, USA) coupled to a Dionex Ultimate 3000 SL system (Thermo Fisher Scientific, Waltham, MA, USA). Configuration of amino acids was evaluated via Marfey’s assay after hydrolysis of the amid bonds, derivatization of the free amino acids and chromatographic comparison to derivatized reference amino acids. Samples were analyzed on the maXis 4G UHR-TOF (Bruker Daltonics, Billerica, MA, USA) coupled to a Dionex Ultimate 3000 SL system (Thermo Fisher Scientific, Waltham, MA, USA). Analytes were separated on a Waters Acquity BEH C18 column (50×2.1 mm, 1.7 μm) using the following multi-step gradient with A=0.1% formic acid in ddH2O and B=0.1% formic acid in acetonitrile: 5-10% B in 1 min, 10-35% B in 14 min, 35-55% in 7 min, 55-80% in 3 min, hold at 80% for 1 min and reequilibration at 5% B for 5 min. UV spectra were acquired at 340 nm together with MS detection in centroid mode from 150 to 1000 m/z using a standard ESI source.

### Isolation of sorangicin X and Y

*Sorangium* So ce429 was fermented in CyH medium (Soy meal (Hensel) 2 g/L, Bacto Casitone 3 g/L, Glucose 2 g/L, Soluble starch (Roth) 8 g/L, Bacto Yeast extract 1.5 g/L, CaCl_2_ x 2 H_2_O 1 g/L, MgSO_4_ x 7 H_2_O 1 g/L, HEPES 50mM, Fe-EDTA 8 mg/L, pH adjusted to 7.2 with 10N KOH) under supplementation of 4% XAD-16 adsorber resin. After 23 days of incubation at 30 °C, 160 rpm, freeze-dried cells and resin were extracted with 1.25 L MeOH at a flowrate of 5 mL/min in an empty flash chromatography column casing. The dried extract was partitioned between n-hexane (3 x 500 mL) and MeOH /H_2_O (500 mL, 9:1). Afterwards, the MeOH/H_2_O phase was dried, dissolved in H_2_O (300 mL), and extracted with CHCl_3_ (3 x 500 mL). The CHCl_3_ phase containing sorangicins X and Y was dried and dissolved in MeOH before separation on a Pure C-815 normal phase flash chromatography flash system (Büchi). A 50 g Sfär Silica D column (Biotage) was used as the stationary phase, and hexane (solvent A), DCM (solvent B), EtOAc (solvent C), and MeOH (solvent D) were used as mobile phases. The flow rate was set to 45 mL/min with a multistep gradient for elution: (1) linear gradient from 100% A to B over 12 CV (column volume; 1 CV = 45 mL), followed by 4 CV 100% B; (2) 20 CV linear gradient from 100% B to C, followed by 5 CV of C; and (3) 20 CV linear gradient from 100% C to 50% D and 5 CV linear gradient from 50% to 100% D. Fractions were analyzed by HPLC-MS and further purified on a Waters Autopurifier high-pressure gradient system. Separation was carried out on a Waters X-Bridge prep C-18 5 µm OBD, 150 × 19 mm column using H_2_O + 0.1% FA (A) and ACN + 0.1% FA (B) as mobile phase at a flow rate of 25 mL/min. Separation was started with 5% B for 1 min, followed by a ramp to 50% B over 4 min and an increase to 70% B over 17 min. The column was then flushed with a ramp to 95% B over 1 min and at 95% B for 3 min, brought back to 5% B within 2 min, and re-equilibrated at 5% for 2 min.

### Isolation of myxopentacin A

MCy9003 was fermented in RG5 medium (Soya peptone (Roth) 0.5g/L, Soytone (BD) 0.5g/L, Soya meal (Hansel) 0.2g/L, Corn steep solids (Sigma) 1g/L, Yeast extract (BD) 0.5g/L, Soluble starch (Roth) 8g/L, Baker’s yeast 5g/L, Gluten from wheat (Sigma) 2g/L, MgSO_4_ 7H_2_O 1g/L, CaCl_2_ 2H_2_O 1g/L, HEPES 5.95g/L, pH adjusted to 7.2 with KOH) under supplementation of 2% XAD-16 adsorber resin. After 12 days of incubation at 30 °C, 180 rpm, cells and resin were extracted four times with 100 mL acetone. The dried extract partitioned between n-hexane (2 x 100 mL) and MeOHl/H_2_O (100 mL, 9:1). Afterwards, the MeOH/H_2_O phase was dried, dissolved in H_2_O (300 mL), and extracted with CHCl_3_ (3 x 500 mL). The myxopentacin A containing MeOH-layer was fractionated on Sephadex LH-20 resin using MeOH as eluent. A flow rate of 30 drops/ min was applied and 600 drops were collected for each fraction. An aliquot of every fraction was taken and measured using LC-MS to monitor elution. Further separation was achieved on a Dionex Ultimate 3000 coupled with a Bruker High-Capacity Trap mass spectrometer (HCT); Column: XSelect CSH (PFP) 250 x 10 mm, 4 μm; flowrate 4 ml/min under HILIC conditions and column temperature 40 °C with H_2_O as eluent A, ACN as eluent B and 100 mM Ammonium formiate (AmFo) in dH_2_O as eluent C. The following gradient was applied: 0-1 min 97 % eluent B; 1-21 min linear decrease of eluent B to 47 %; 21-25 min eluent B at 47 %, 25-26 min linear increase of eluent B to 97 %, 26-30 min re-equilibration with 97 % eluent B. Eluent C was constantly at 3% over the whole gradient. Final separation was achieved using a Jupiter Proteo 250 x 10 mm, 4 μm column; flowrate 5 ml/min and column temperature 40 °C with H_2_O as eluent A, ACN as eluent B and 100 mM AmFo in dH2O as eluent C. The gradient, 0-1 min linear increase of eluent B from 5 % to 15 % and eluent constant at 3 %; 1-21 min linear increase of eluent B from 15 % to 25 % and eluent C constant at 3 %; 21-22 min linear increase of eluent B from 25 % to 95 % and eluent C constant at 3 %; 22-27 min 95 % eluent B and 3 % eluent C; 27-28 min linear decrease of eluent B from 95 % to 5 % and eluent C constant at 3 %; 28-31 min re-equilibration with 5 % eluent B and 3 % eluent C, was applied.

### Stable isotope-feeding

To test substrate incorporation, 10 µL of the labelled substrate-solutions (L-valine *d*_*8*_ 0.25 M in MeOH/H_2_O= 1:1, ^13^C_2_-acetate 0.5 M, 1-^13^C-acetate 0.5, 2-^13^C-acetate, L-arginine^15^N_4_^13^C_6_ 0.25 M, L/D-aminobutyric acid *d*_*6*_ 0.25 M) were added to a 10 mL culture in 50 mL flasks each, twice over two days. 24 h after the last addition 400 µl of XAD-16 suspension were added and cultures were harvested after 9 days total fermentation time. The supernatant was discarded and the cells and the XAD were frozen and lyophilized before extraction as described for the isolation. The dried extract was dissolved in 1 mL MeOH. To purify 1-^13^C-acetate labelled myxopentacin A for NMR confirmation, a culture of MCy9003/ *pSBtn5Kan001nrps12* mutant in CyS media with 75 µg/ml kanamycin was used. For four days, 500 µl of a 0.5 M 1-^13^C acetate solution were added. Four days after inoculation, 2 mL of a XAD-16 suspension were added to the cultures with subsequent processing as described in isolation of myxopentacin A.

### Construction of plasmids for mxp inactivation mutants

Routine handling of nucleic acids, such as isolation of plasmid DNA, restriction endonuclease digestions, DNA ligations, and other DNA manipulations, was performed according to standard protocols^80^. *E. coli* DH10β (Invitrogen) was used as the host for standard cloning procedures. *E. coli* strains were cultured in 2TY medium or on 2TY agar (1.6 % tryptone, 1 % yeast extract, 0.5 % NaCl, (1.8 % agar), dH*2*O) at 37 °C, at 200 rpm overnight. Transformation of *E. coli* strains was achieved via electroporation in 0.1 cm-wide cuvettes at 1350 V, 200 Ω, 25 µF. Antibiotics were used at the following final concentrations: 100 μg mL-1 ampicillin, 50 μg mL-1 kanamycin. Plasmid DNA was purified by standard alkaline lysis or by using the GeneJet Plasmid Miniprep Kit (Thermo Fisher Scientific). Restriction endonucleases, alkaline phosphatase (FastAP) and T4 DNA ligase were obtained from Thermo Fisher Scientific. Oligonucleotides used for PCR and sequencing were obtained from Sigma-Aldrich. PCRs were carried out in Mastercycler® pro (Eppendorf) using Phusion™ High-Fidelity or Taq DNA polymerase (Thermo Fisher Scientific) according to the manufacturer’s protocol. For Phusion™: Initial denaturation (2 min, 98 °C); 30 cycles of denaturation (15 s, 98 °C), annealing (30 s, Tm of the lower primer) and elongation (based on PCR product length 0.5 kb/min, 72 °C); and final extension (10 min, 72 °C). For Taq: Initial denaturation (2 min, 95 °C); 30 cycles of denaturation (30 s, 95 °C), annealing (30 s, lower primer’s Tm) and elongation (based on PCR product length 1 kb/min, 72 °C); and final extension (10 min, 72 °C). PCR products or DNA fragments from restriction digests were purified by agarose gel electrophoresis and isolated using NucleoSpin Gel and PCR Clean-up kit (Macherey-Nagel). The digested PCR products were cloned into suitable plasmids pSBtn5Kan and verified by restriction analysis and sequencing. The PCR products cloned into pCR2.1 vector were adenylated prior to use of the TOPO TA Cloning Kit (Invitrogen). Details on the construction of all plasmids used and generated in this study are given in Supplementary Table 17.

For inactivation of *mxpA* gene, DNA fragment 001inact39A was PCR amplified from MCy9003 genomic DNA using 001-inact-nrps39A-F and 001-inact-nrps39A-R primers (Supplementary Figure 59). The DNA fragment was adenylated prior to cloning into pCR2.1 vector using TOPO TA Cloning Kit (Invitrogen), yielding pCR2.1-001inact39A plasmid. The orientation of the homology region in a plasmid was determined using PCR21R in combination with 001-inact-nrps39A-F and 001-inact-nrps39A-R primer. After the orientation confirmation of the homology region in the plasmid, the primer G001-inact39Ao1 was designed aligning to the genome in the vicinity of the homology. The plasmid was electroporated into MCy9003 WT, the transformants plated on CYHv3 agar plates supplemented with kanamycin 75 µg/mL and incubated at 30 °C for 6-10 days. Grown transformants were replated onto new CYHv3 Kn75 agar plates that were incubated at 30 °C. After 2-3 days, some cell mass was collected and resuspended in 300 µl ddH_2_O, and genomic DNA was extracted using Gentra Puregene Cell Kit. The plasmid location verification was done by PCR with two sets of primers: The homology itself was PCR-amplified as a positive control using 001-inact-nrps39A-F and 001-inact-nrps39A-R primers and the integration into desired gene was confirmed using G001-inact39Ao1 primer, located outside of homology region, in combination with PCR21R primer, aligning on the plasmid. Plasmid integration in MCy9003 was confirmed by fermentation of MCy9003 WT and MCy9003 intact-*mxpA* in CYHv3 medium (0.2% Bacto™ soytone, 0.3% Bacto™ casitone, 0.2% glucose, 0.3% soluble starch, 0.15% Bacto™ yeast extract, 0.05% CaCl_2_×2H_2_O, 0.1 % MgSO_4_×7H_2_O, 50 mM HEPES, 8 mg/L Fe-EDTA, dH_2_O; pH 7.2) with supplementation of 2 % XAD-16 resin and detection of the depleted myxopentacin A production by HPLC-MS.

## ACKNOWLEDGEMENTS

This work was partially funded by the Gottfried Wilhelm Leibniz Prize from the German Research Foundation (DFG, FKZ MU 1254/33-1 and MU 1254/32-1) and the Helmholtz Association. A.A.A. was partially funded by the HelmholtzAI project AMR-XAI. G.C. was partially funded by the Leibniz Science Campus “Living Therapeutic Materials”. O.V.K. acknowledges funding from the Klaus Faber Foundation. The authors thank Cathrin Spröer for genome sequencing and Vera Junker, Nicole Heyer, Aileen Gollasch and Birte Trunkwalter for excellent technical assistance. This work was partially funded by the German Center for Infection Research (DZIF) grant no. TTU-09.720. We thank the Citizen scientists who participated in the MICROBELIX sample collection project for their contribution of soil samples from Germany. Furthermore, we thank Prof. Snædís Huld Björnsdóttir from University of Iceland for support with sample collection and Dr. F. P. Jake Haeckl for fruitful discussion.

## AUTHOR CONTRIBUTIONS

R.G. isolated, cultivated, characterized, and fermented the myxobacterial strains and contributed to the revision of the taxonomy. M.V.E.G. contributed to strain isolation, cultivation, and characterization. A.A.A. performed bioinformatics analysis to analyze the biosynthetic potential of *Myxococcota*. G.C. performed the computational taxonomic analysis and contributed to the revision of taxonomy. T.M. contributed to bioinformatics genome analysis. P.S. performed taxonomic comparison of *Actinomycetota* and *Myxococcota*. C.D.B., A.P., M.H., and S.W. performed *in silico* analysis of the BGCs. A.P. and M.H. performed inactivation and feeding experiments. C.F., A.P., E.O., H.Z., and S.W. have isolated and characterized natural products. S.K., P.H., A.T., A.G., and A.K. created and curated data for the ABC-Myxo web resource. J.B., B.B., J.O., and U.N. performed bioinformatics metabolic analysis. A.A.A., R.G., G.C., C.D.B., and D.K. wrote the manuscript. O.V.K. and R.M. devised the study, wrote the manuscript, and provided funding. All authors reviewed and edited the manuscript and approved the final version.

