## Supporting Information for "Unveiling the taxonomic diversity and unprecedented biosynthetic treasure of the phylum *Myxococcota*"

Supplementary Information

### List of Supplementary Notes

**Supplementary Note 1** Reclassification suggestions……………………………………………8-10
**Supplementary Note 2** Structure elucidation of myxolutamid A and identification of the putative *mxl* BGC……………………………………………………………………………………………….25
**Supplementary Note 3** Structure elucidation of sorangicin X and Y…………………………….36
**Supplementary Note 4** Analysis of the *srx* BGC and comparison to the *sor* BGC…………54-55
**Supplementary Note 5** Selecting the clustering thresholds for BiG-SCAPE and BiG-SLiCE………………………………………………………………………………………61-62
**Supplementary Note 6** Selection of *Actinomycetota* strains comparable to the in-house *Myxococcota* based on 16S DNA sequence…………………………………………………64
**Supplementary Note 7** Structure elucidation of myxopentacin A………………………………...68
**Supplementary Note 8** Biosynthesis of myxopentacin……………………………………………77

### List of Supplementary Tables

**Supplementary Table 1** Genome assembly statistics. (see separate file)................................
**Supplementary Table 2** Type strain list (see separate file).....................................................
**Supplementary Table 3** Classifications for the 154 new strains. (see separate file) ................
**Supplementary Table 4** Reclassified *Mycoccocota* Taxonomy (see separate file)...................

**Supplementary Figure 60** Upset plot showing the number of unique and shared GCFs between different sub-orders belonging to Myxococcota phylum…………….……………………………..82

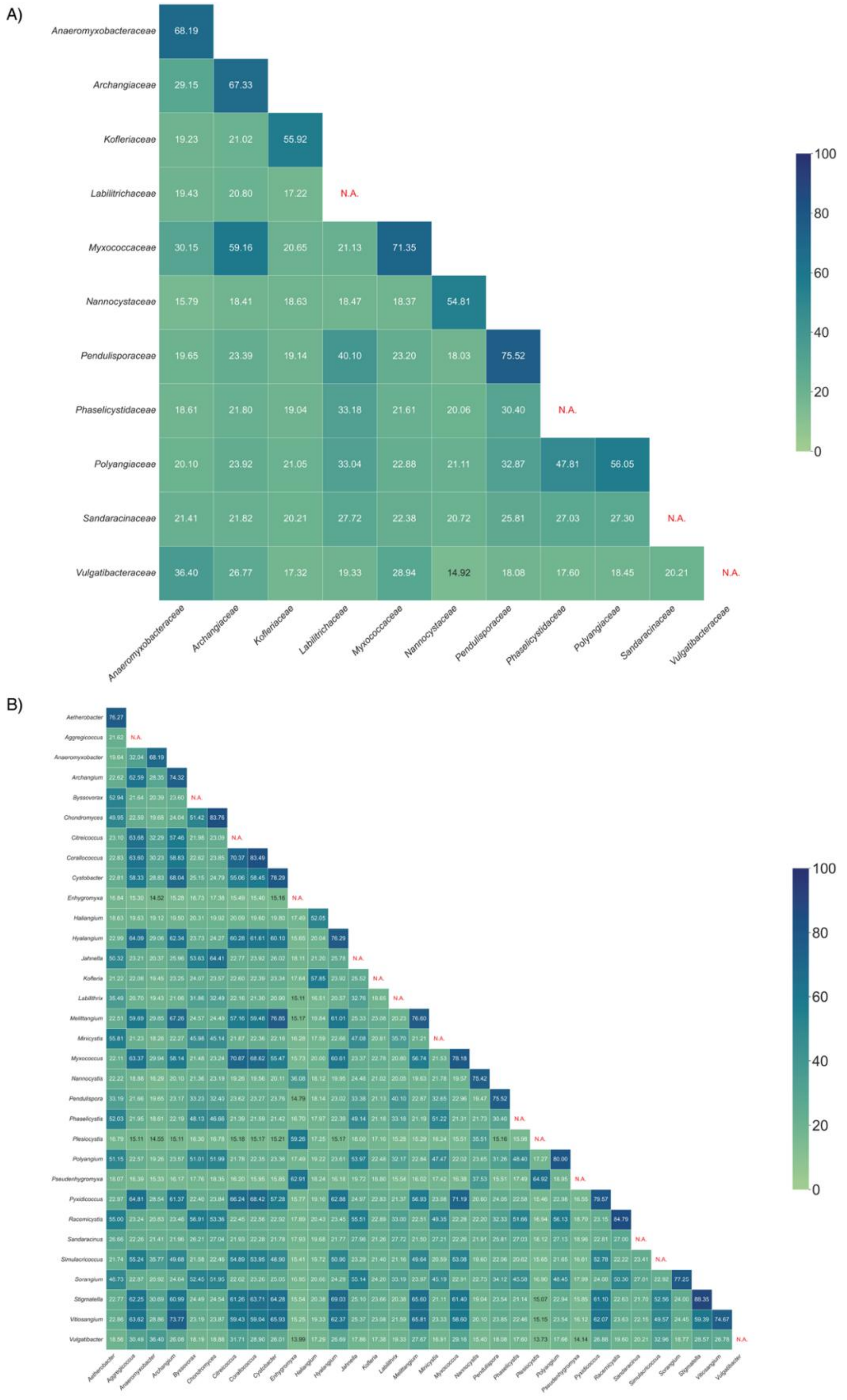

**Supplementary Figure 1** The averaged pairwise POCP values of (A) families and (B) genera calculated by *Myxococcota* type strains. Single-member taxa, for which intra-taxon POCP cannot be calculated, are indicated as ‘N.A.’.

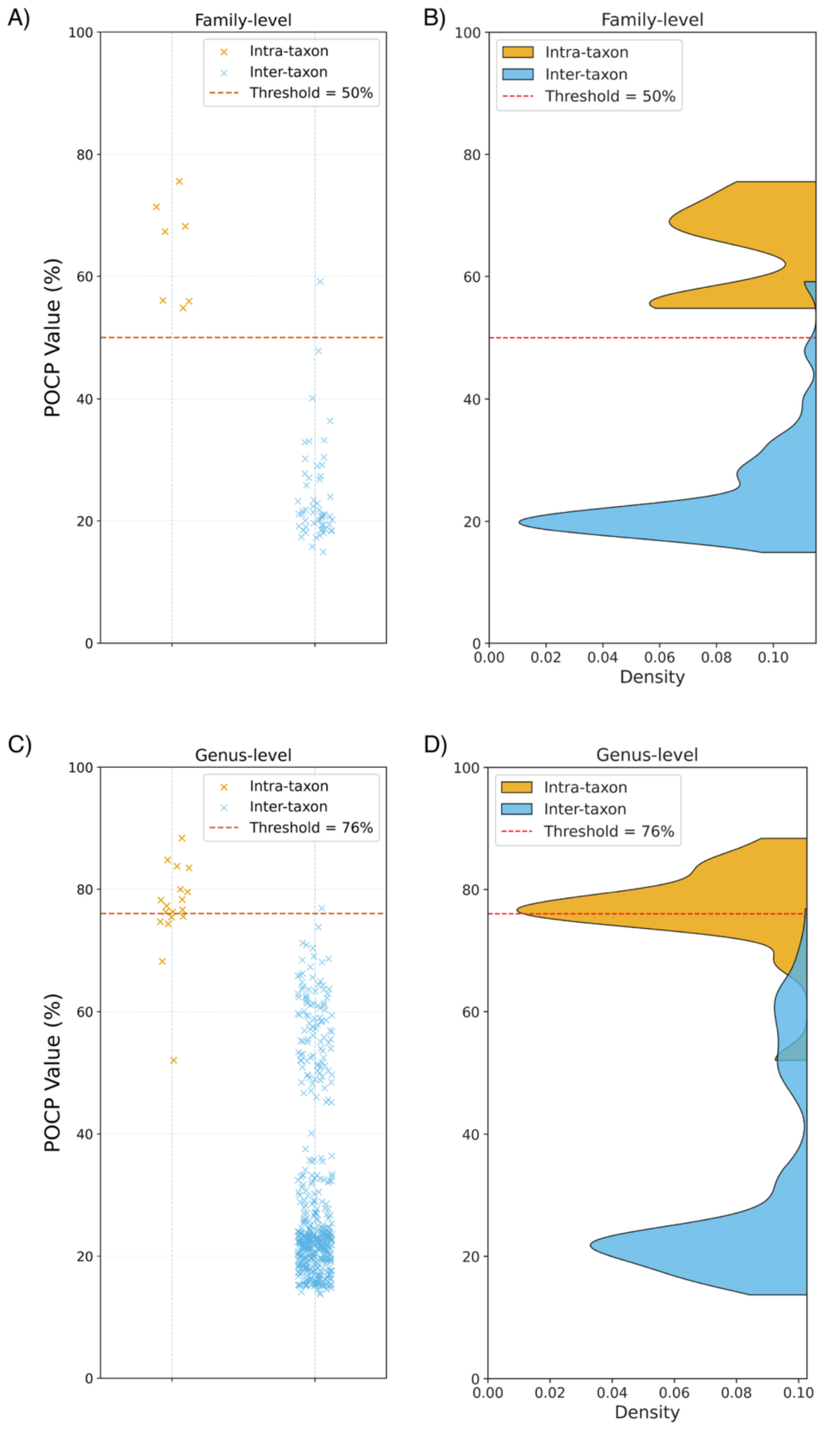

**Supplementary Figure 2** Distribution of average inter- and intra-species POCP values on genus and family levels of *Myxococcota* type-strains. (A) Scatter plot of average inter- and intra-species POCP values on family level; (B) Kernal density plot of average inter- and intra-species POCP values on family level; (C) Scatter plot of average inter- and intra-species POCP values on genus level; (D) Kernal density plot of average inter- and intra-species POCP values on genus level.

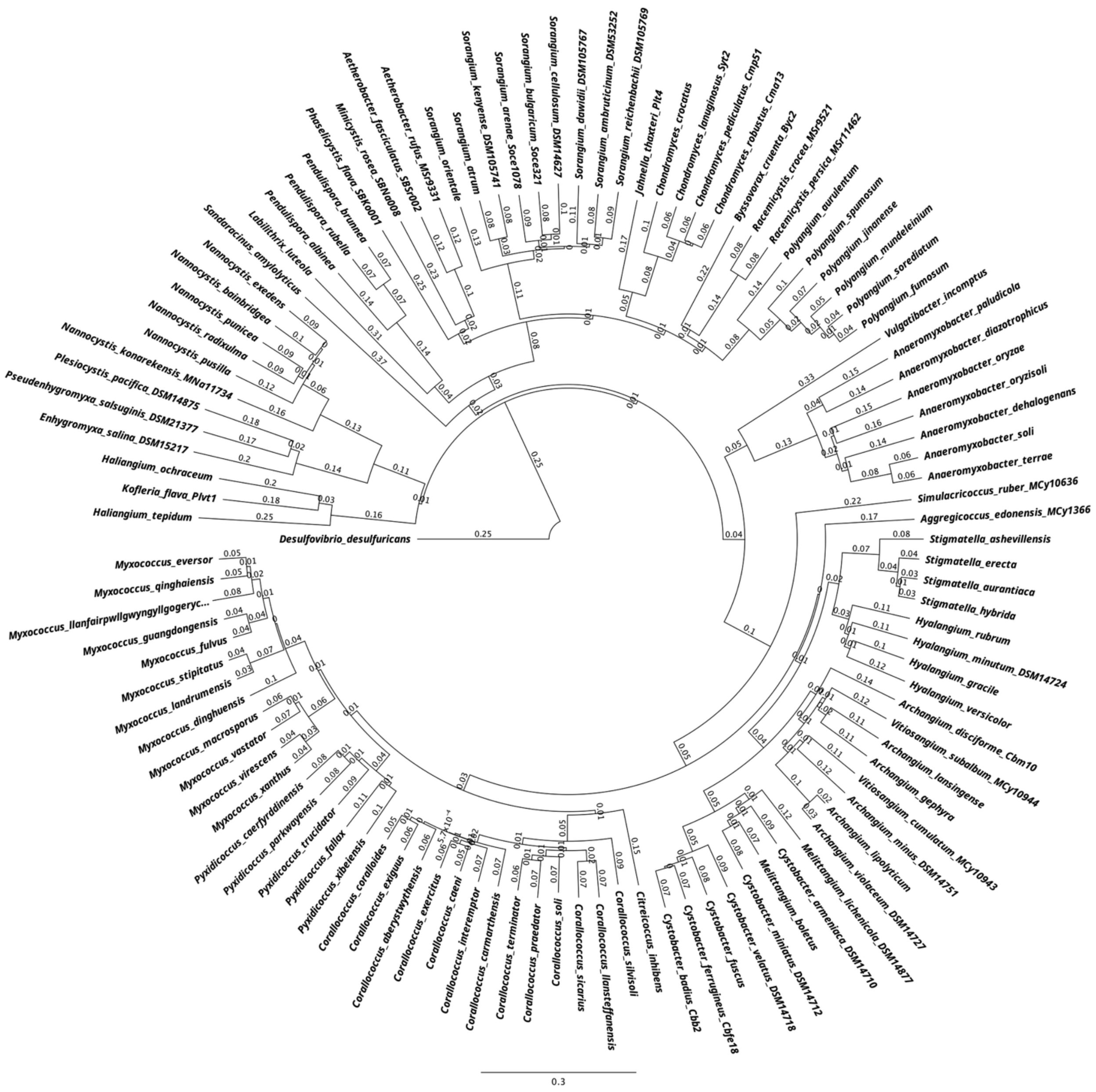

**Supplementary Figure 3** Phylogenomic reconstruction by the 109 *Myxococcota* type strains using BIONJ from the pairwise POCP distances. *Desulfovibrio desulfuricans* serves as the outgroup. Leaves are annotated with the established taxonomy at species level.

**Supplementary Note 1** Reclassification suggestions.

**Reclassification suggestions on the Family level**

- *Myxococcaceae* and *Archangiaceae* united as one family *Myxococcaceae* (Supplementary Figure 4)
- *Enhygromyxa*, *Plesiocystis*, and *Pseudenhygromyxa* belong a new family *New Family 8*, separated from *Nannocystaceae* (Supplementary Figure 5)
- *Labilitrichaceae*, *Phaselicystaceae,* and *Polyangiaceae* are three separate families. *Aetherobacter* and *Minicystis* belong to *Phaselicystaceae* (Supplementary Figure 6)

**Reclassification suggestions on the Genus level**

- ***Anaeromyxobacter* split into 6 genera (Supplementary Figure 7):**

  (Original) *Anaeromyxobacter*:
  Species: *Anaeromyxobacter dehalogenans* (Type Species)
  *Anaeromyxobacter*-derived New Genus 1:
  Species: *Anaeromyxobacter diazotrophicus*
  *Anaeromyxobacter*-derived New Genus 2:
  Species: *Anaeromyxobacter oryzae*
  *Anaeromyxobacter*-derived New Genus 3:
  Species: *Anaeromyxobacter oryzisoli*
  *Anaeromyxobacter*-derived New Genus 4:
  Species: *Anaeromyxobacter paludicola*
  *Anaeromyxobacter*-derived New Genus 5:
  Species: *Anaeromyxobacter soli, Anaeromyxobacter terrae*

- ***Haliangium* split into 2 genera (Supplementary Figure 8):**
  (Original) *Haliangium*:
  Species: *Haliangium ochraceum*
  *Haliangium*-derived New Genus 1:
  Species: *Haliangium tepidum*

- ***Myxococcus* split into 2 genera (Supplementary Figure 9):**
  (Original) *Myxococcus*:
  Species: *Myxococcus stipitatus*, *Myxococcus landrumensis*, *Myxococcus guangdongensis*, *Myxococcus fulvus* (Type Species), *Myxococcus llanfairpwllgwyngyllgogerychwyrndrobwllllantysiliogogogochensis, Myxococcus qinghaiensis, Myxococcus eversor, Myxococcus dinghuensis*
  *Myxococcus*-derived New Genus 1:
  Species: *Myxococcus xanthus*, *Myxococcus virescens*, *Myxococcus vastator*, *Myxococcus macrosporus*

- ***Nannocystis* split into 2 genera (Supplementary Figure 10):**
  (Original) *Nannocystis*:

Species: *Nannocystis bainbridgea*, *Nannocystis exedens* (Type Species), *Nannocystis punicea*, *Nannocystis pusilla*, *Nannocystis radixulma*

*Nannocystis*-derived New Genus 1:

Species: *Nannocystis konarekensis*

- ***Pendulispora* split into 2 genera (Supplementary Figure 11):**

(Original) *Pendulispora*:

Species: *Pendulispora albinea*

*Pendulispora*-derived New Genus 1:

Species: *Pendulispora brunnea*, *Pendulispora rubella*

- ***Polyangium* split into 2 genera (Supplementary Figure 12):**

(Original) *Polyangium*:

Species: *Polyangium fumosum*, *Polyangium jinanense*, *Polyangium mundeleinium*, *Polyangium sorediatum*, *Polyangium spumosum* (Type Species), *Polyangium vitellinum*'s genome is not available. We refer it as *Polyangium* based on previous publication[1].

*Polyangium*-derived New Genus 1:

Species: *Polyangium aurulentum*

- ***Sorangium* split into 4 genera (Supplementary Figure 13):**

(Original) *Sorangium:*

Species: *Sorangium arenae, Sorangium bulgaricum, Sorangium cellulosum* (Type Species)

*Sorangium*-derived New Genus 1:

Species: *Sorangium atrum*, *Sorangium kenyense*

*Sorangium*-derived New Genus 2:

Species: *Sorangium dawidii*, *Sorangium ambruticinum*, *Sorangium reichenbachii*

*Sorangium*-derived New Genus 3:

Species: *Sorangium orientale*

- ***Archangium-Vitiosangium* complex (Supplementary Figure 14):**

Reclassification back of *Archangium disciforme* to the original genus *Angiococcus*

Genus *Archangium* includes the following species:

*Archangium gephyra* (Type species)
*Archangium lansingense*
*Vitiosangium subalbum*, reclassify to *Archangium subalbum*

Genus *Vitiosangium* includes the following species:

*Vitiosangium cumulatum* (Type species)
*Archangium minus*, reclassify to *Vitiosangium minus*
*Archangium violaceum*, reclassify to *Vitiosangium violaceum*
*Archangium*. lipolyticum, reclassify to *Vitiosangium lipolyticum*

- ***Cystobacter-Melittangium* complex (Supplementary Figure 15):**

  Genus *Cystobacter* includes the following species:

  *Cystobacter fuscus* (Type species)
  *Cystobacter ferrugineus*
  *Cystobacter badius*
  *Cystobacter velatus*

  Genus *Melittangium* includes the following species:

  *Melittangium lichenicola*
  *Melittangium boletus* (Type species)
  *Cystobacter miniatus*, reclassify to *Melittangium miniatus*
  *Cystobacter armeniaca*, reclassify to *Melittangium armeniaca*
  *Melittangium alboraceum* (Type strain genome not available) We refer it as *Melittangium* based on previous publication[2]

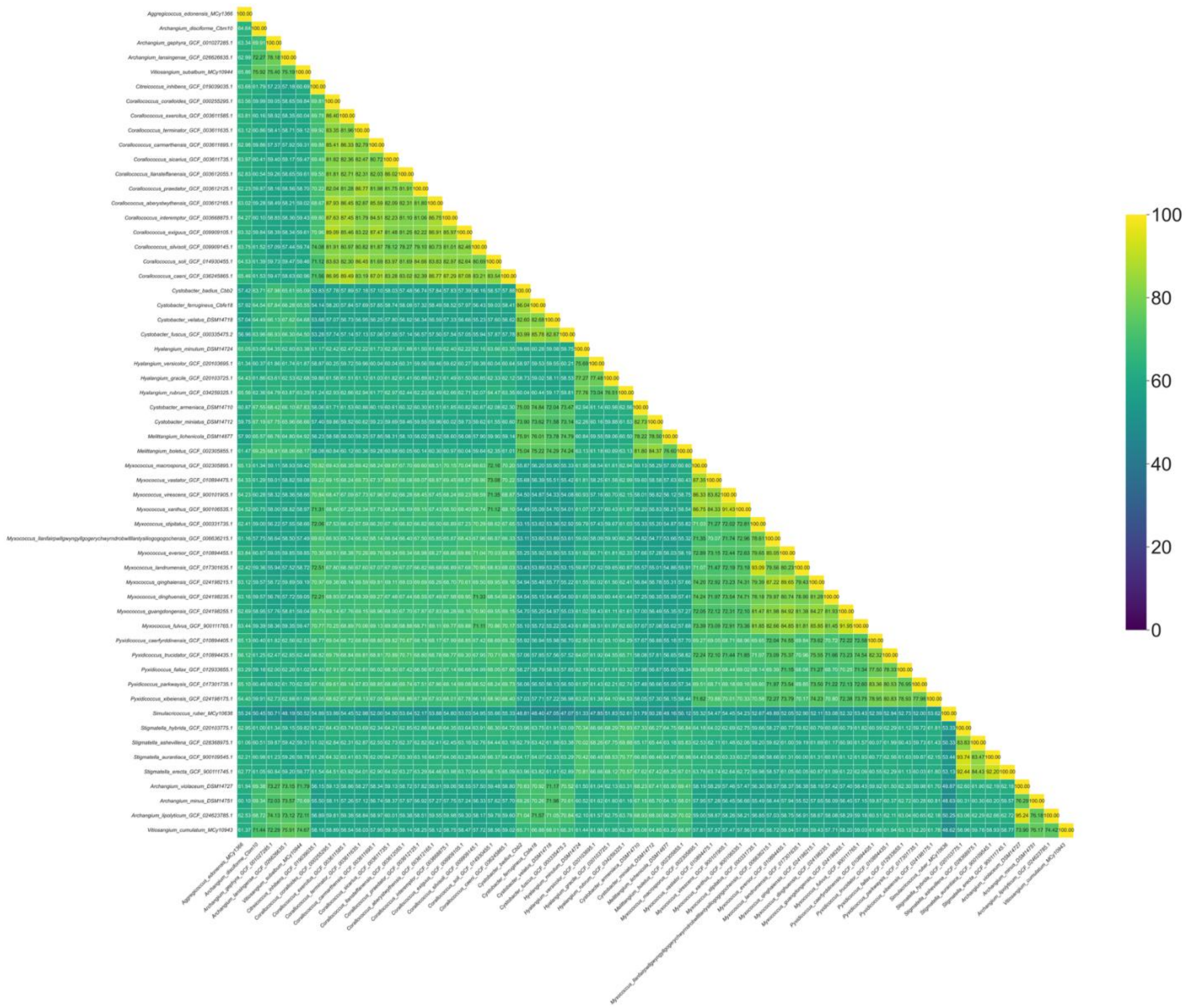

**Supplementary Figure 4** The reclassified family *Myxococcaceae*. The pairwise POCP values of species within the family *Myxococcaceae* and *Archangiaceae* leading to a suggested merge to one family *Myxococcaceae.* All species reclassified as *Myxococcaceae* shown here.

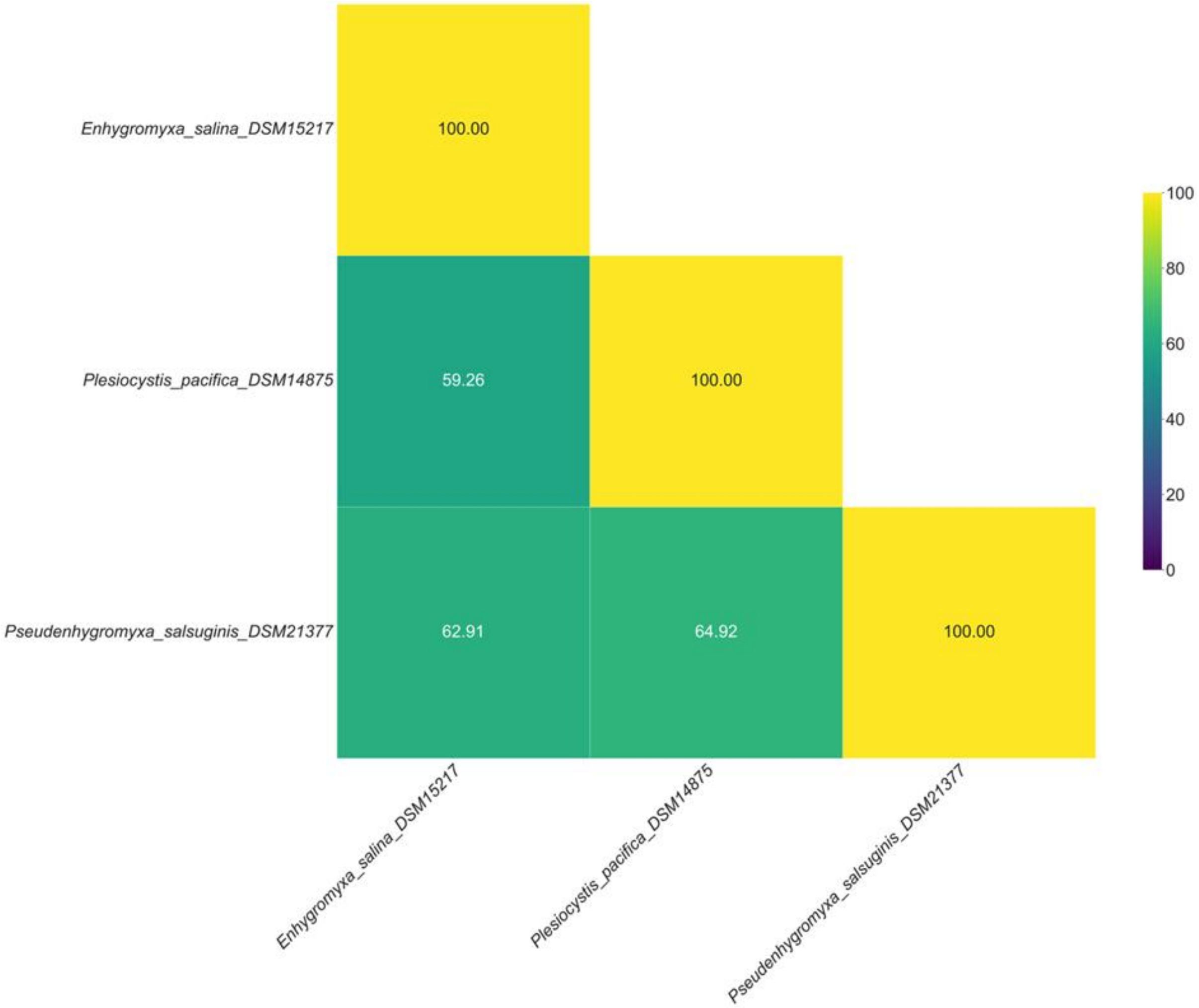

**Supplementary Figure 5** The New Family 8. The pairwise POCP values of species within *Enhygromyxa, Plesiocystis, and Pseudenhygromyxa* suggest they form a new family (New Family 8) separated from *Nannocystaceae.* All species reclassified as New Family 8 are shown here.

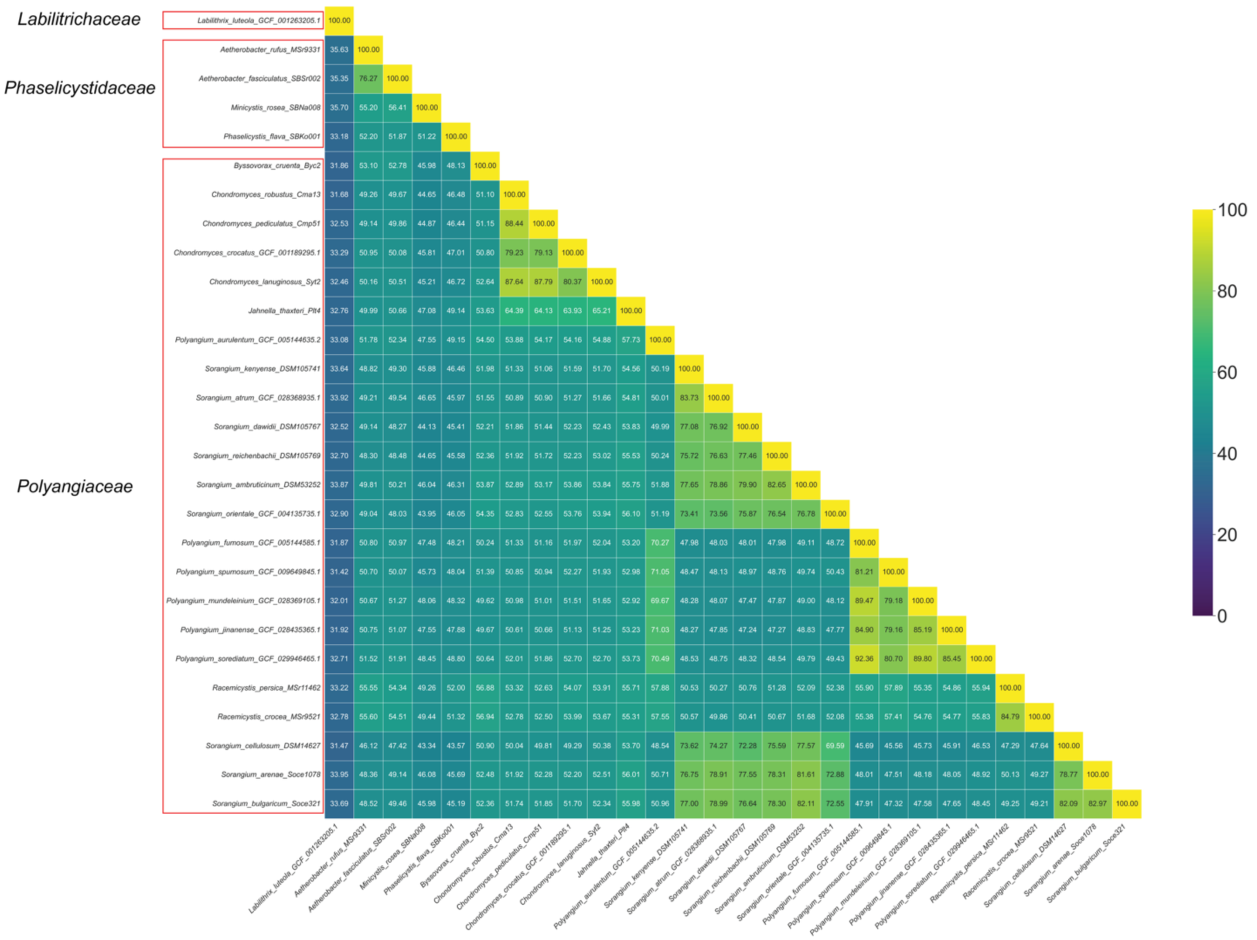

**Supplementary Figure 6** The reclassified family *Labilitrichaceae*, *Phaselicystaceae* and *Phaselicystacea.* The pairwise POCP values of species within *Labilitrichaceae, Phaselicystaceae, and Polyangiaceae* suggest they are three separate families. *Aetherobacter* and *Minicystis* belong to *Phaselicystaceae.* All species reclassified as *Labilitrichaceae*, *Phaselicystaceae* and *Phaselicystaceae* shown here.

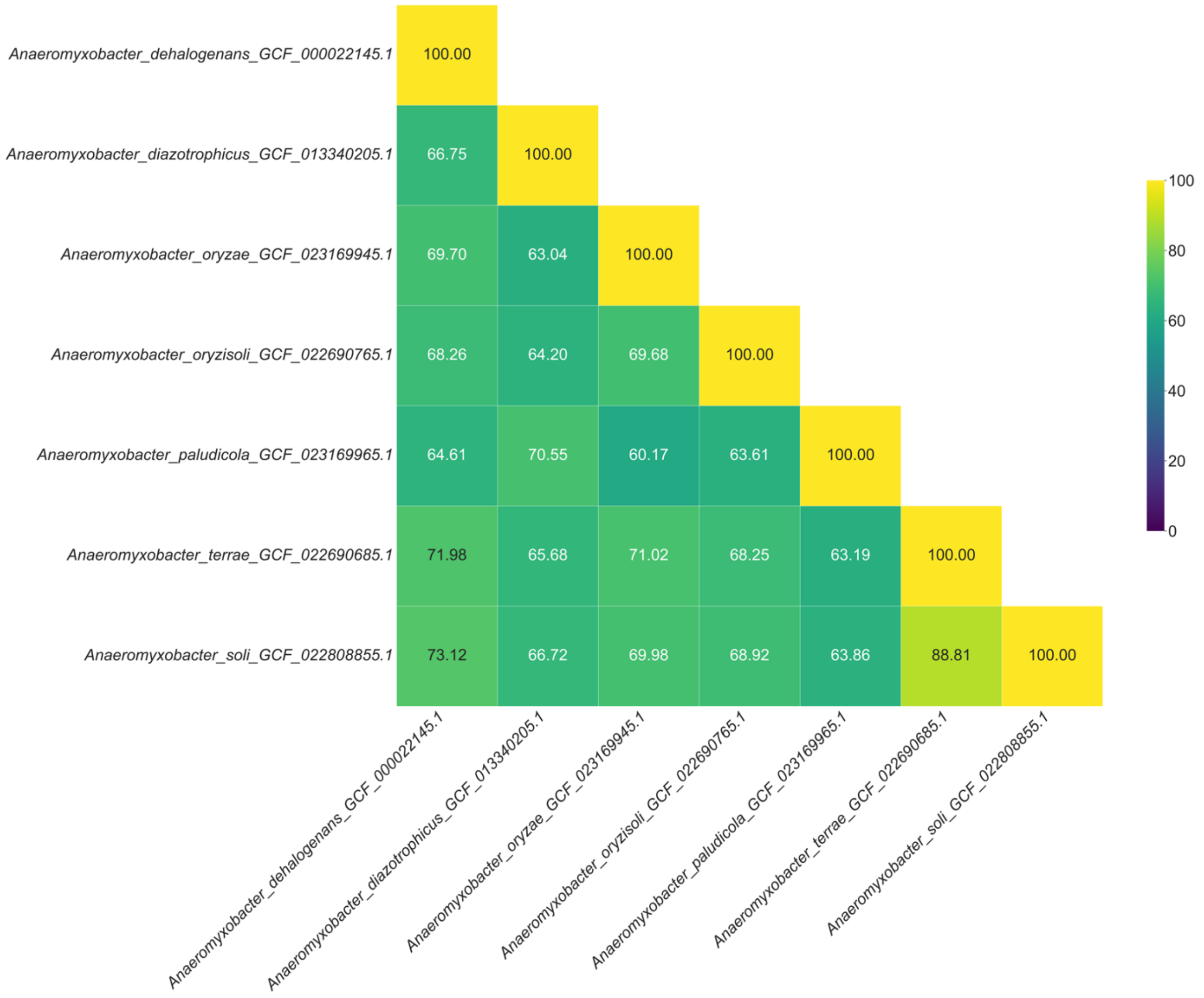

**Supplementary Figure 7** The reclassified genus *Anaeromyxobacter.* The pairwise POCP values of species within *Anaeromyxobacter* suggest they are six separate genera: *Anaeromyxobacter* (represented by *Anaeromyxobacter dehalogenans* (Type species)), *New Genus 1* (represented by *Anaeromyxobacter diazotrophicus*), *New Gebnus 2* (represented by *Anaeromyxobacter oryzae*), *New Genus 3* (represented by *Anaeromyxobacter oryzisoli*), *New Genus 4* (represented by *Anaeromyxobacter paludicola*), *New Genus 5* (represented by *Anaeromyxobacter soli and Anaeromyxobacter terrae*).

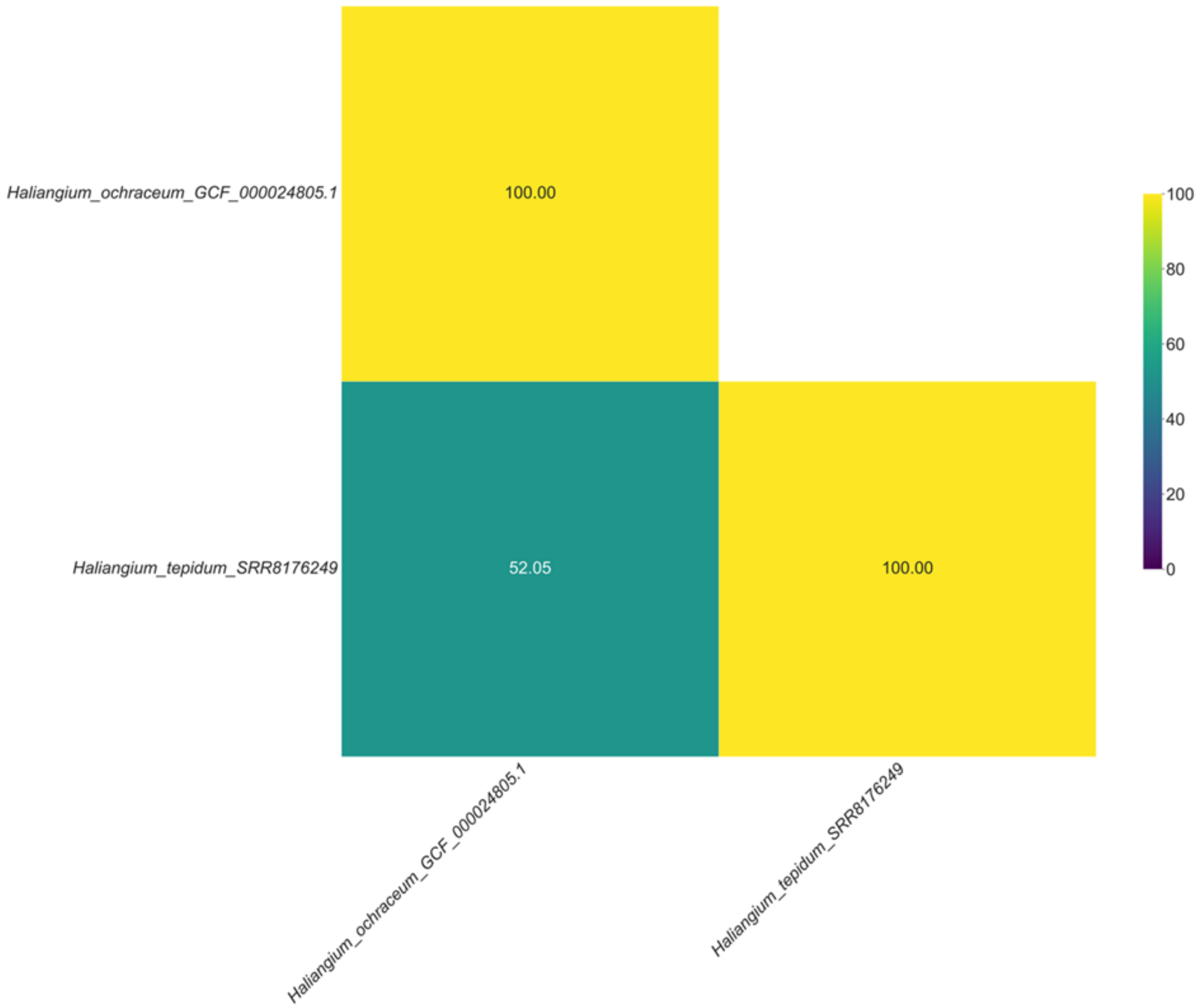

**Supplementary Figure 8** The reclassified genus *Haliangium.* The pairwise POCP values of species within *Haliangium* suggest splitting the genus in two genera: *Haliangium* (represented by *Haliangium ochraceum)* and New Genus 1 (represented by *Haliangium tepidum)*.

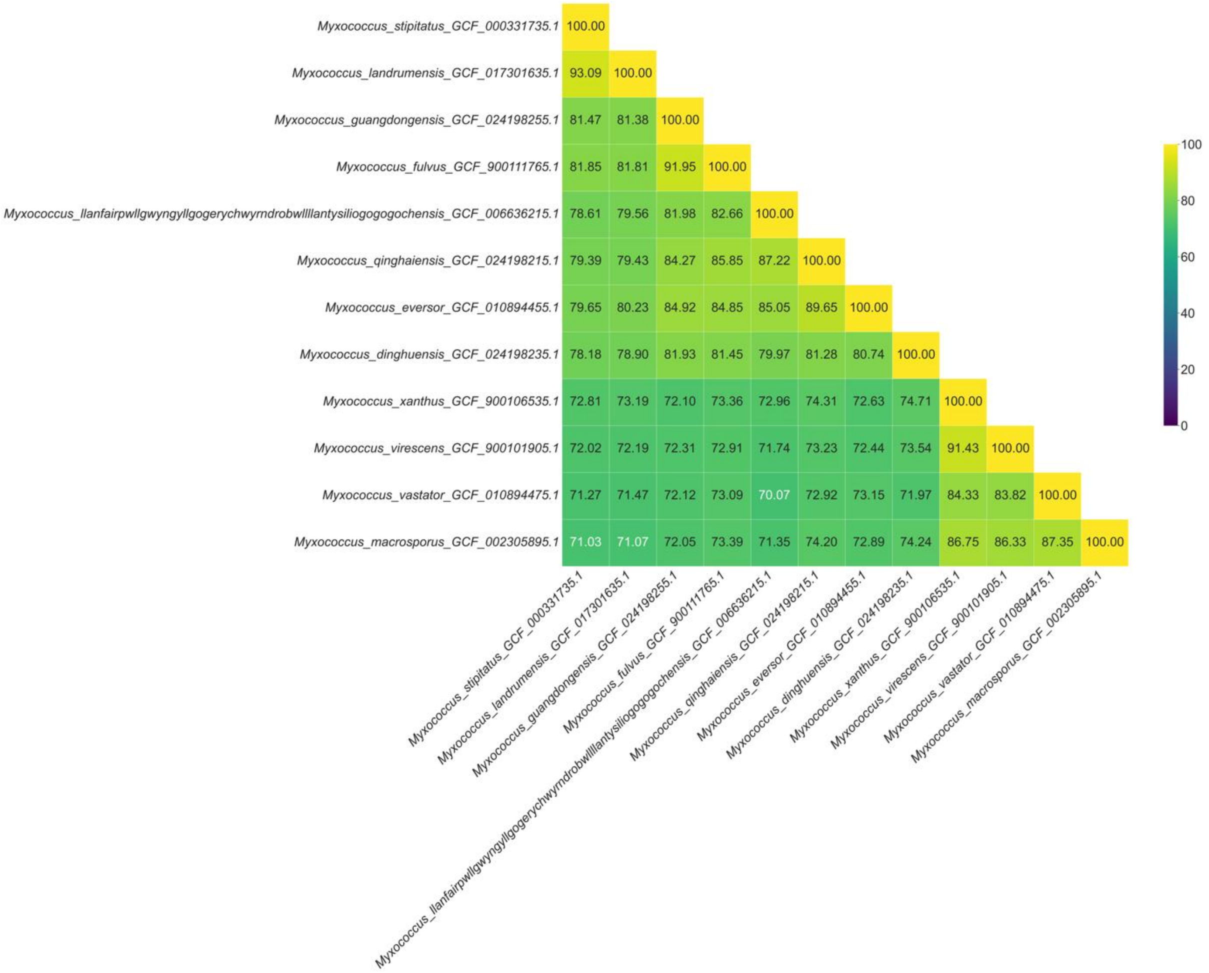

**Supplementary Figure 9** The reclassified genus *Myxococcus.* The pairwise POCP values of species within *Myxococcus* suggest splitting the genus in two genera: *Myxococcus* (represented by *Myxococcus stipitatus*, *Myxococcus landrumensis*, *Myxococcus guangdongensis*, *Myxococcus fulvus* (Type species), *Myxococcus llanfairpwllgwyngyllgogerychwyrndrobwllllantysiliogogogochensis, Myxococcus qinghaiensis, Myxococcus eversor,* and *Myxococcus dinghuensis)* and New Genus 1 (represented by *Myxococcus xanthus*, *Myxococcus virescens*, *Myxococcus vastator*, *Myxococcus macrosporus)*

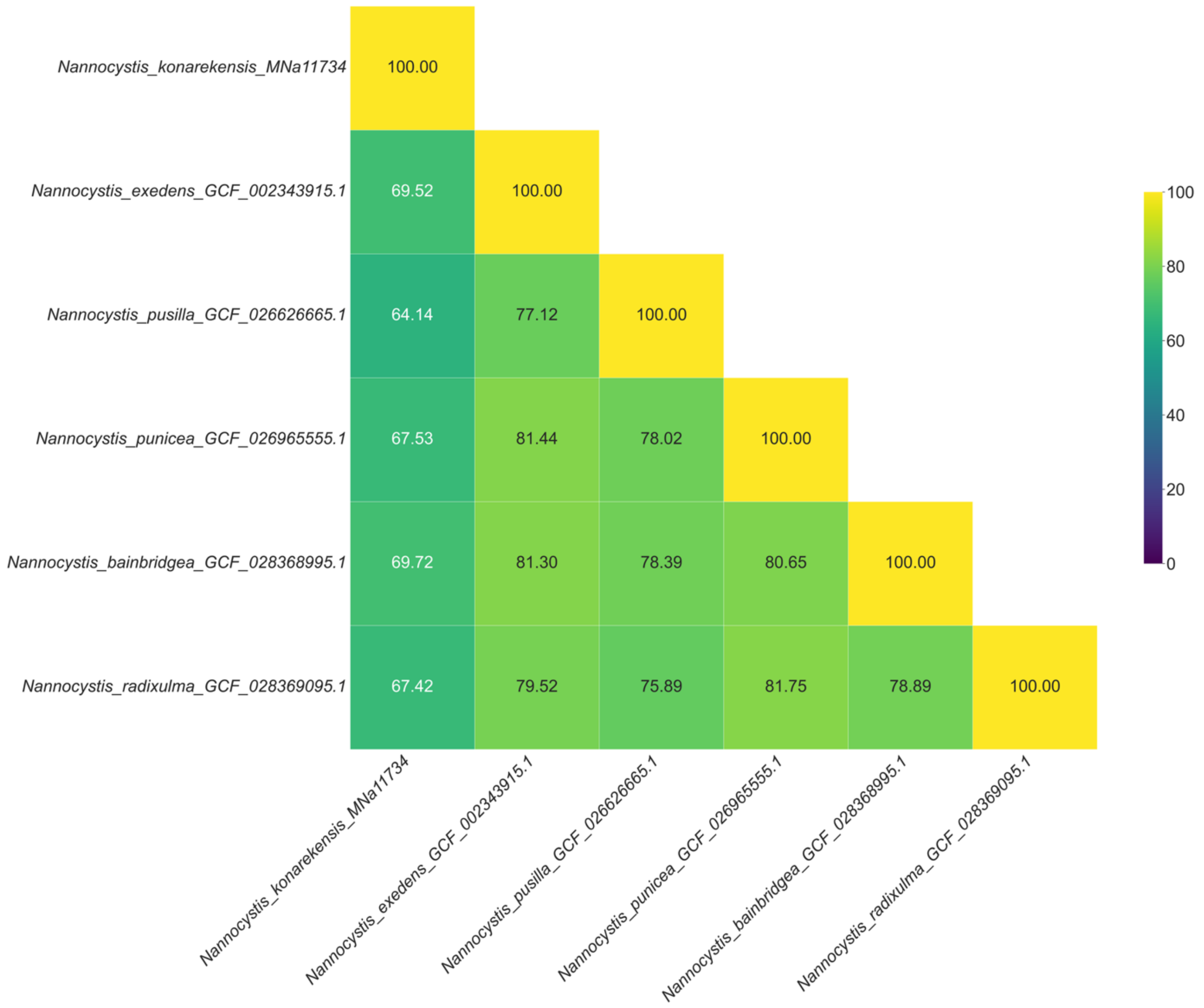

**Supplementary Figure 10** The reclassified genus *Nannocystis.* The pairwise POCP values of species within *Nannocystis* suggest splitting the genus in two genera: *Nannocystis* (represented by *Nannocystis bainbridgea*, *Nannocystis exedens* (Type Species), *Nannocystis punicea*, *Nannocystis pusilla* and *Nannocystis radixulma)* and New Genus 1 (represented by *Nannocystis konarekensis).*

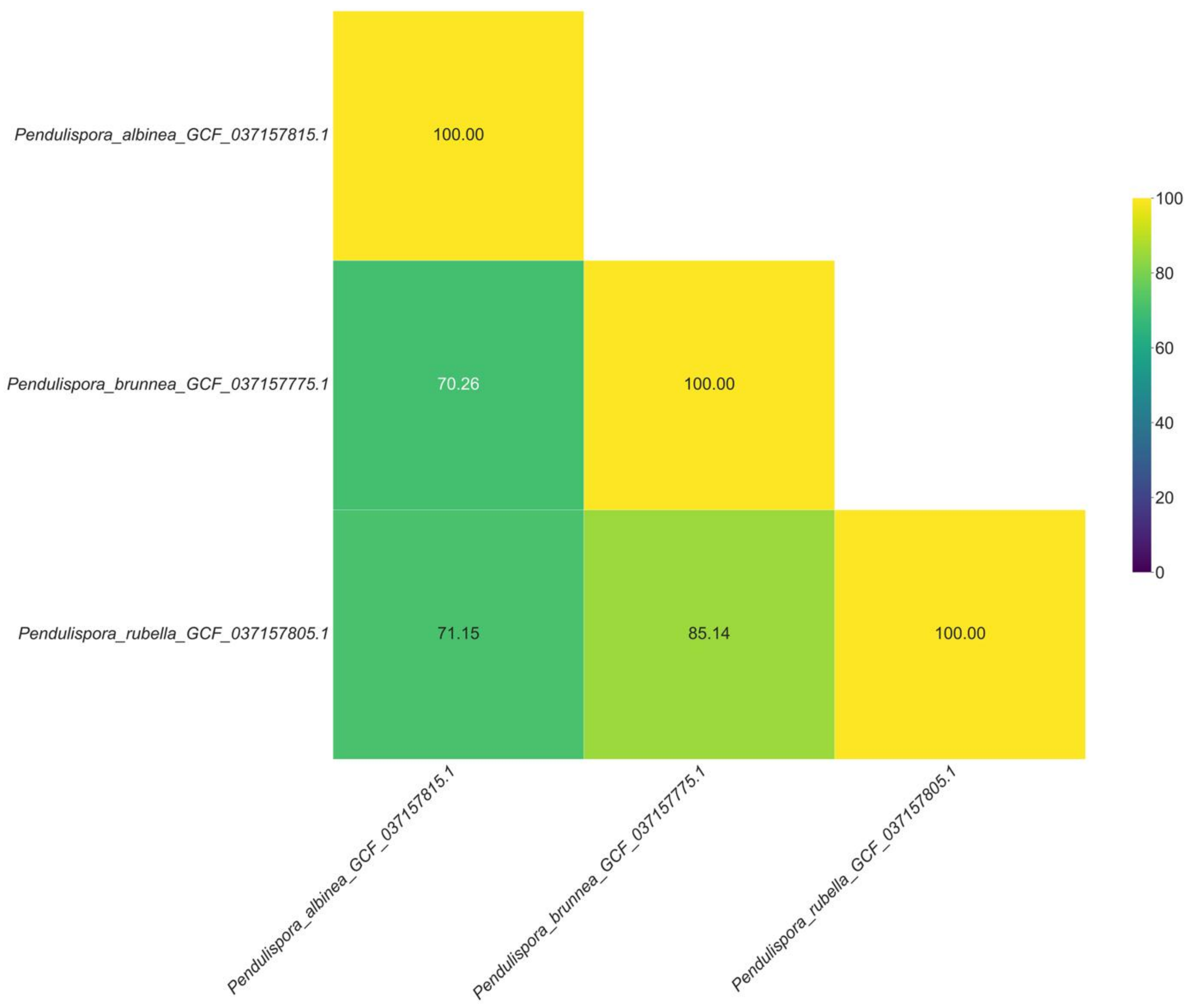

**Supplementary Figure 11** The reclassified genus *Pendulispora.* The pairwise POCP values of species within *Pendulispora* suggest splitting the genus in two genera: *Pendulispora* (represented by *Pendulispora albinea* and New Genus 1 (represented by *Pendulispora brunnea* and *Pendulispora rubella).*

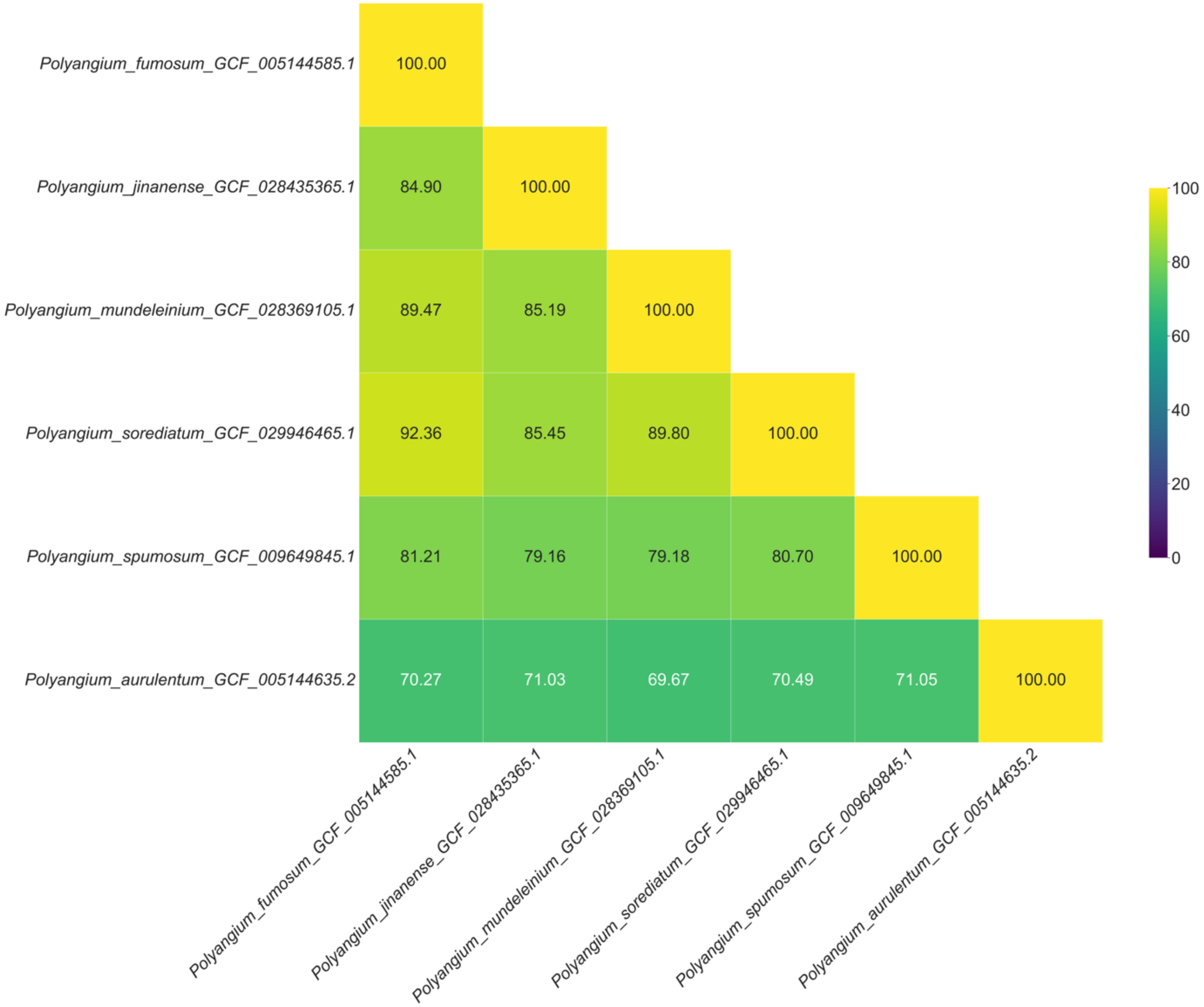

**Supplementary Figure 12** The reclassified genus *Polyangium.* The pairwise POCP values of species within *Polyangium* suggest splitting the genus in two genera: *Polyangium* (represented by *Polyangium fumosum, Polyangium jinanense, Polyangium mundeleinium, Polyangium sorediatum, Polyangium spumosum* (Type species)) and New Genus 1 (represented by *Polyangium aurulentum).*

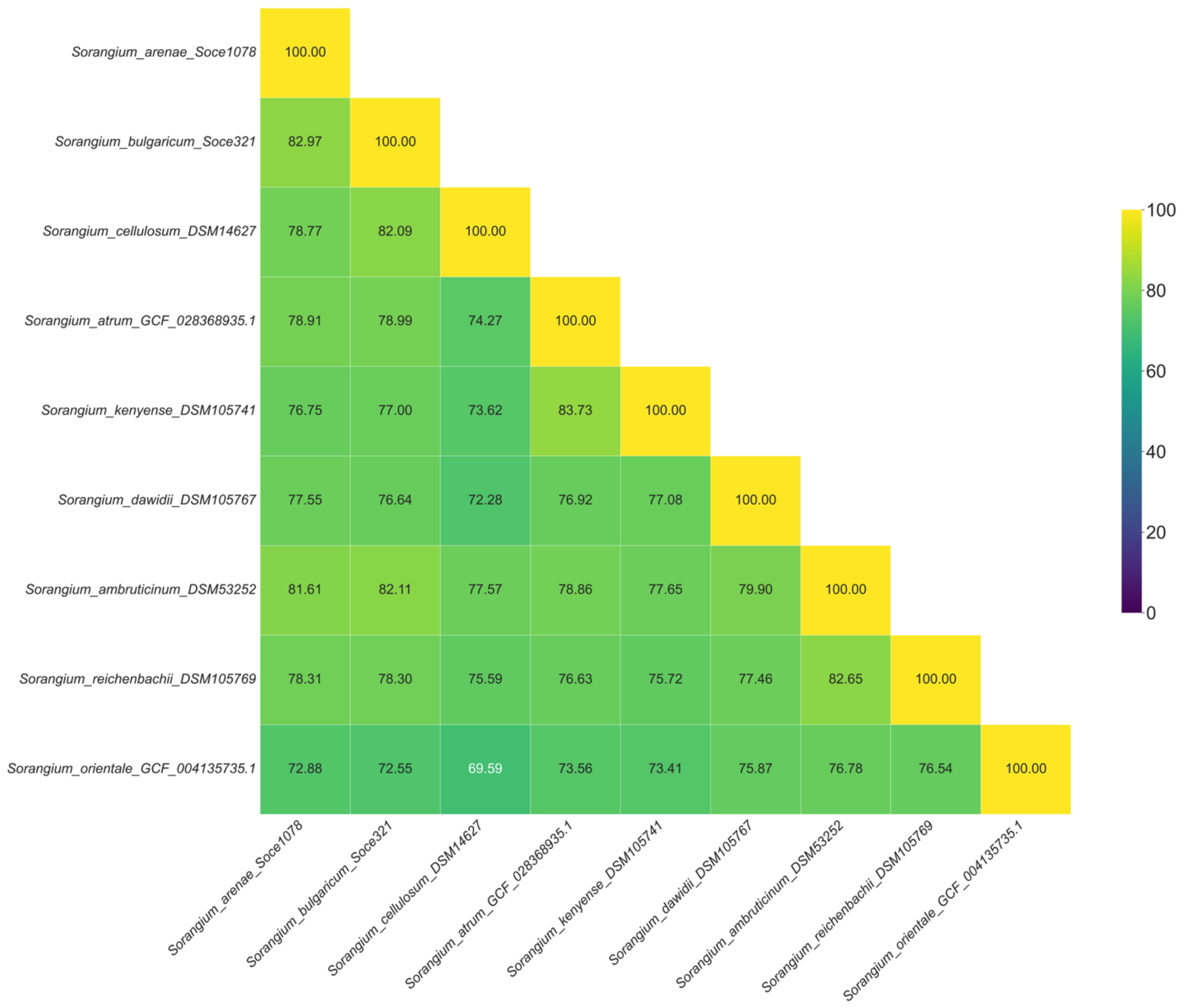

**Supplementary Figure 13** The reclassified genus *Sorangium.* The pairwise POCP values of species within *Sorangium* suggest splitting the genus in four genera: *Sorangium* (represented by *Sorangium arenae, Sorangium bulgaricum, Sorangium cellulosum* (Type species)), New Genus 1 (represented by *Sorangium atrum, Sorangium kenyense),* New Genus 2 (represented by *Sorangium dawidii, Sorangium ambruticinum, Sorangium reichenbachii),* New Genus 3 (represented by *Sorangium orientale).*

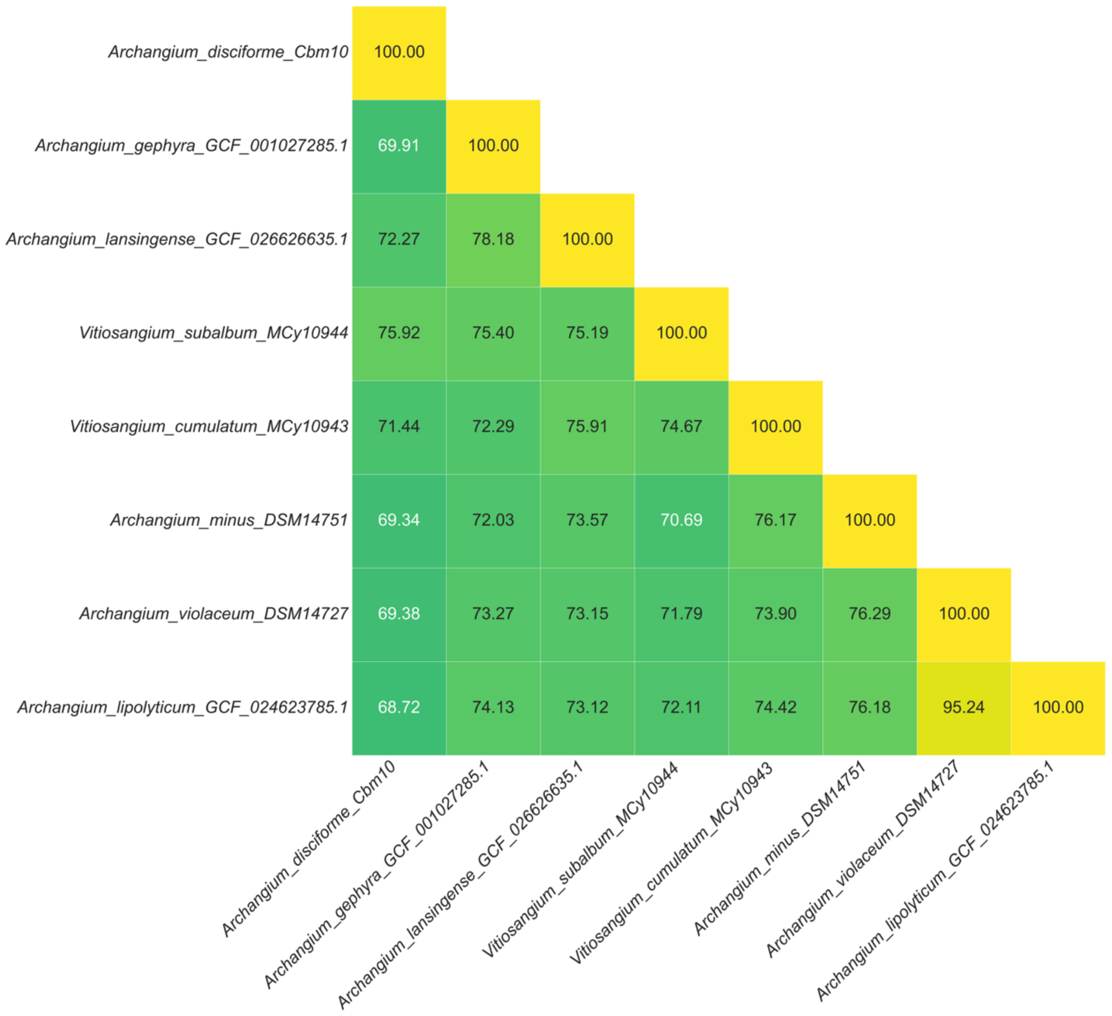

**Supplementary Figure 14** The reclassified genera *Archangium* and *Vitiosangium.* The pairwise POCP values of species within *Archangium* and *Vitiosangium* suggest *Archangium gephyra* (Type species), *Archangium lansingense*, *Vitiosangium* subalbum (reclassify to *Archangium subalbum*) beong to the genus *Archangium*, while *Vitiosangium cumulatum* (Type species), *Archangium minus* (reclassify to *Vitiosangium minus*), *Archangium violaceum* (reclassify to *Vitiosangium violaceum*), *Archangium*. lipolyticum (reclassify to *Vitiosangium lipolyticum*) belong to the genus *Vitiosangium*.

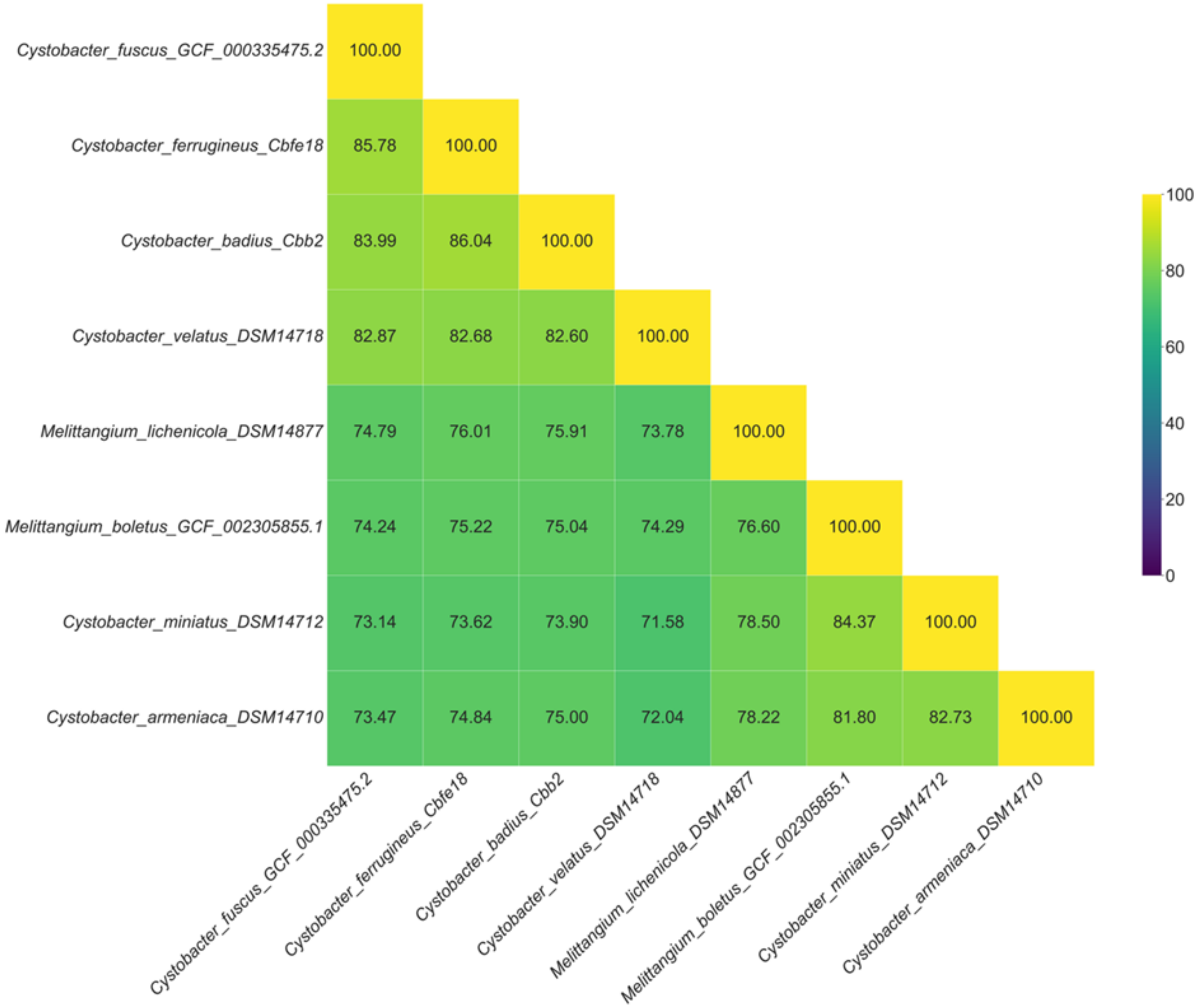

**Supplementary Figure 15** The reclassified genera *Cystobacter* and *Melittangium*. The pairwise POCP values of species within *Cystobacter* and *Melittangium* suggest *Cystobacter fuscus* (Type species)*, Cystobacter ferrugineus, Cystobacter badius* and *Cystobacter velatus* to belong to the genus Cystobacter, while *Melittangium lichenicola, Melittangium boletus* (Type species)*, Cystobacter miniatus* (reclassified to *Melittangium miniatus*), *Cystobacter armeniaca* (reclassify to *Melittangium armeniaca*) and *Melittangium alboraceum* belong to the genus *Melittangium.*

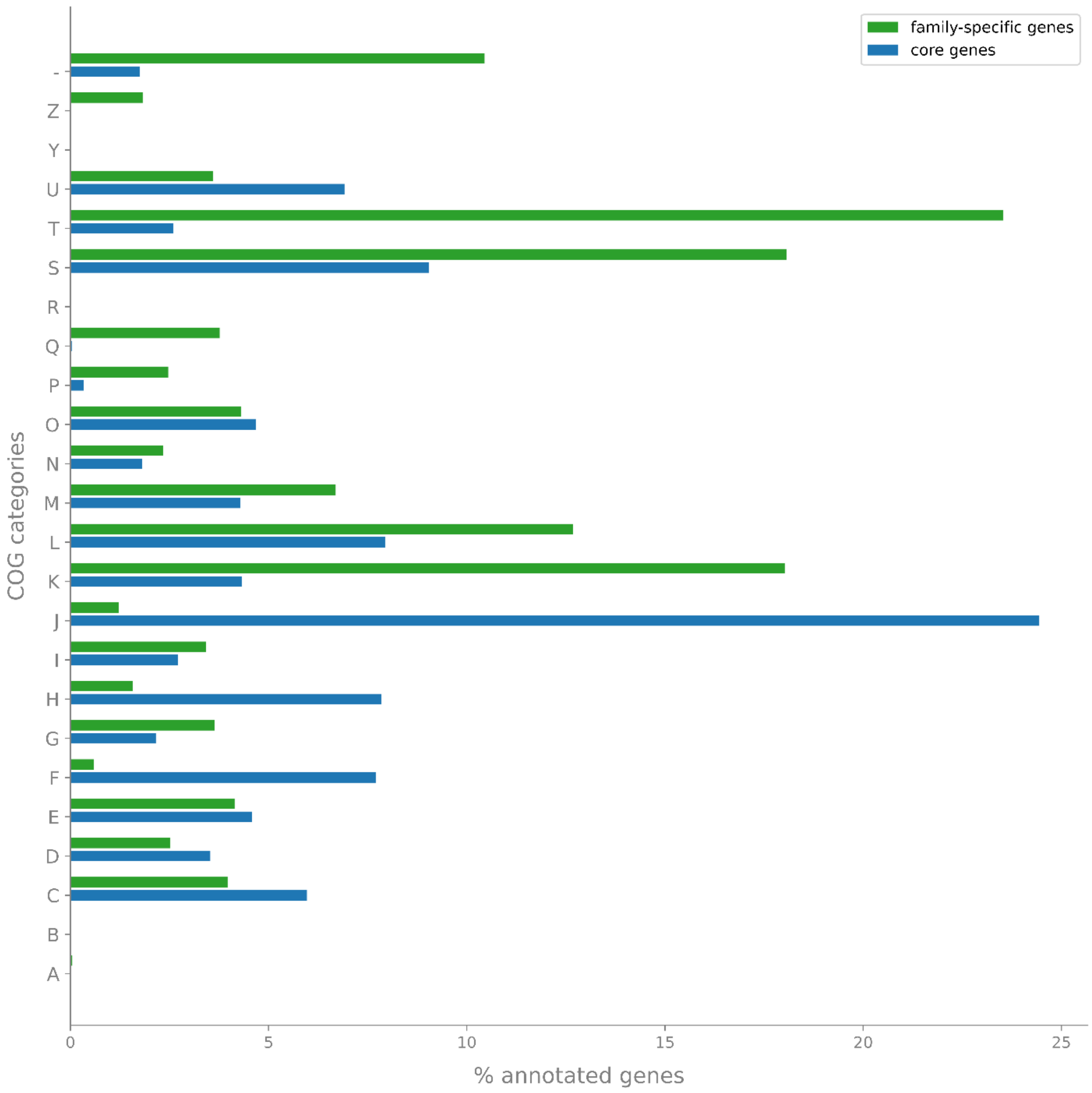

**Supplementary Figure 16** Distribution of COG (Clusters of Orthologous Groups) categories among core genes (blue) and family-specific genes (green). Core genes are shared by all 153 myxobacterial genomes. The y-axis shows for each COG category on the x-axis its percentage of the annotated genes across the family-specific genes or the core genes. COG-categories:

| | |
|---|---|
| J | Translation, ribosomal structure and biogenesis |
| A | RNA processing and modification |
| K | Transcription |
| L | Replication, recombination and repair |
| B | Chromatin structure and dynamics |
| D | Cell cycle control, cell division, chromosome partitioning |
| Y | Nuclear structure |
| V | Defense mechanisms |
| T | Signal transduction mechanisms |

| | |
|---|---|
| M | Cell wall/membrane/envelope biogenesis |
| N | Cell motility |
| Z | Cytoskeleton |
| W | Extracellular structures |
| U | Intracellular trafficking, secretion, and vesicular transport |
| O | Posttranslational modification, protein turnover, chaperones |
| X | Mobilome: prophages, transposons |
| C | Energy production and conversion |
| G | Carbohydrate transport and metabolism |
| E | Amino acid transport and metabolism |
| F | Nucleotide transport and metabolism |
| H | Coenzyme transport and metabolism |
| I | Lipid transport and metabolism |
| P | Inorganic ion transport and metabolism |
| Q | Secondary metabolites biosynthesis, transport and catabolism |
| R | General function prediction only |
| S | Function unknown |
| - | No annotation |

**Supplementary Note 2** Structure elucidation of myxolutamid A and identification of the putative *mxl* BGC.

HRESIMS measurements revealed an $[M+H]^+$ peak for myxolutamid A at *m/z* 779.5400 (Supplementary Figure 22) corresponding to the molecular formula $C_{39}H_{71}N_8O_8$ (*m/z* calcd for $[M+H]^+$ 779.5389) featuring 9 double-bond equivalents (DBE). $^1$H-NMR and HSQC experiments in DMSO-$d_6$ (Supplementary Figure 18 and 20) display characteristic signals for an octapeptid with eight α-proton signals (Supplementary Table 5). Combination of COSY and HMBC correlations (Supplementary Figure 19 and 21) reveal the octapeptide to consist of five leucines as well as three alanines and the amino acid sequence was elucidated as Leu-Leu-Leu-Ala-Ala-Leu-Leu-Ala via HMBC correlations from the respective α-protons to the adjacent carbonyl groups and confirmed by the fragmentation sequence observed in the $MS^2$ spectra (Supplementary Figure 23). Noteworthy, the α-proton of Leu-1 does not show a correlation to the carbonyl of Leu-2, yet, the established connections between the other amino acids only allow ring closure in this position. Marfey's analysis (Supplementary Figure 25 and 26) was used to confirm an all-D configuration for the comprised amino acids.
*Pendulispora* MSr11367 contains 10 NRPS and 14 NRPS hybrid biosynthetic gene cluster predicted with antiSMASH 8.0[3]. Only in cluster 1.35, the predicted A-domain specificity fits the amino acid composition of myxolutamid A, which is why we propose this BGC to encode for the biosynthetic machinery involved in myxolutamid formation. The nearest Stachelhaus code[4] match of the A-domain in module 1, 3 and 4 is Leu or 3-OH Leu with their sequences showing 91, 91 and 77 % similarities, respectively. Only module 2 is predicted to incorporate alanine with the nearest Stachelhaus code match of the A-domain at 94 % similarity. Interestingly, the NRPS core gene only contains four A-domains which would require at least two modules in the BGC to act repetitively for forming the full peptide before release by the TE in module 4 (Supplementary Figure 27). The amino acid order determined by NMR and $MS^2$ indicates the *mxi*BGC as non-linear NRPS (Type C) rather than an iterative one (Type B[5]). Additionally, the esterase domain in module 3 seems to act *in trans* on the other modules to generate the all-D configuration observed in myxolutamid A, which is also consistent with the C-domains in module 2, 3, and 4 predicted as $^{D}C_{L}$ domains[6].

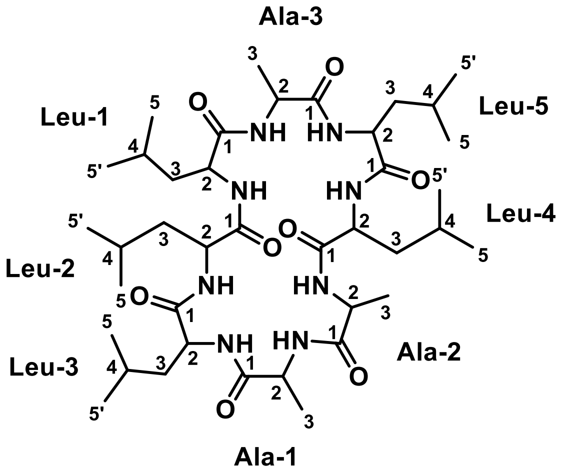

**Supplementary Figure 17** Chemical structure of myxolutamid A with atom numbering used for NMR-based structure elucidation.

**Supplementary Table 5** NMR spectroscopic data of myxolutamid A in DMSO-d6 at 500/125 MHz.

| # | δ $^{13}$C [ppm] | δ $^{1}$H [ppm], mult (*J* [Hz]) | COSY | HMBC |
|---|---|---|---|---|
| **Leu-1** | | | | |
| 1 | 175.1 | - | - | - |
| 2 | 54.7 | 4.19, m | 3, NH | 1, 3, 4 |
| 3 | 39.6 | 1.48, 1.73, m | 2, 4 | 2, 4, |
| 4 | 24.0 | 1.86, m | 3, 5’ | 3, 5’ |
| 5 | 20.5 | 0.93, d* | 3, 4, 5’ | 3, 4, 5’ |
| 5’ | 22.8 | 1.01, d (6.26) | 4, 5 | 3, 4, 5 |
| NH | - | 9.02, bd | 2 | - |
| **Leu-2** | | | | |
| 1 | 172.0 | - | - | - |
| 2 | 52.5 | 4.12, m | 3, NH | 1, Leu-3 1 |
| 3 | 39.3 | 1.48, 1.90, m | 2, 4 | 2, 4, 5, 5’ |
| 4 | 24.1 | 1.69, m | 3, 5, 5’ | 3, 5, 5’ |
| 5 | 21.4 | 0.96, d (6.56) | 3, 4, 5’ | 3, 4, 5’ |
| 5’ | 23.1 | 1.02, d (6.64) | 4, 5 | 3, 4, 5 |
| NH | - | 7.54, bd | 2 | - |
| **Leu-3** | | | | |
| 1 | 171.3 | - | - | - |
| 2 | 50.4 | 4.32, m | 3, 4, NH | 1, 3, 4, NH, Ala-1 1 |
| 3 | 38.4 | 1.69, 1.79, m | 2, 4, 5, 5’ | 2, 4, 5, 5’ |
| 4 | 22.9 | 1.66, m | 3, 5, 5’ | 2, 3, 5 |
| 5 | 20.6 | 0.91, d (6.50) | 3, 4, 5’ | 3, 4, 5’ |
| 5’ | 22.8 | 0.99, d (6.71) | 4, 5 | 3, 4, 5 |
| NH | - | 7.63, d (7.71) | 2 | Ala-1 1 |
| **Ala-1** | | | | |
| 1 | 172.7 | - | - | - |
| 2 | 51.7 | 3.89, m | 3, NH | 1, 3 |
| 3 | 15.8 | 1.38, d (7.30) | 2 | 1, 2 |
| NH | - | 8.71, bd | 2 | 1, Ala-2 1 |
| **Ala-2** | | | | |
| 1 | 173.6 | - | - | - |
| 2 | 48.5 | 4.50, m | 3, NH | 1, 3, Leu-4 1 |
| 3 | 15.5 | 1.23, d (6.33) | 2 | 1, 2 |
| NH | - | 7.40, bs | 2 | Leu-4 1 |
| **Leu-4** | | | | |
| 1 | 171.3 | - | - | - |
| 2 | 48.6 | 4.78, m | 3, NH | 1, 3 |
| 3 | 40.8 | 1.81, 1.54, m | 2, 4, 5, 5’ | 2, 4, 5, 5’ |
| 4 | 23.9 | 1.58, m | 3, 5, 5’ | - |
| 5 | 20.3 | 0.90, d (6.41) | 3, 4, 5’ | 3, 4, 5’ |
| 5’ | 22.8 | 1.01, d (6.26) | 4, 5 | 3, 4, 5 |
| NH | - | 7.14, bd (9.40) | 2 | Leu-5 1 |
| **Leu-5** | | | | |
| 1 | 171.6 | - | - | - |
| 2 | 49.6 | 4.37, m | 3 | 1, 3 |
| 3 | 39.9 | 1.69, 1.75, m | 2, 4, 5, 5’ | 2, 5, 5’ |

| | | | | |
|---|---|---|---|---|
| 4 | 23.8 | 1.81, m* | 3, 5, 5’ | - |
| 5 | 20.6 | 0.94, d* | 3, 4, 5’ | 3, 4, 5’ |
| 5’ | 23.3 | 0.99, d (5.34) | 3, 4, 5’ | 3, 4, 5 |
| NH | - | 7.59, bd (9.99) | 2 | Ala-3 1 |
| **Ala-3** | | | | |
| 1 | 173.0 | - | - | - |
| 2 | 50.8 | 4.09, m | 3, NH | 1, 3, Leu-1 1 |
| 3 | 16.1 | 1.40, d (7.50) | 2 | 1, 2 |
| NH | - | 8.74, bd | 2 | - |

* no assignment possible due to overlapping signals

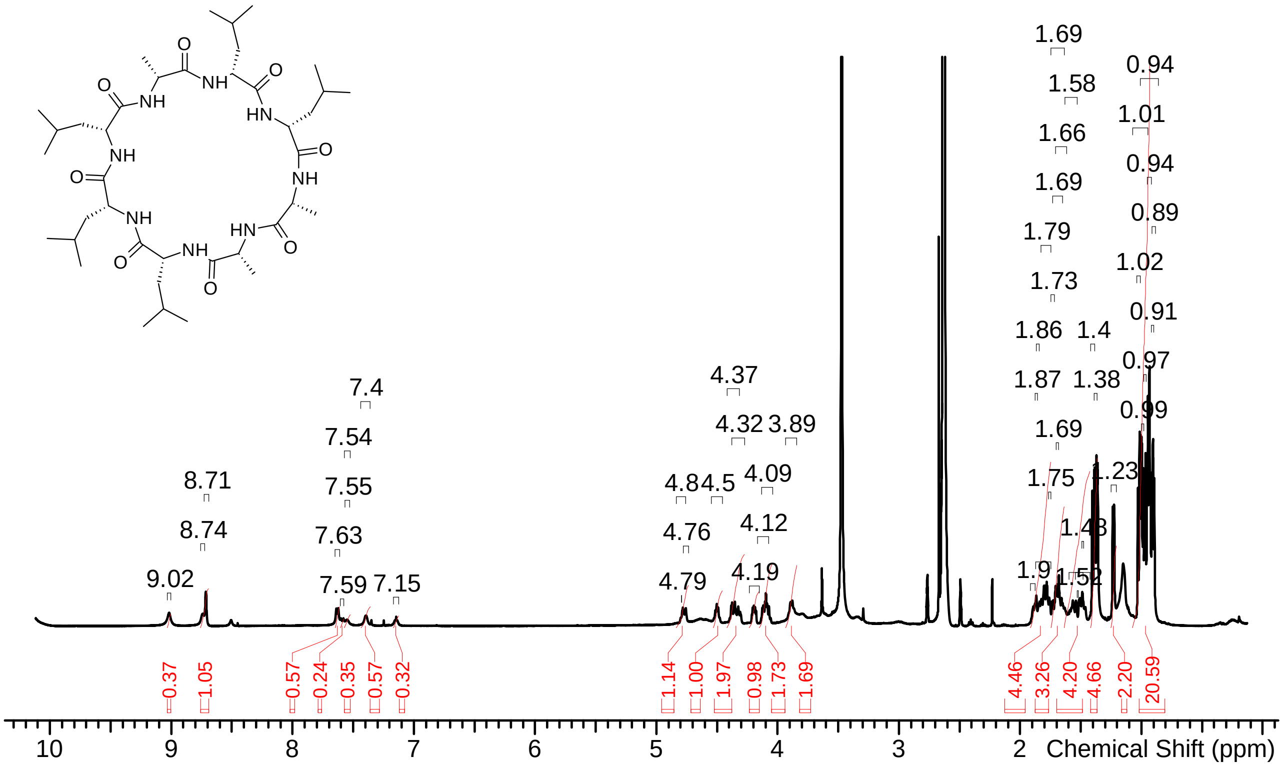

**Supplementary Figure 18** $^{1}$H-spectrum of myxolutamid A in DMSO-$d_6$ at 500 MHz.

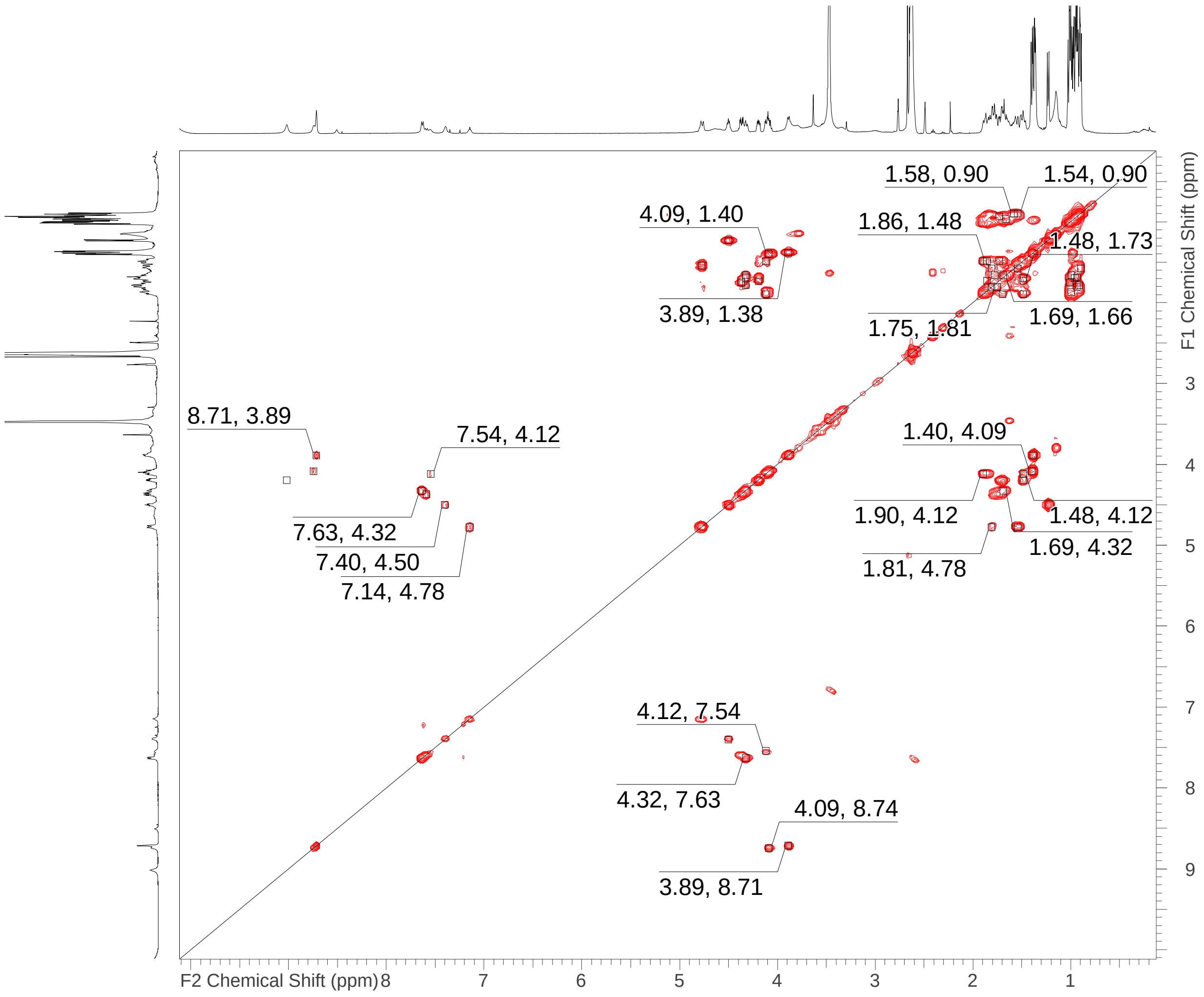

**Supplementary Figure 19** COSY-spectrum of myxolutamid A in DMSO-$d_6$ at 500 MHz.

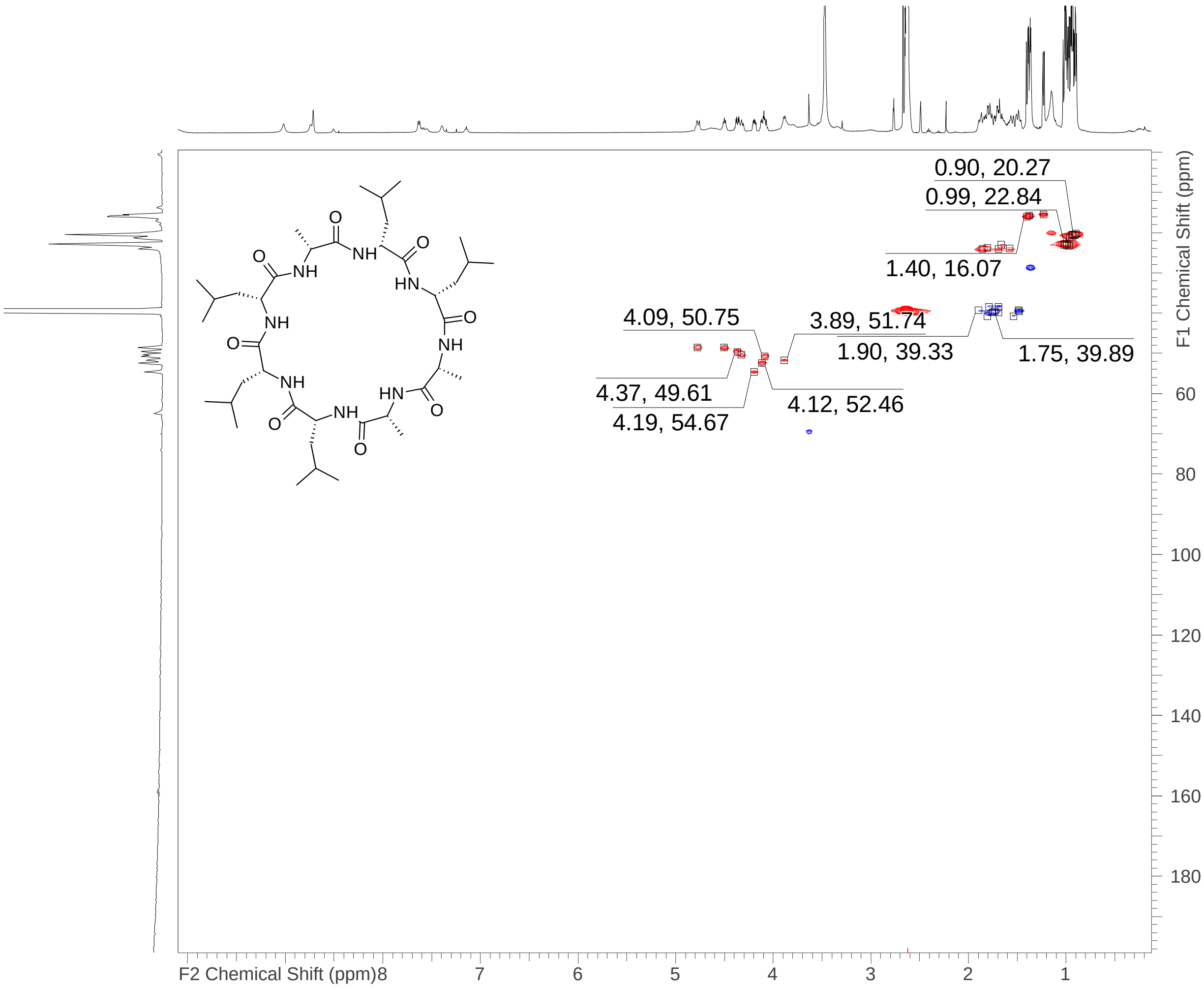

**Supplementary Figure 20** HSQC-spectrum of myxolutamid A in DMSO-$d_6$ at 500 MHz/125 MHz.

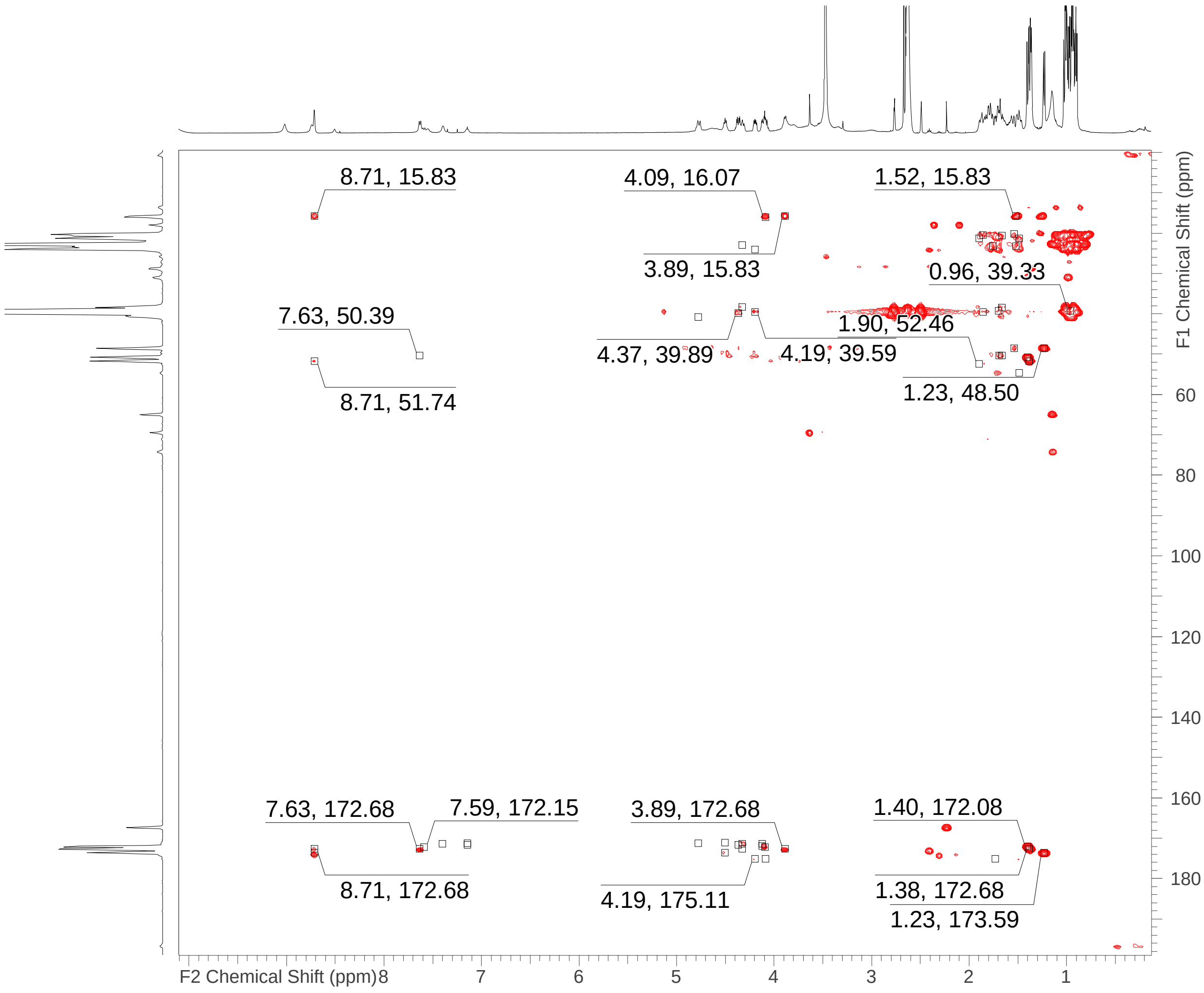

**Supplementary Figure 21** HMBC-spectrum of myxolutamid A in DMSO-d6 at 500 MHz/125 MHz.

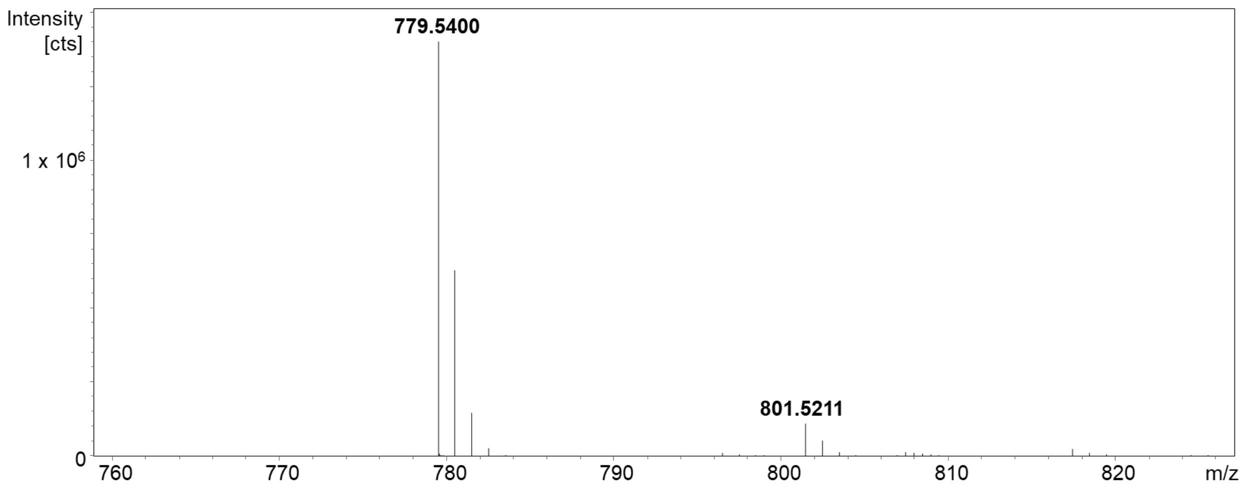

**Supplementary Figure 22** MS spectrum for myxolutamid A showing $[M+H]^+$ at *m/z* 779.5400 and $[M+Na]^+$ at *m/z* 801.5211 as most prominent ion types.

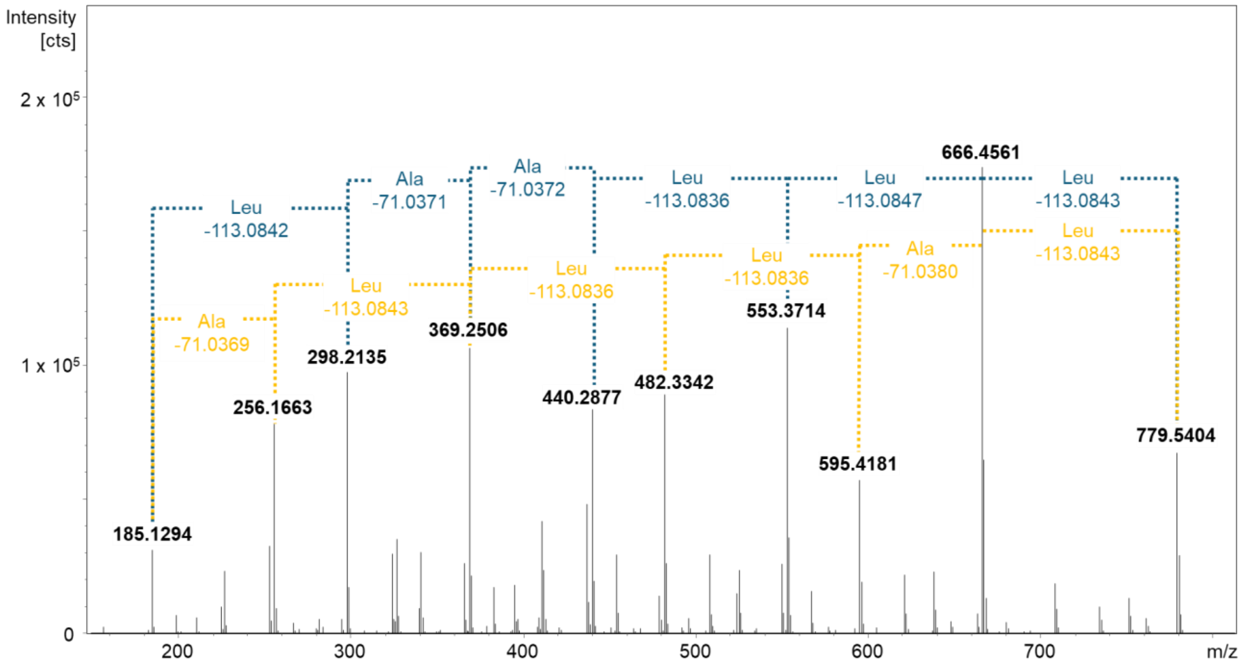

**Supplementary Figure 23** $MS^2$ fragment spectrum for myxolutamid A showing sequential losses of leucine and alanine in myxolutamid A.

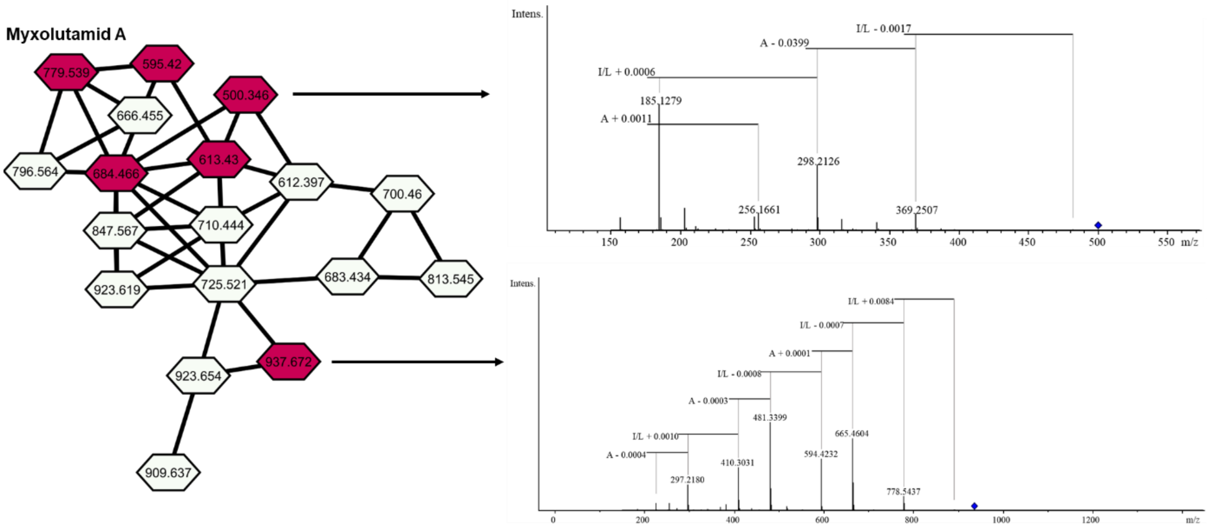

**Supplementary Figure 24** Myxolutamid molecular network generated in GNPS and $MS^2$ spectra of selected family members showing sequential losses of leucine and alanine. The myxolutamid derivative at $[M+H]^+$ shows an m/z at 595.4190 for $[M+H]^+$ in line with the sum formula $C_{30}H_{55}N_6O_6^+$ (*m/z* calcd for $[M+H]^+$ 595.4178 Δ2.1 ppm), indicating a six membered cyclic structure reduced by one leucine and one alanine in comparison to myxolutamid A. We also observe the corresponding ring opened analogues through water addition at *m/z* 684.4658 as $[M+H]^+$ with the molecular formula $C_{33}H_{62}N_7O_8^+$ (*m/z* calcd for $[M+H]^+$ 684.4654 Δ0.57 ppm) as well as at m/z 613.4297 as $[M+H]^+$ matching the sum formula $C_{30}H_{57}N_6O_7^+$ (*m/z* calcd for $[M+H]^+$ 595.4178 Δ2.1 ppm).

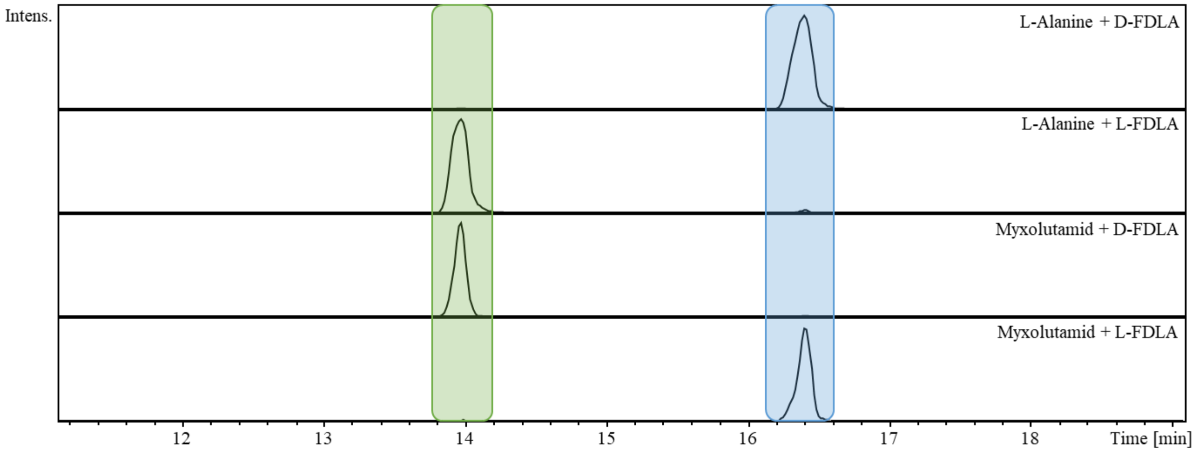

**Supplementary Figure 25** Marfey's derivatization of reference L-alanine with D-FDLA and L-FDLA (upper 2 chromatograms) and myxolutamid A with D-FDLA and L-FDLA (lower 2 chromatograms) for retention time comparison resulted in assignment of D-alanine incorporation in myxolutamid A.

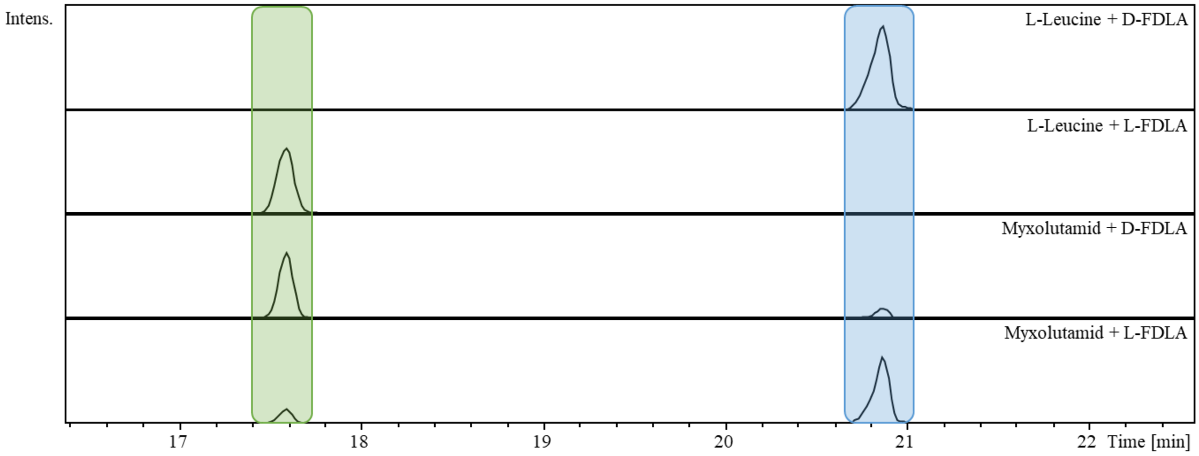

**Supplementary Figure 26** Marfey's derivatization of reference L-leucine with D-FDLA and L-FDLA (upper 2 chromatograms) and myxolutamid A with D-FDLA and L-FDLA (lower 2 chromatograms) for retention time comparison resulted in assignment of D-leucine incorporation in myxolutamid A.

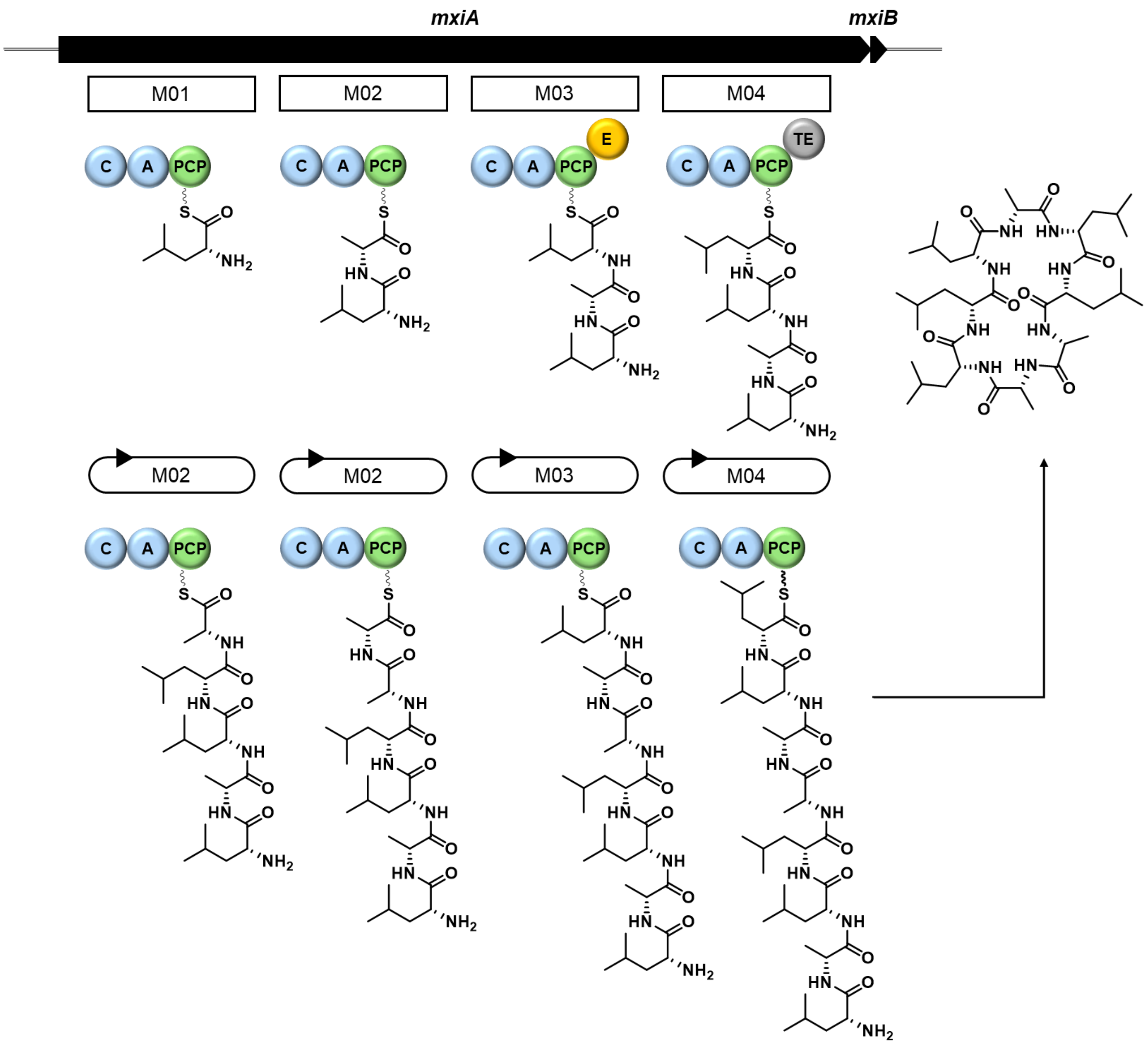

**Supplementary Figure 27** *mxl*BGC from *Pendulispora* MSr11367 putatively involved in formation of myxolutamid A. The BGC is proposed to be a non-linear NRPS with Module 2 acting repetitively twice after a first pass through the assembly line (marked with circular arrows) before the growing peptide chain is passed again to module 3 and 4 and released by the TE of module 4. M= module; C = condensation domain; A = adenylation domain; PCP = peptide carrier protein domain; E = esterase domain; TE = thioesterase domain.

**Supplementary Table 6** Open reading frames in the putative *mxl* BGC from *Pendulispora* MSr11367 with proposed function and closest homologues.

| **Name** | **Size [AA]** | **Proposed function/ homologue** | **Organism** | **Accession number of closest protein homologue** | **Sequence identity [%]** |
|---|---|---|---|---|---|
| MxiA | 4906 | ATP-dependent valine adenylase | *Brevibacillus parabrevis* | Q70LM5.1 | 39% |
| MxiB | 89 | Amino acid activation/ Protein MbtH | *Mycobacterium tuberculosis* | P59965.1 | 62% |

**Supplementary Note 3** Structure elucidation of sorangicin X and Y.

HRESIMS data of sorangicin X (Supplemental Figure 41) showed an $[M-H_2O+H]^+$ signal at *m/z* 783.4472, corresponding to a molecular sum formula of $C_{48}H_{63}O_9$ (calc. 783.4467; Δ 0.6 ppm), which contains 18 double-bond equivalents (DBEs). The chemical structure of sorangicin X was elucidated from 1D and 2D NMR data in methanol-$d_4$ (Supplemental Table 7). It differs from the reported sorangicin A [7,8] in several parts of the large macrolactone moiety (Supplemental Figure 29). Firstly, in accordance with the recently reported sorangicin P[9], sorangicin X features an additional triene-moiety at C-15 to C-20. The positions of the two additional olefinic protons H-17 and H-18 were determined by COSY correlations with the adjacent methines ($\delta_{H\text{-}16}$ 6.04, $\delta_{H\text{-}17}$ 6.32, $\delta_{H\text{-}18}$ 6.11, $\delta_{H\text{-}19}$ 6.25). Another difference in sorangicin X is the methyl-residue C-46 ($\delta_{C\text{-}46}$ 9.0, $\delta_{H\text{-}46}$ 0.8) bound to C-22, replacing the hydroxyl-residue bound to the respective carbon in sorangicin A. Moreover, sorangicin X features a ketone ($\delta_{C\text{-}25}$ 213.0) that is absent in sorangicin A. Its position was established based on HMBC correlations of the surrounding protons ($\delta_{H\text{-}23}$ 3.14, $\delta_{H\text{-}24a}$ 2.46, $\delta_{H\text{-}24a}$ 2.29, $\delta_{H\text{-}26}$ 2.14, $\delta_{H\text{-}27}$ 3.38). The configuration of *Z* and *E* double bonds was determined by their vicinal coupling constants of < 12 Hz and 14-15 Hz, respectively. This analysis yielded similar results and configurations for all double bonds that are shared with sorangicin A and point towards a *Z* configuration of the additional double bond at C-17 and C-18. The high structural similarity between sorangicin X and sorangicin A is further reinforced by a high resemblance of the respective $\delta_C$ and $\delta_H$ (Supplemental Table 8, 9). Some shift differences could be caused by a change in the ring tension of the macrolactone through the additional double bond at C-17 to C-18 as well as slight changes in the conformation caused by the substitution of two hydroxyl-residues in sorangicin A by the methyl-residue at C-22 and the ketone at C-25 in sorangicin X. The absence of the hydroxyl-residue at C-22 in sorangicin X might be also the reason for the surprisingly large differences in the chemical shifts of the sidechain, as it was hypothesized to be stabilized within the macrolactone ring by a hydrogen bond bridge between the hydroxyl-residue at C-22 and the carboxylic acid of C-1 in sorangicin A[7].

Sorangicin Y (Supplemental figure 42) shows a HRESIMS $[M-H_2O+H]^+$ signal at *m/z* 785.4637 (calc. 785.4624 Δ = 1.7 ppm), corresponding to the molecular sum formula of $C_{48}H_{65}O_9$ that has an unsaturation degree of 17 DBEs. Its structure was elucidated from 1D and 2D NMR data in methanol-$d_4$ (Supplementary Figure 30, Supplementary Table 10) and shows a high similarity to the structure of sorangicin X. However, unlike sorangicin X, it lacks the signal of the ketone C-25 at 213.0 ppm. Instead, the HSQC spectrum features a further hydroxylated methine ($\delta_{C\text{-}25}$ 71.2, $\delta_{H\text{-}25}$ 3.79), which position was established based on clear COSY correlations with the adjacent methylene and methine ($\delta_{H\text{-}24a}$ 1.67, $\delta_{H\text{-}24a}$ 1.48, $\delta_{H\text{-}26}$ 1.50).

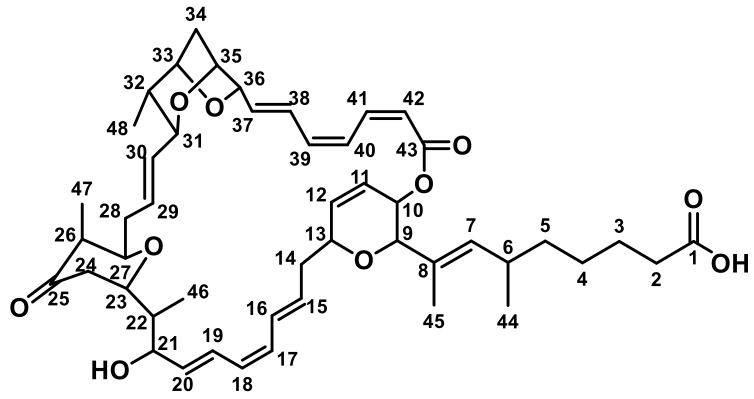

**Supplementary Figure 28** Chemical structure of sorangicin X with atom numbering used for NMR-based structure elucidation.

**Supplementary Table 7** NMR spectroscopic data of sorangicin X in methanol-$d_4$ at 500/125 MHz.

| # | δ $^{13}$C [ppm] | δ $^{1}$H [ppm], mult (*J* [Hz]) | COSY | HMBC |
|---|---|---|---|---|
| 1 | 177.6 | | | |
| 2 | 35.8 | 2.19, td (7.6, 7.6, 1.2) | 3 | 1, 3, 4 |
| 3 | 26.3 | 1.47, m | 2, 4, 5a | 1, 2, 4, 5 |
| 4 | 28.5 | 1.13, m | 3, 5a, 5b | 2, 3, 5 |
| 5a | 39.4 | 1.08, m | 3, 4 | 4, 6, 7 |
| 5b | | 0.86, m | 4, 6 | 3, 4, 6, 7 |
| 6 | 32.6 | 2.27, m | 5b, 7, 44 | 3, 7, 8 |
| 7 | 133.1 | 5.15, d (10.1) | 6, 9, 45 | 5, 6, 8, 9, 44, 45 |
| 8 | 129.8 | | | |
| 9 | 73.4 | 4.06, s | 7, 10, 45 | 5, 7, 8, 10, 13, 45 |
| 10 | 65.7 | 5.45, m | 9, 11 | 9, 11, 12, 43 |
| 11 | 124.5 | 6.09, m | 10, 12, 13 | 9, 10, 13 |
| 12 | 136.9 | 6.16, dd (9.9, 2.8) | 11, 13 | 10, 11, 13 |
| 13 | 75.0 | 4.52, m | 11, 12, 14a, 14b | 9 |
| 14a | 35.7 | 2.44, m | 13, 15 | 13, 16 |
| 14b | | 2.26, m | 13, 15, 16 | 13, 15 |
| 15 | 134.7 | 5.84, ddd (14.9, 10.4, 4.3) | 14a, 14b, 16 | 13, 14, 16, 17 |
| 16 | 133.7 | 6.04, br dd (15.2, 11.1) | 14b, 15, 17 | 14, 17, 18, 19 |
| 17 | 135.7 | 6.32, m | 16, 18 | 15, 18 |
| 18 | 130.8 | 6.11, m | 17, 19 | 16, 17, 19, 20 |
| 19 | 135.6 | 6.25, dd (15.1, 10.8) | 18, 20 | 17, 18, 21 |
| 20 | 130.8 | 5.65, br dd (15.0, 9.3) | 19, 21 | 18, 19, 21, 22 |
| 21 | 73.9 | 4.53, m | 20, 22 | 19, 20, 22, 23, 46 |
| 22 | 45.0 | 2.00, m | 21, 23, 46 | 20, 21, 23, 24, 46 |
| 23 | 81.1 | 3.14, m | 22, 24a, 24b | 21, 22, 25, 27, 46 |
| 24a | 43.2 | 2.46, m | 23, 27 | 22, 23, 25, 26 |
| 24b | | 2.29, m | 23 | 25, 26 |
| 25 | 213.0 | | | |
| 26 | 48.8 | 2.14, m | 27, 47 | 24, 25, 28, 47 |
| 27 | 78.7 | 3.38, m | 24a, 26, 28 | 23, 25, 28, 29, 47 |
| 28 | 35.7 | 2.40, m | 27, 29 | 26, 27, 29, 30 |
| 29 | 130.2 | 5.27, ddd (15.1, 10.7, 4.1) | 28, 30 | 27, 28, 30, 31 |
| 30 | 134.6 | 5.46, m | 29, 31 | 28, 29, 32 |

| | | | | |
|---|---|---|---|---|
| 31 | 80.9 | 3.80, t (9.1, 9.1) | 30, 32 | 29, 30, 32, 33, 48 |
| 32 | 41.9 | 1.41, m | 31, 33, 48 | 30, 31, 33, 34, 48 |
| 33 | 81.0 | 4.30, m | 32, 34a, 34b | 31, 34, 35, 48 |
| 34a | 39.5 | 2.03, m | 33, 35 | 33, 36 |
| 34b | | 1.92, br dd (11.5, 1.1) | 33, 35 | 32, 33, 35, 36, 37 |
| 35 | 77.4 | 4.40, m | 34a, 34b, 36 | 31, 33, 36 |
| 36 | 81.4 | 4.60, m | 35, 37, 38, 40 | |
| 37 | 135.5 | 6.30, m | 36, 38 | 35, 36, 38, 39 |
| 38 | 128.1 | 7.05, m | 36, 37, 39 | 36, 39, 40 |
| 39 | 137.9 | 6.45, br t (11.1, 11.1) | 38, 40, 42 | 37, 38, 41 |
| 40 | 127.3 | 7.26, m | 36, 39, 41, 42 | 38, 41, 42 |
| 41 | 139.3 | 7.11, m | 40, 42 | 39, 40, 42, 43 |
| 42 | 119.5 | 5.54, br d (11.4) | 39, 40, 41 | 40, 41, 43 |
| 43 | 167.9 | | | |
| 44 | 22.4 | 0.69, br d (6.6) | 6 | 3, 4, 5, 6, 7, 8 |
| 45 | 14.8 | 1.54, s | 7, 9 | 5, 7, 8, 9 |
| 46 | 9.0 | 0.88, br d (7.0) | 22 | 21, 22, 23 |
| 47 | 10.3 | 1.06, br d (7.2) | 26 | 25, 26, 27 |
| 48 | 15.4 | 0.76, br d (6.7) | 32 | 31, 33 |

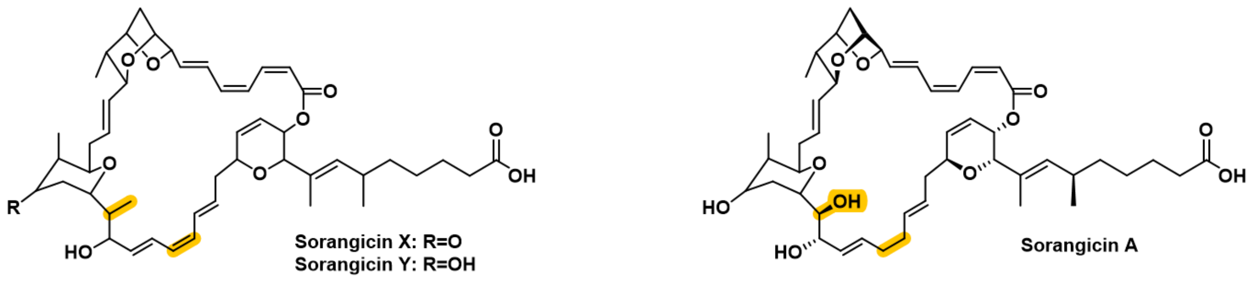

**Supplementary Figure 29** Structural comparison of sorangicins X and Y with sorangicin A. Differences are highlighted in yellow.

**Supplementary Table 8** $^{1}$H NMR comparison of sorangicin X and sorangicin A in methanol-d$_4$.

| **#** | **δ $^{1}$H [ppm] sorangicin X** | **#** | **δ $^{1}$H [ppm] sorangicin A** | **Δ (δ $^{1}$H sorangicin X - δ $^{1}$H sorangicin A)** |
|---|---|---|---|---|
| 1 | | 1 | | 0.00 |
| 2 | 2.19 | 2 | 2.3 | -0.11 |
| 3 | 1.47 | 3 | 1.62 | -0.16 |
| 4a | 1.13 | 4a | 1.36 | -0.23 |
| 4b | 1.08 | 4b | 1.3 | -0.22 |
| 5a | 1.08 | 5a | 1.42 | -0.34 |
| 5b | 0.86 | 5b | 1.25 | -0.39 |
| 6 | 2.27 | 6 | 2.43 | -0.16 |
| 7 | 5.15 | 7 | 5.34 | -0.19 |
| 8 | - | 8 | - | - |
| 9 | 4.06 | 9 | 4.28 | -0.22 |
| 10 | 5.45 | 10 | 5.35 | 0.10 |
| 11 | 6.09 | 11 | 6.05 | 0.04 |
| 12 | 6.16 | 12 | 6.17 | -0.01 |
| 13 | 4.52 | 13 | 4.43 | 0.09 |
| 14a | 2.44 | 14a | 2.43 | 0.01 |
| 14b | 2.26 | 14b | 2.17 | 0.09 |
| 15 | 5.84 | 15 | 5.58 | 0.26 |
| 16 | 6.04 | 16 | 5.58 | 0.46 |
| 17 | 6.32 | - | - | - |
| 18 | 6.11 | - | - | - |
| - | - | 17a | 2.24 | - |
| - | - | 17b | 2.14 | - |
| - | - | 18a | 2.24 | - |
| - | - | 18b | 2.17 | - |
| 19 | 6.25 | 19 | 5.79 | 0.46 |
| 20 | 5.65 | 20 | 5.64 | 0.01 |
| 21 | 4.53 | 21 | 4.19 | 0.34 |
| 22 | 2.00 | 22 | 3.52 | -1.52 |
| 23 | 3.14 | 23 | 3.73 | -0.59 |
| 24a | 2.46 | 24a | 1.76 | 0.70 |

| | | | | |
|---|---|---|---|---|
| 24b | 2.29 | 24b | 1.7 | 0.59 |
| 25 | - | 25 | 3.87 | - |
| 26 | 2.14 | 26 | 1.59 | 0.55 |
| 27 | 3.38 | 27 | 3.89 | -0.51 |
| 28a | 2.40 | 28a | 2.32 | 0.08 |
| 28b | 2.40 | 28b | 2.17 | 0.23 |
| 29 | 5.27 | 29 | 5.54 | -0.27 |
| 30 | 5.46 | 30 | 5.42 | 0.04 |
| 31 | 3.80 | 31 | 3.87 | -0.07 |
| 32 | 1.41 | 32 | 1.46 | -0.05 |
| 33 | 4.30 | 33 | 4.32 | -0.02 |
| 34a | 2.03 | 34a | 2.09 | -0.06 |
| 34b | 1.92 | 34b | 1.97 | -0.05 |
| 35 | 4.40 | 35 | 4.45 | -0.05 |
| 36 | 4.60 | 36 | 4.61 | -0.02 |
| 37 | 6.30 | 37 | 6.26 | 0.04 |
| 38 | 7.05 | 38 | 7.03 | 0.02 |
| 39 | 6.45 | 39 | 6.48 | -0.03 |
| 40 | 7.26 | 40 | 7.24 | 0.02 |
| 41 | 7.11 | 41 | 7.19 | -0.08 |
| 42 | 5.54 | 42 | 5.66 | -0.12 |
| 43 | - | 43 | - | - |
| 44 | 0.69 | 44 | 0.93 | -0.25 |
| 45 | 1.54 | 45 | 1.68 | -0.14 |
| 46 | 0.88 | - | - | - |
| 47 | 1.06 | 46 | 0.93 | 0.13 |
| 48 | 0.76 | 47 | 0.86 | -0.11 |

**Supplementary Table 9** $^{13}C$ NMR comparison of sorangicin X and sorangicin A in methanol-$d_4$.

| # | δ $^{13}C$ [ppm] sorangicin X | # | δ $^{13}C$ [ppm] sorangicin A | Δ (δ $^{13}C$ sorangicin X - δ $^{13}C$ sorangicin A) |
|---|---|---|---|---|
| 1 | 177.6 | 1 | 177.9 | -0.3 |
| 2 | 35.8 | 2 | 35.3 | 0.5 |
| 3 | 26.3 | 3 | 26.3 | 0.1 |
| 4 | 28.5 | 4 | 28.2 | 0.3 |
| 5 | 39.4 | 5 | 38.5 | 0.9 |
| 6 | 32.6 | 6 | 33.0 | -0.3 |
| 7 | 133.1 | 7 | 134.2 | -1.1 |
| 8 | 129.8 | 8 | 131.2 | -1.5 |
| 9 | 73.4 | 9 | 74.4 | -0.9 |
| 10 | 65.7 | 10 | 66.9 | -1.2 |
| 11 | 124.5 | 11 | 123.8 | 0.7 |
| 12 | 136.9 | 12 | 136.9 | 0.1 |
| 13 | 75.0 | 13 | 75.3 | -0.3 |
| 14 | 35.7 | 14 | 35.5 | 0.3 |
| 15 | 134.7 | 15 | 128.3 | 6.4 |
| 16 | 133.7 | 16 | 133.6 | 0.1 |
| 17 | 135.7 | 17 | 33.4 | 102.4 |
| 18 | 130.8 | 18 | 34.0 | 96.8 |
| 19 | 135.6 | 19 | 134.4 | 1.3 |
| 20 | 130.8 | 20 | 130.2 | 0.7 |
| 21 | 73.9 | 21 | 74.4 | -0.6 |
| 22 | 45.0 | 22 | 77.8 | -32.8 |
| 23 | 81.1 | 23 | 75.1 | 6.1 |
| 24 | 43.2 | 24 | 30.9 | 12.4 |
| 25 | 213.0 | 25 | 71.1 | 141.9 |
| 26 | 48.8 | 26 | 38.5 | 10.3 |
| 27 | 78.7 | 27 | 74.9 | 3.8 |
| 28 | 35.7 | 28 | 37.1 | -1.5 |
| 29 | 130.2 | 29 | 133.0 | -2.8 |
| 30 | 134.6 | 30 | 132.8 | 1.8 |
| 31 | 80.9 | 31 | 81.2 | -0.3 |
| 32 | 41.9 | 32 | 42.2 | -0.3 |
| 33 | 81.0 | 33 | 81.0 | 0.0 |
| 34 | 39.5 | 34 | 39.9 | -0.4 |
| 35 | 77.4 | 35 | 77.6 | -0.1 |
| 36 | 81.4 | 36 | 82.3 | -0.9 |
| 37 | 135.5 | 37 | 134.9 | 0.6 |
| 38 | 128.1 | 38 | 127.8 | 0.2 |
| 39 | 137.9 | 39 | 137.6 | 0.3 |
| 40 | 127.3 | 40 | 127.0 | 0.3 |
| 41 | 139.3 | 41 | 139.1 | 0.2 |

| | | | | |
|---|---|---|---|---|
| 42 | 119.5 | 42 | 119.7 | -0.2 |
| 43 | 167.9 | 43 | 167.7 | 0.2 |
| 44 | 22.4 | 44 | 21.7 | 0.7 |
| 45 | 14.8 | 45 | 14.3 | 0.5 |
| 46 | 9.0 | - | - | - |
| 47 | 10.3 | 46 | 10.9 | -0.6 |
| 48 | 15.4 | 47 | 15.4 | 0.0 |

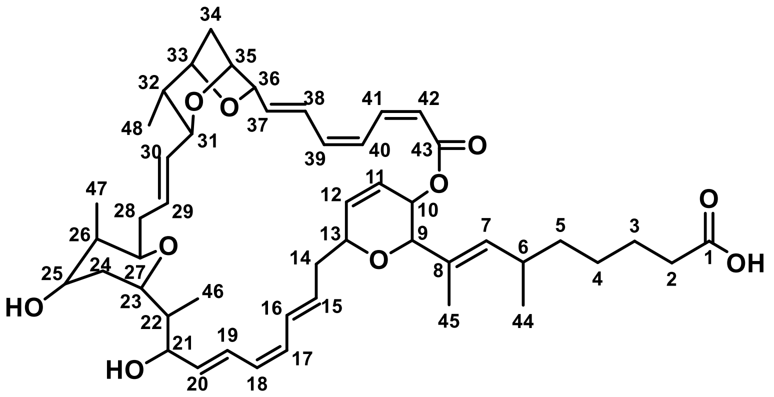

**Supplementary Figure 30** Chemical structure of sorangicin Y with atom numbering used for NMR-based structure elucidation.

**Supplementary Table 10** NMR spectroscopic data of sorangicin Y in methanol-$d_4$ at 500/125 MHz.

| # | δ $^{13}$C [ppm] | δ $^{1}$H [ppm], mult (*J* [Hz]) | COSY | HMBC |
|---|---|---|---|---|
| 1 | 178.8 | | | |
| 2 | 36.3 | 2.24, m | 3a, 3b | 1, 3, 4 |
| 3a | 26.2 | 1.62, m | 2, 4a | 1, 2, 4 |
| 3b | | 1.50, m | 2, 4b | 1, 2, 4 |
| 4a | 28.4 | 1.25, m | 3a, 5a, 5b | 2, 3, 5, 6 |
| 4b | | 1.18, m | 3b, 5a, 5b | 2, 3, 5, 6 |
| 5a | 38.8 | 1.41, m | 4a, 4b, 6 | 3, 4, 6, 7, 44 |
| 5b | | 1.00, m | 4a, 4b, 6 | 3, 6, 7, 44 |
| 6 | 32.5 | 2.32, m | 5a, 5b, 7, 44 | 4, 5, 7, 8, 44 |
| 7 | 133.7 | 5.22, br d (10.2, 1.2, 1.2) | 6, 9, 45 | 5, 6, 8, 9, 44, 45 |
| 8 | 129.5 | | | |
| 9 | 73.6 | 4.10, s | 7, 10, 45 | 7, 8, 10, 13, 45 |
| 10 | 66.1 | 5.42, dd (6.0, 1.1) | 9, 11 | 9, 11, 12, 43 |
| 11 | 124.5 | 6.09, m | 10, 12, 13 | 9, 10, 12, 13, 14 |
| 12 | 136.9 | 6.15, dd (9.8, 2.6) | 11, 13 | 10, 11 |
| 13 | 75.0 | 4.50, m | 11, 12, 14a, 14b | 9, 11, 14, 15 |
| 14a | 35.6 | 2.45, dt (14.1, 10.7, 10.7) | 13, 15 | 12, 13, 15, 16 |
| 14b | | 2.26, m | 13, 15, 16 | 15 |
| 15 | 133.9 | 5.82, ddd (14.8, 10.0, 4.5) | 14a, 14b, 16 | 13, 14, 16 |
| 16 | 131.5 | 6.12, m | 14b, 15, 17 | 13, 14, 18 |
| 17 | 135.0 | 6.28, m | 16 | 15, 16, 18 |
| 18 | 134.0 | 6.13, m | 19 | 16, 19, 20 |
| 19 | 135.3 | 6.25, m | 18, 20 | 17, 18, 20, 21 |
| 20 | 131.3 | 5.68, dd (14.9, 9.0) | 19, 21 | 16, 18, 19, 21, 22 |
| 21 | 73.8 | 4.51, m | 20, 22 | 19, 20, 22, 23, 46 |
| 22 | 44.7 | 1.78, ddd (9.9, 6.9, 4.4) | 21, 23, 46 | 20, 21, 23, 24, 46 |
| 23 | 76.8 | 3.36, m | 22, 24a, 24b | 22, 24, 25, 27, 46 |
| 24a | 33.6 | 1.67, m | 23, 24, 25 | 22, 25, 26 |
| 24b | | 1.48, m | 23, 24, 25 | 22, 23, 25 |
| 25 | 71.2 | 3.79, m | 24a, 24b, 26 | 23, 26,27, 47 |
| 26 | 37.4 | 1.50, m | 25, 27, 47 | 25, 47 |
| 27 | 74.6 | 3.72, m | 26, 28 | 23, 25, 28, 29, 47 |

| | | | | |
|---|---|---|---|---|
| 28 | 36.9 | 2.22, m | 27, 29, 30 | 26, 27, 29, 30 |
| 29 | 131.9 | 5.37, m | 28 | 28, 30, 31 |
| 30 | 133.0 | 5.36, m | 28, 31 | 28, 31, 32 |
| 31 | 81.2 | 3.81, m | 30, 32 | 29, 30, 32, 33, 48 |
| 32 | 42.0 | 1.38, m | 31, 33, 48 | 30, 31, 33, 34, 48 |
| 33 | 81.1 | 4.29, d (6.4) | 32, 34a, 34b | 31, 34, 35, 48 |
| 34a | 39.6 | 2.03, m | 33, 35 | 32, 33 |
| 34b | | 1.92, br dd (11.6, 1.3) | 33, 35 | 32, 33, 35 |
| 35 | 77.5 | 4.37, m | 34a, 34b, 36 | 31, 33, 34, 36 |
| 36 | 81.4 | 4.58, m | 35, 37, 38 | 37, 38 |
| 37 | 135.4 | 6.30, m | 36, 38 | 35, 36, 38, 39 |
| 38 | 128.6 | 7.02, m | 36, 37, 39 | 36, 39, 40 |
| 39 | 138.0 | 6.46, br t (11.2, 11.2) | 38, 40, 42 | 37, 38, 41 |
| 40 | 127.4 | 7.26, m | 39, 41, 42 | 36, 38, 41, 42, 43 |
| 41 | 139.3 | 7.11, m | 40, 42 | 39, 43 |
| 42 | 119.6 | 5.56, d (11.4) | 39, 40, 41 | 40, 41, 43 |
| 43 | 168.0 | | | |
| 44 | 22.6 | 0.76, d (6.6) | 6 | 3, 5, 6, 7 |
| 45 | 14.6 | 1.56, d (0.6) | 7, 9 | 7, 8, 9 |
| 46 | 9.7 | 0.86, br d (6.9) | 22 | 22, 23 |
| 47 | 10.7 | 0.85, br d (7.2) | 26 | 25, 26, 27 |
| 48 | 15.5 | 0.78, br d (6.7) | 32 | 31, 32, 33 |

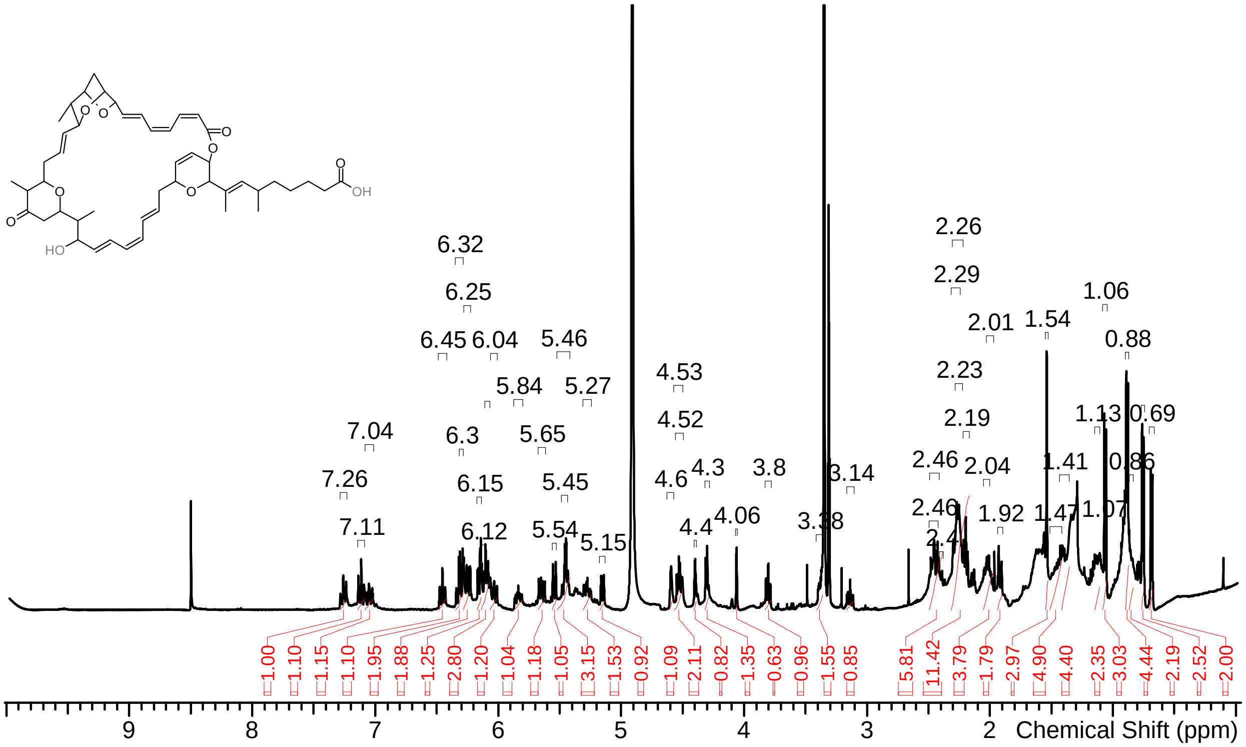

**Supplementary Figure 31** $^{1}$H-spectrum of sorangicin X in methanol-d$_4$ at 500 MHz.

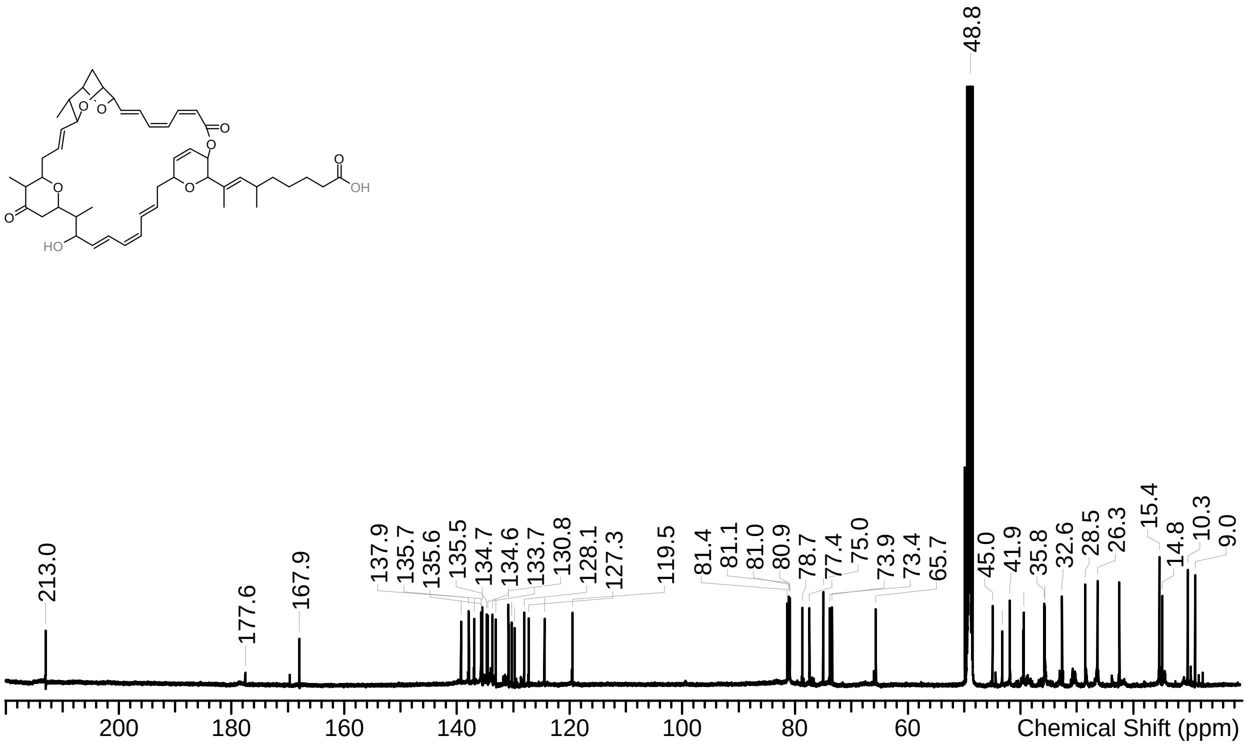

**Supplementary Figure 32** $^{13}$C-spectrum of sorangicin X in methanol-d$_4$ at 125 MHz.

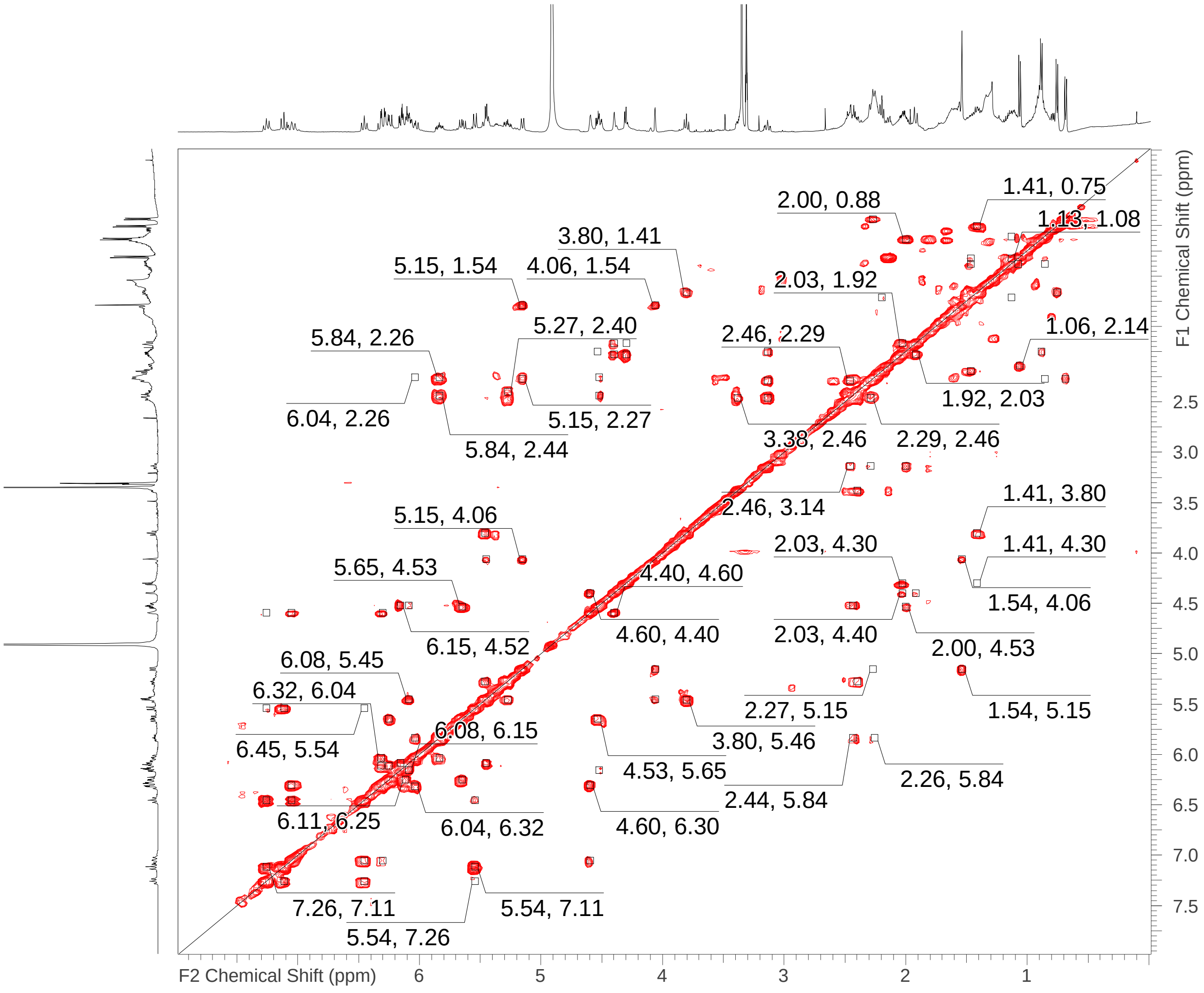

**Supplementary Figure 33** COSY spectrum of sorangicin X in methanol-$d_4$ at 500 MHz.

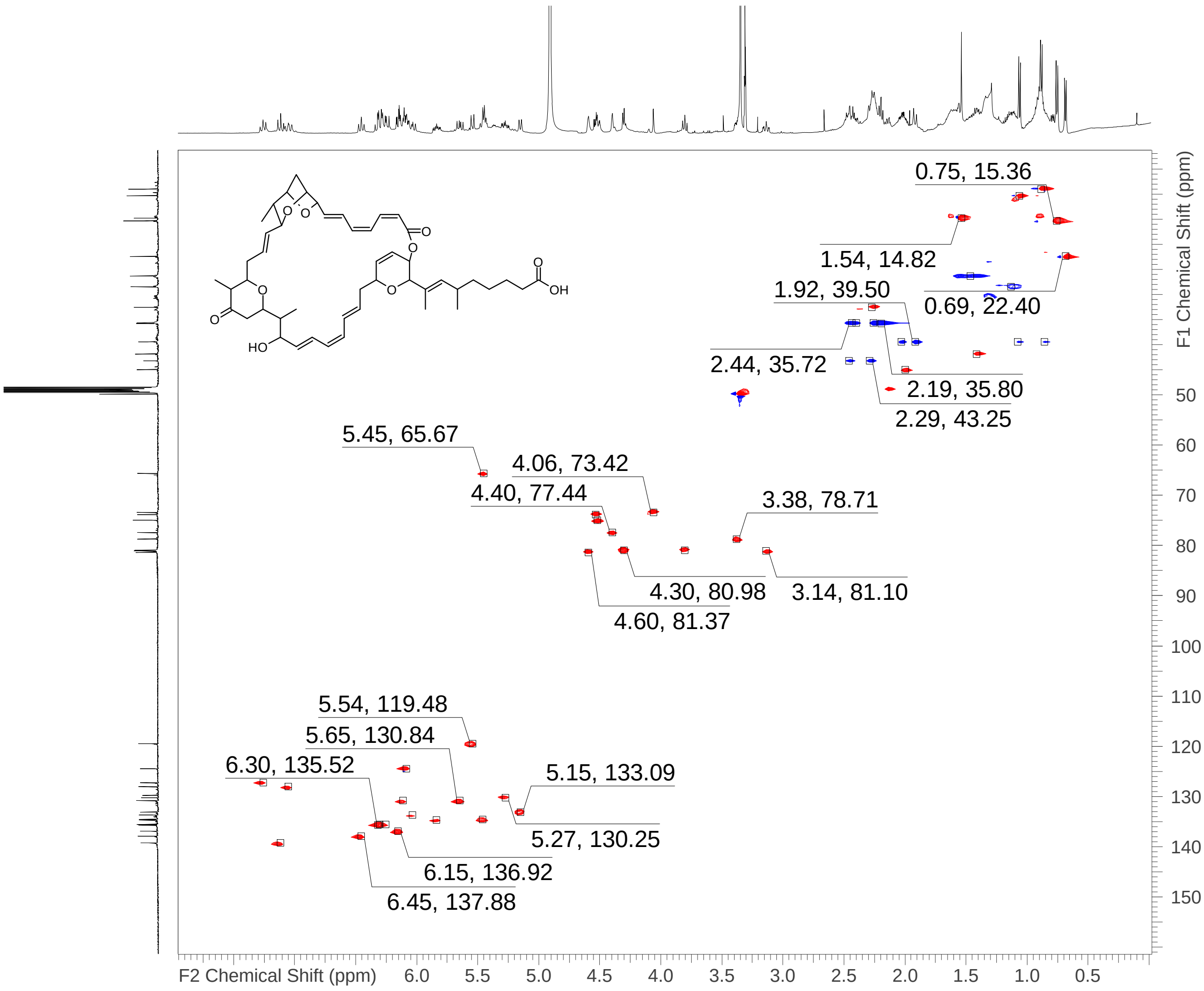

**Supplementary Figure 34** HSQC spectrum of sorangicin X in methanol-d$_4$ at 500/125 MHz.

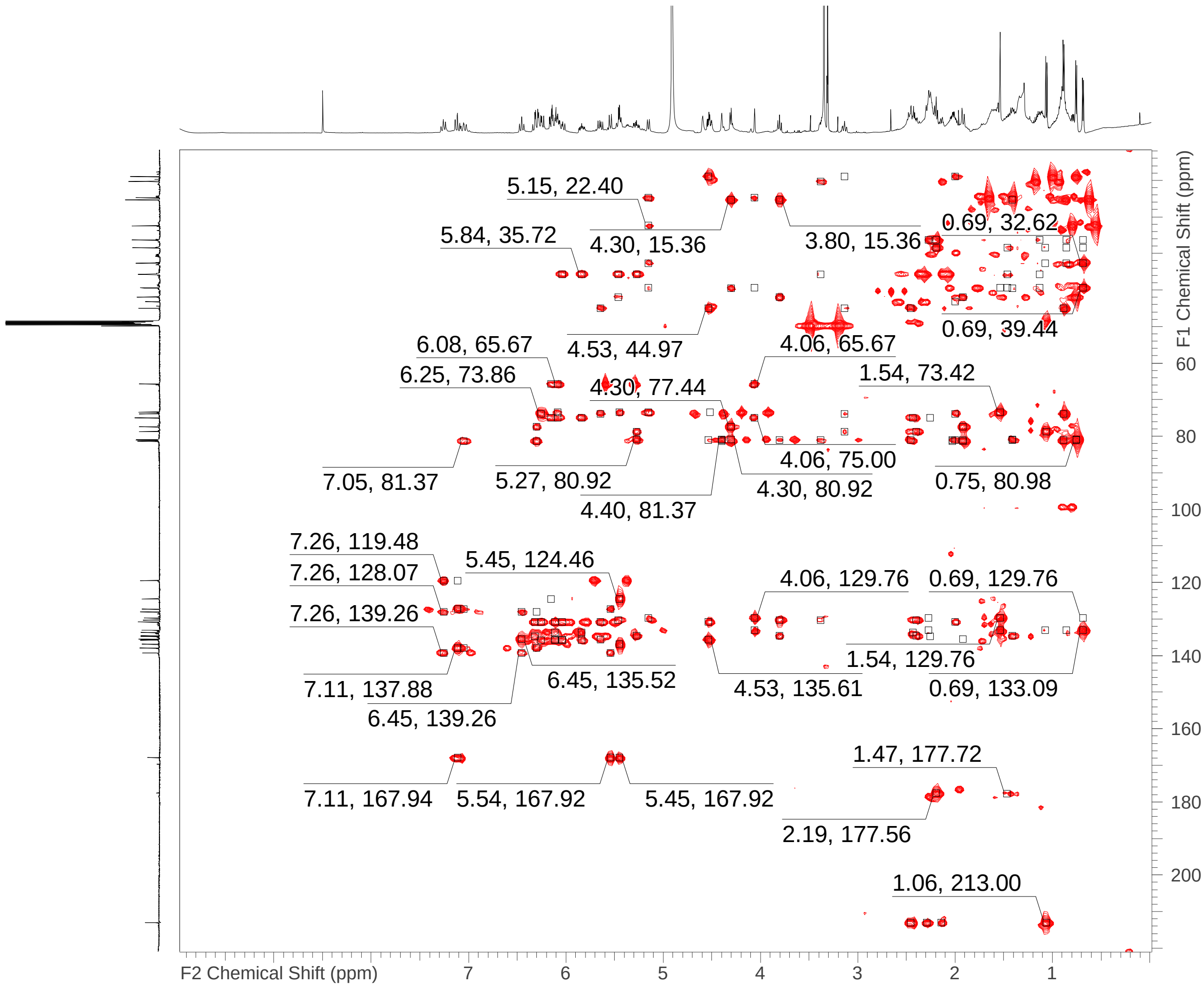

**Supplementary Figure 35** HMBC spectrum of sorangicin X in methanol-$d_4$ at 500/125 MHz.

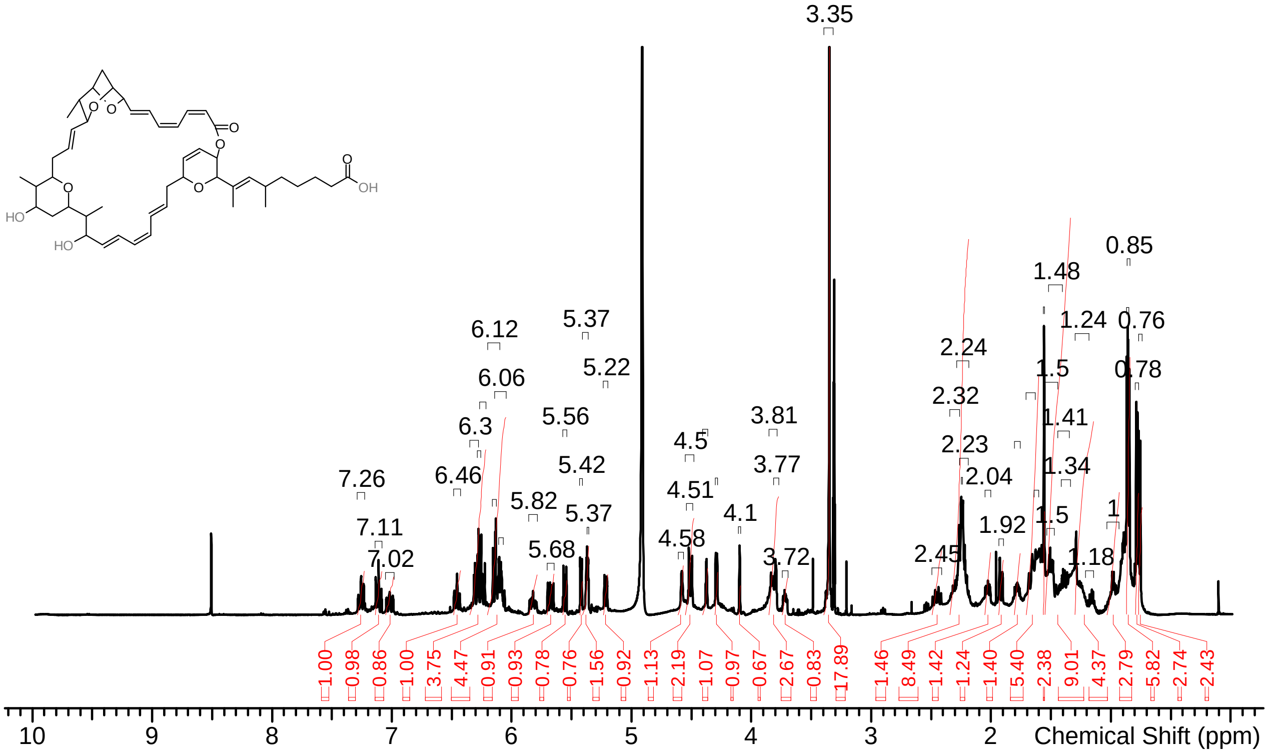

**Supplementary Figure 36** $^{1}$H-spectrum of sorangicin Y in methanol-$d_4$ at 500 MHz.

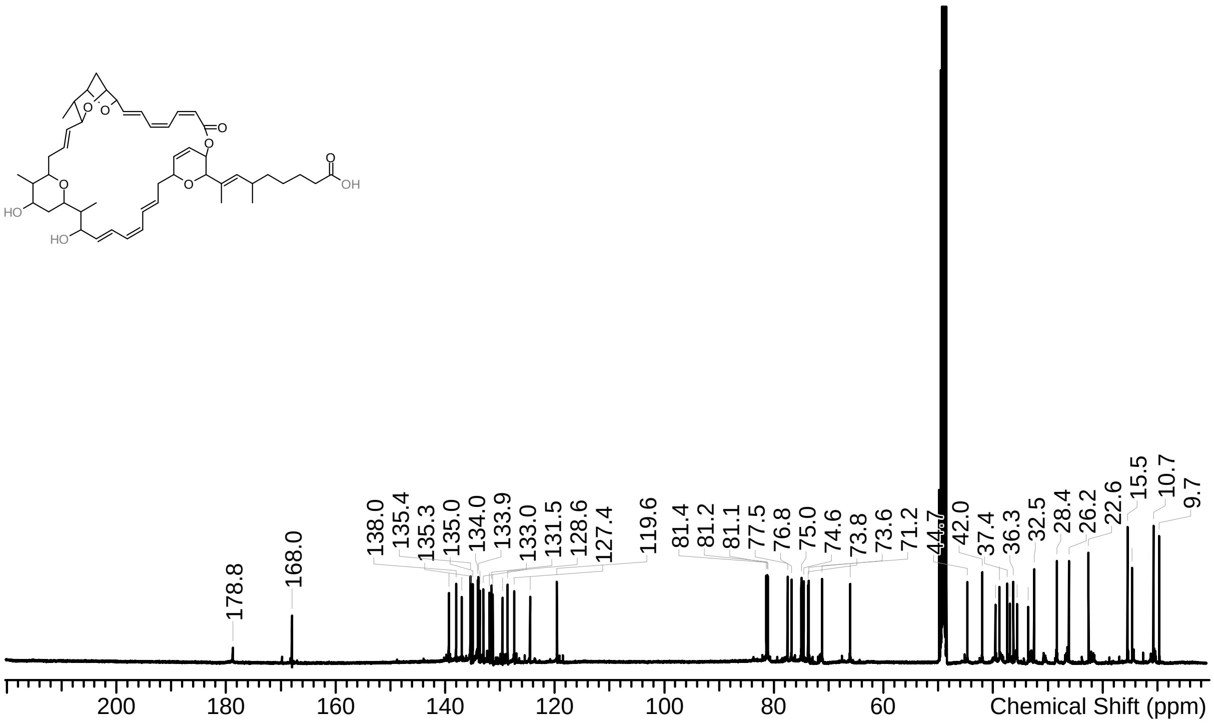

**Supplementary Figure 37** $^{13}$C-spectrum of sorangicin Y in methanol-$d_4$ at 125 MHz.

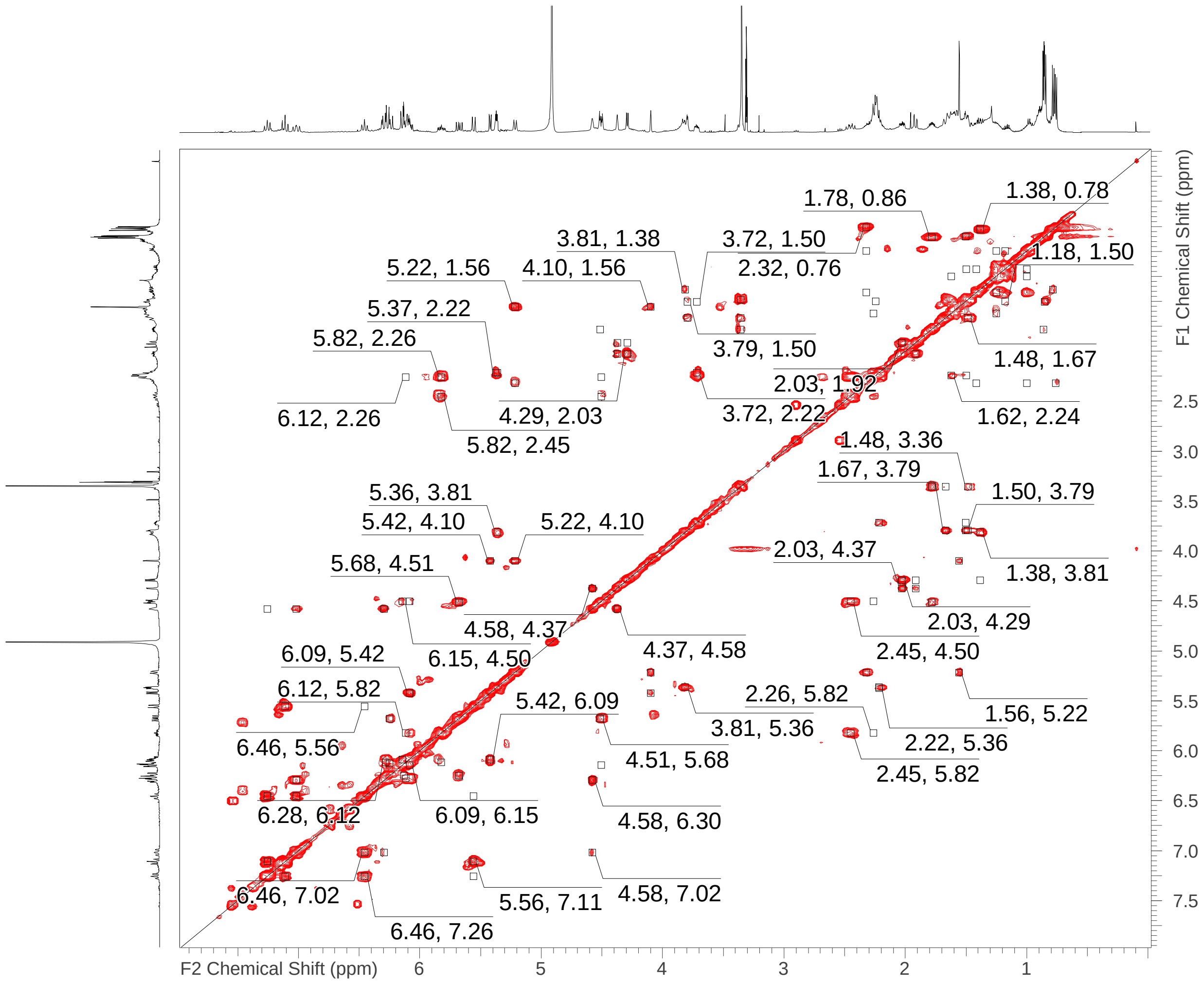

**Supplementary Figure 38** COSY spectrum of sorangicin Y in methanol-$d_4$ at 500 MHz.

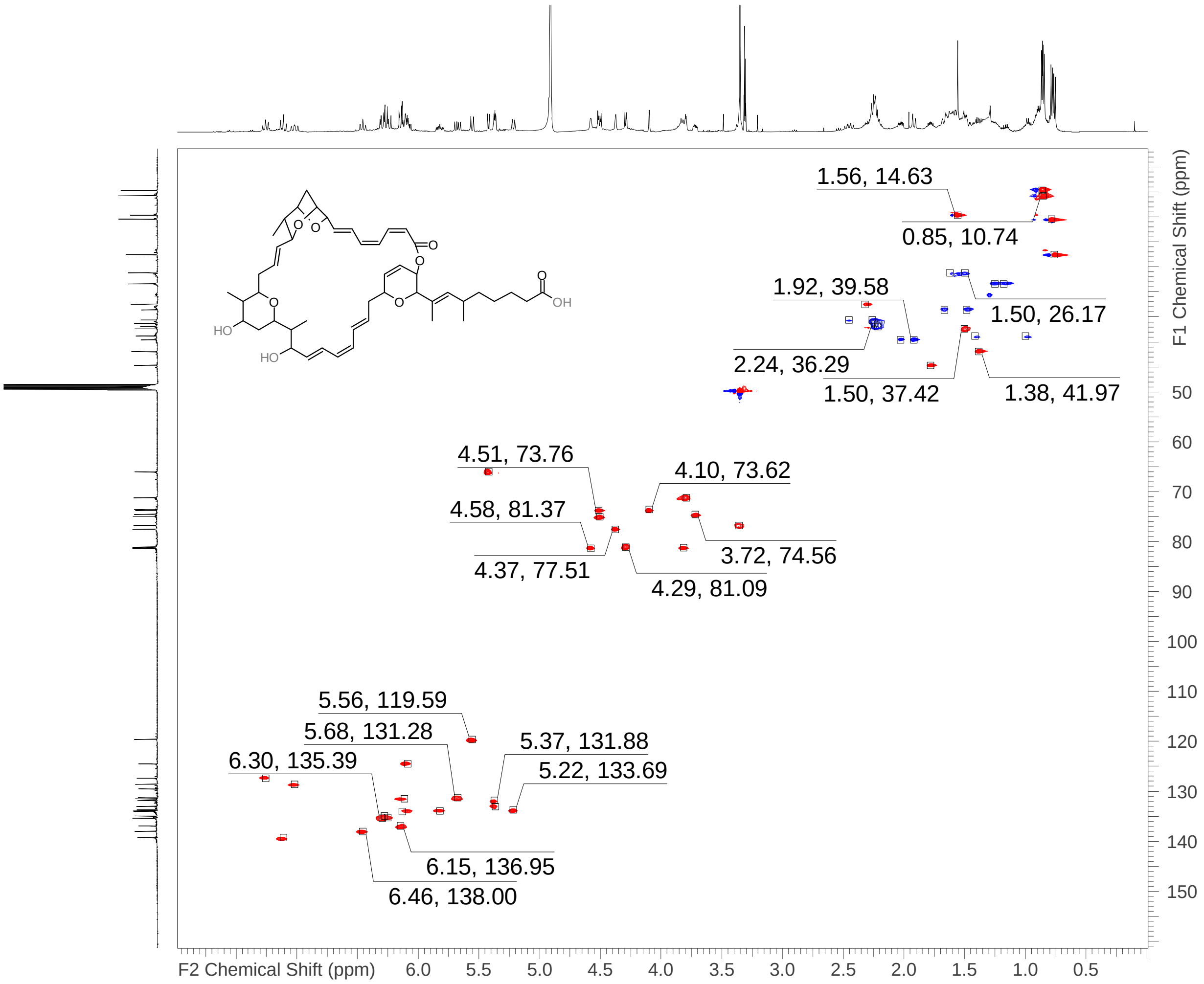

**Supplementary Figure 39** HSQC spectrum of sorangicin Y in methanol-$d_4$ at 500/125 MHz.

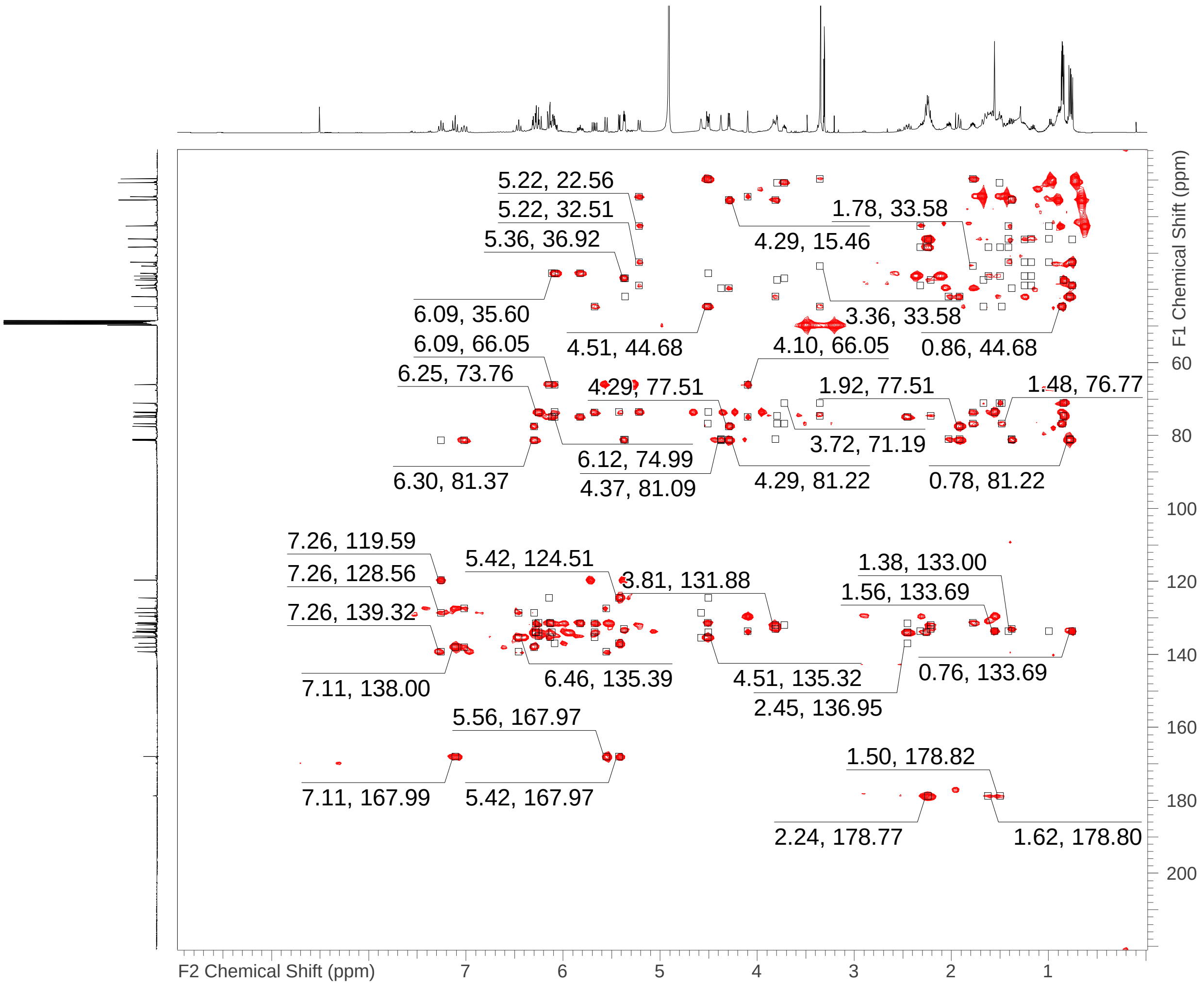

**Supplementary Figure 40** HMBC spectrum of sorangicin Y in methanol-$d_4$ at 500/125 MHz.

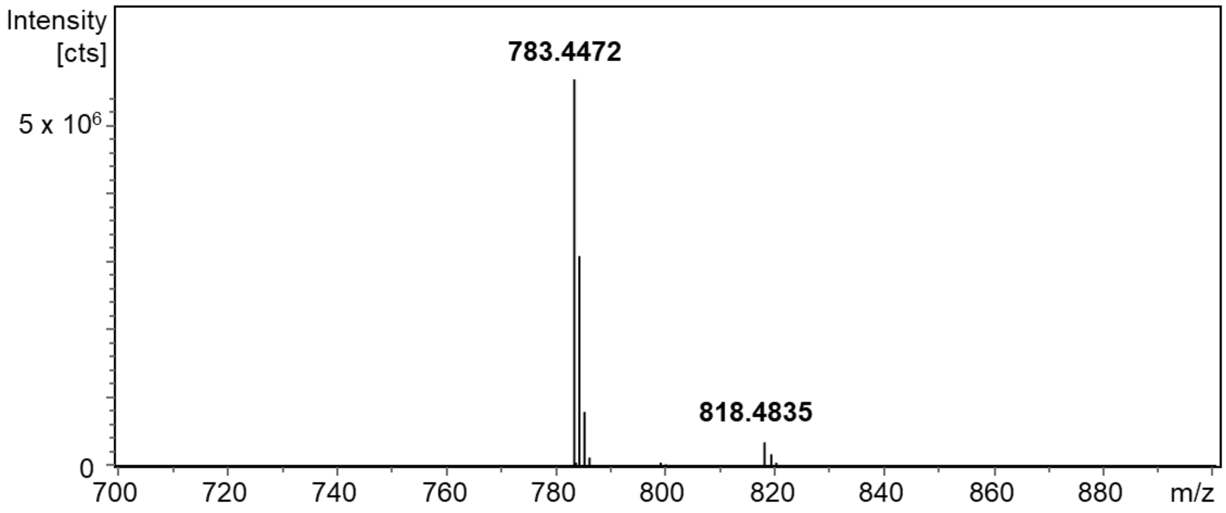

**Supplementary Figure 41** MS spectrum of sorangicin X showing $[M-H_2O+H]^+$ at *m/z* 783.4472 and $[M+NH_4]^+$ at *m/z* 818.4835 as most prominent ion types.

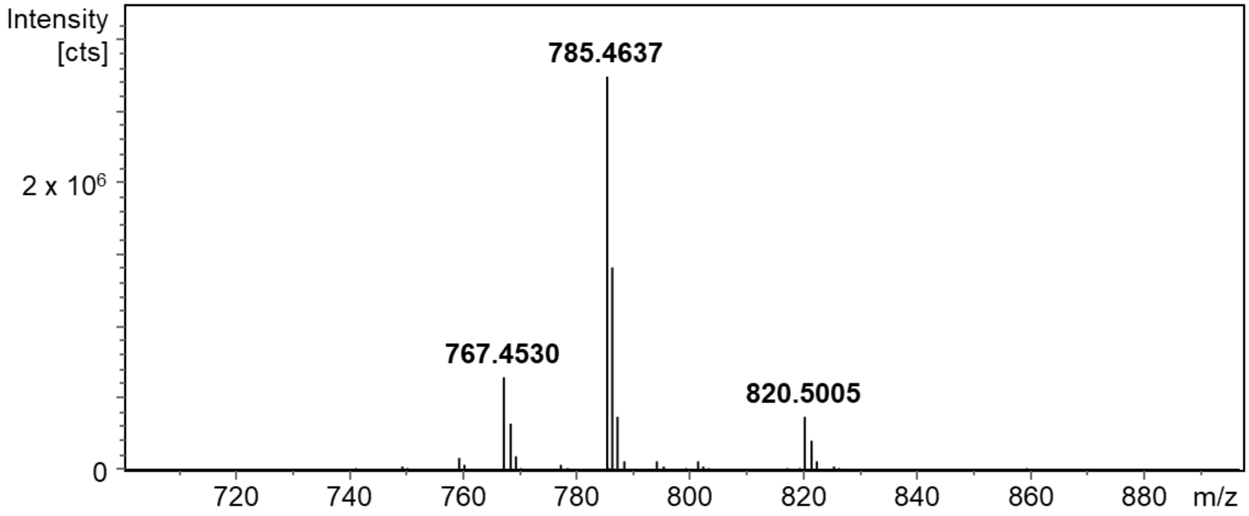

**Supplementary Figure 42** MS spectrum of sorangicin Y showing $[M-2H_2O+H]^+$ at 767.4530, $[M-H_2O+H]^+$ at *m/z* 785.4637 and $[M+NH_4]^+$ at *m/z* 820.5005 as most prominent ion types.

**Supplementary Note 4.** Analysis of the *srx* BGC and comparison to the *sor* BGC.

Comparison of the published *sor* BGC from *S. cellulosum* So ce12[10] with the *sor* BGC from So ce1525 and the new *srx* BGC observed in So ce429 showed a high level of conversation (Supplementary Figure 43A). Differing from the published *sor* BGC from So ce12, the homologs of the large biosynthetic genes *sorG* and *sorH* are condensed into one larger gene in both sorangicin BGCs observed in our studies. This observation was verified by manual analysis of the sequences of all three BGCs. The biggest differences of the *srx* BGC compared to the *sor* BGCs are the absence of a homolog of *sorK* and a much lower identity between *srxC* and *sorC*, compared to all other genes in the BGC.

A comparison of the cluster architecture of the *srx* BGC in So ce429 and the *sor* BGC in So ce1525 (Supplementary Table 11) displayed additional keto-reductase (KR) and dehydratase (DH) domains in modules 11 and 12, both encoded by *srxC*. Moreover, this comparison showed an additional DH domain in module 2, additional acyl carrier protein (ACP) domains in modules 13 and 14 as well as an O-methyl-transferase (oMT) domain in module 18 of the *srx* BGC. However, the additional domains in modules 2 and 18 of the *srx* BGC did not yield in structural differences between sorangicins X and Y compared to sorangicin A. While module 2 of the *sor* BGC needs to accommodate *in trans* acting DH, MT, KR and enoylreductase (ER) domains to yield the final configuration of the incorporated ketide[10], module 2 of the *srx* BGC already features the DH domain, but still needs the other domains to act *in trans* to explain its final product. The additional oMT domain in module 18 on the other hand is in accordance with the predicted substrate specificity of the ketosynthase (KS) domains in module 19 of the *srx* and *sor* BGC, as they are predicted to incorporate a β-methoxy-residue, according to a phylogenetic analysis[10,11]. In the matured natural product carbon C-33, which might have been methoxylated in module 18, is part of the unusual bicyclic ether moiety that is shared between sorangicin X, Y, A, and P. As the biochemical reactions that lead to this dioxabicyclooctane moiety remain elusive[9,10], the original hydroxyl- or methoxy-residue at C-33 is likely lost during the maturation of the sorangicin scaffold.

The additional KR and DH domains in modules 11 and 12 of the *srx* BGC are of greater interest, as the structural differences between sorangicin X and Y compared to sorangicin A are incorporated between modules 10 and 14 of the respective biosynthetic machineries (Supplementary Figure 43B). The additional double bond in sorangicin X and Y is incorporated by module 10 of the biosynthetic machinery and fits the observed set of reducing domains (KR and DH) of that module in the *srx* and *sor* BGCs. However, the KS domains in module 11 of both pathways share the predicted substrate specificity of a fully reduced ketide[10]. Consequently, matching the predicted substrate specificity of this KS domain as well as the observed double bond in the final natural product would entail a full reduction of the growing natural product in module 10 by an *in trans* acting enoyl reductase (ER) domain and a subsequent oxidation at a later step of the biosynthesis. Therefore, it is likely that, in contrast to the prediction, the KS domain in module 11 of the *srx* BGC accepts substrates with a double bond at the α, β position. Module 11 of the *srx* BGC differs from the respective module in the *sor* BGC as it features a KR domain, yielding the observed double bond in sorangicins X and Y. The corresponding double bond in sorangicin A can be only explained by an *in trans* acting KR domain in that module.

When describing the biosynthesis of sorangicin A, the authors discussed the discrepancy between the biosynthetically predicted incorporation of an α-methyl, β-hydroxyl moiety by module 12 and the observed α-hydroxyl, β-hydroxyl moiety that was incorporated instead. They hypothesized that in module 12 the first acyl transferase ($AT_a$) domain of the *sor* BGC incorporates hydroxymalonate instead of malonate and that the hydroxyl functionality at the α position precludes a subsequent C-methylation reaction[10]. Interestingly, sorangicin X and Y feature the α-methyl, β-hydroxyl moiety that was predicted to be incorporated by module 12 of the *sor* BGC. While both AT domains of the *srx* BGC share an amino acid similarity of 91.3% and 91.0% as

well as substrate specific fingerprints with their respective counterparts, module 12 of the *srx* BGC contains a DH domain that is absent in module 12 of the *sor* BGC. As this additional DH domain in module 12 of the *srx* BGC is the only obvious difference to the *sor* BGC, it seems possible that it might enable the methylation of the α-carbon. However, this possibility needs further experimental proof as, to the best of our knowledge, there are no reports about non-canonical DH domains that act on α-hydroxyl residues or on vicinal alcohols.
The additional tandem ACP domains in modules 13 and 14 of the *srz* BGC do not affect the structure of the molecule, such tandem ACP domains are believed to increase the efficiency of rate-limiting biosynthetic steps[12]. Therefore, the most likely explanation for the structural differences between sorangicin X and Y is an incomplete reduction of the growing polyketide by the KR domain in module 14. Next to the architecture of the *srx* BGC, the differences in their production titers point towards sorangicin Y as main product and sorangicin X as byproduct of this biosynthetic pathway (Supplementary Figure 43C).

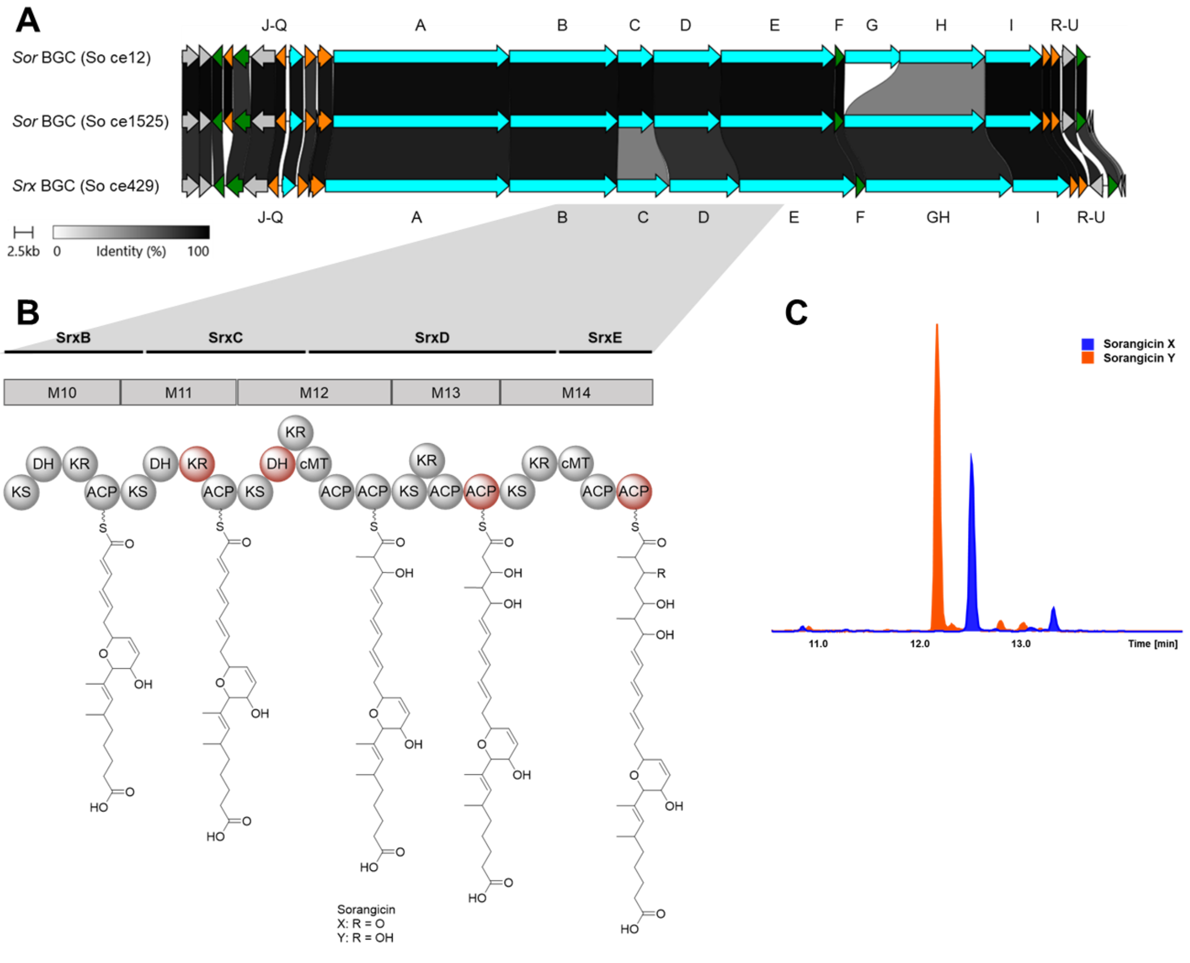

**Supplementary Figure 43** The srx BGC and the biosynthesis of sorangicins X and Y compared to the sor BGC and the biosynthesis of sorangicin A. A The published sor BGC from S. cellulosum So ce12 compared to the sor BGC from S. cellulosum So ce1525 and the srx BGC from S. cellulosum So ce429. Core biosynthetic genes are displayed in blue, additional biosynthetic genes in orange, resistance genes in green, other genes are colored in grey. B Incorporation of differences in the chemical structures of sorangicin X and Y compared to sorangicin A by the modules 10 to 14 in the srx BGC. Additional domains compared to the sor BGC are marked in red. C Relative production of sorangicin X (blue) and Y (red) in CyH medium.

**Supplementary Table 11** Comparison of the cluster architecture of the *srx* BGC in *S. cellulosum* So ce429 and the *sor* BGC in *S. cellulosum* So ce1525. Differencence compared to the published *sor* BGC in *S. cellulosum* So ce12[10] are marked bold.

| ***srx* BGC So ce429** | | | ***sor* BGC So ce1525** | | |
|---|---|---|---|---|---|
| SrxA | 1 | KS | KS | 1 | SorA |
| SrxA | 1 | DH | DH | 1 | SorA |
| SrxA | 1 | KR | KR | 1 | SorA |
| SrxA | 1 | ACP | ACP | 1 | SorA |
| SrxA | 2 | KS | KS | 2 | SorA |
| SrxA | 2 | **DH** | | 2 | SorA |
| SrxA | 2 | ACP | ACP | 2 | SorA |
| SrxA | 2 | ACP | ACP | 2 | SorA |
| SrxA | 3 | KS | KS | 3 | SorA |
| SrxA | 3 | DH | DH | 3 | SorA |
| SrxA | 3 | KR | KR | 3 | SorA |
| SrxA | 3 | cMT | cMT | 3 | SorA |
| SrxA | 3 | ACP | ACP | 3 | SorA |
| SrxA | 4 | KS | KS | 4 | SorA |
| SrxA | 4 | KR | KR | 4 | SorA |
| SrxA | 4 | ACP | ACP | 4 | SorA |
| SrxA | 5 | KS | KS | 5 | SorA |
| SrxA | 5 | ACP | ACP | 5 | SorA |
| SrxA | 6 | KS | KS | 6 | SorA |
| SrxA | 6 | KR | KR | 6 | SorA |
| SrxA | 6 | ACP | ACP | 6 | SorA |
| SrxA | 7 | KS* | KS* | 7 | SorA |
| SrxB | 7 | DH | DH | 7 | SorB |
| SrxB | 7 | ACP | ACP | 7 | SorB |
| SrxB | 7 | DH | DH | 7 | SorB |
| SrxB | 8 | KS* | KS* | 8 | SorB |
| SrxB | 8 | PS | PS | 8 | SorB |
| SrxB | 8 | ACP | ACP | 8 | SorB |
| SrxB | 9 | KS | KS | 9 | SorB |
| SrxB | 9 | DH | DH | 9 | SorB |
| SrxB | 9 | ACP | ACP | 9 | SorB |
| SrxB | 10 | KS | KS | 10 | SorB |
| SrxB | 10 | DH | DH | 10 | SorB |
| SrxB | 10 | KR | KR | 10 | SorB |
| SrxB | 10 | ACP | ACP | 10 | SorB |
| SrxB | 11 | KS | KS | 11 | SorB |
| SrxC | 11 | DH | DH | 11 | SorC |
| SrxC | 11 | **KR** | | 11 | SorC |
| SrxC | 11 | ACP | ACP | 11 | SorC |

| Gene | Module | Domain (left) | Domain (right) | Module | Gene |
|---|---|---|---|---|---|
| | 12 | KS | KS | 12 | |
| | | **DH** | | | |
| | | KR | KR | | |
| | | cMT | cMT | | |
| | | ACP | **-** | | |
| | | ACP | ACP | | |
| | 13 | KS | KS | 13 | |
| SrxD | | KR | KR | | SorD |
| | | ACP | ACP | | |
| | | **ACP** | | | |
| | 14 | KS | KS | 14 | |
| | | KR | KR | | |
| | | cMT | cMT | | |
| | | ACP | ACP | | |
| | | **ACP** | | | |
| | 15 | KS | KS | 15 | |
| | | DH | DH | | |
| | | PS | PS | | |
| | | KR | KR | | |
| SrxE | | ACP | ACP | | SorE |
| | 16 | KS | KS | 16 | |
| | | DH | DH | | |
| | | KR | KR | | |
| | | ACP | ACP | | |
| | 17 | KS | KS | 17 | |
| | | DH | DH | | |
| | | KR | KR | | |
| | | cMT | cMT | | |
| | | ACP | ACP | | |
| | 18 | KS | KS | 18 | |
| | | **oMT** | | | |
| | | ACP | ACP | | |
| | 19 | KS | KS | 19 | |
| | | DH | DH | | |
| SrxGH | | PS | PS | | SorGH |
| | | KR | KR | | |
| | | ACP | ACP | | |
| | 20 | KS | KS | 20 | |
| | | DH | DH | | |
| | | KR | KR | | |
| | | ACP | ACP | | |
| | 21 | KS | KS | 21 | |
| | | KR | KR | | |

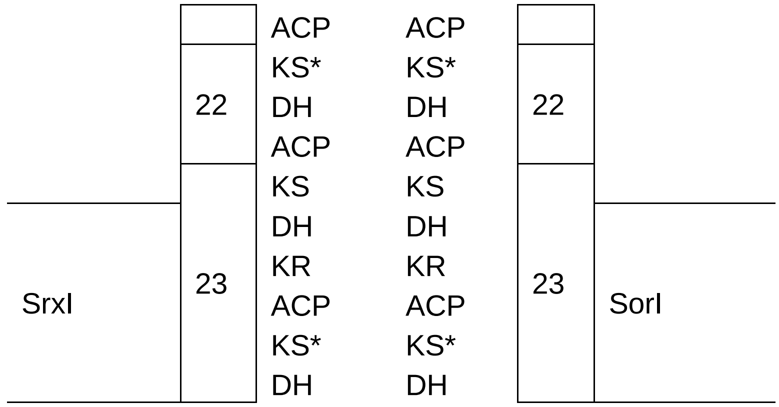
SrxI
22
23
ACP
KS*
DH
ACP
KS
DH
KR
ACP
KS*
DH
ACP
KS*
DH
ACP
KS
DH
KR
ACP
KS*
DH
22
23
SorI

**Supplementary Table 12** Biological activities of sorangicin X and Y.

| Test organism | Sorangicin A | Sorangicin X | Sorangicin Y | Rifampicin (control) |
|---|---|---|---|---|
| *E. faecium* DSM-20477 | 4 | 4 | 8 | >0.64 |
| *S. aureus* Newman | 0.2-0.32 | 0.16-2 | 0.2-0.32 | 0.006 |
| *S. aureus* Newman $RIF^R$ | >64 | >64 | >64 | >64 |
| S. *aureus* N315 (MRSA) | 0.2-0.32 | 0.05-0.08 | 0.16-0.2 | ≤0.003 |
| *A. baumannii* DSM-30008 | 32 | 8 | 16-32 | 4 |
| *A. baumannii* DSM-30008 $RIF^R$ | 64 | 16 | 64 | >64 |
| *E. coli* BW25113 (WT) | >64 | >64 | >64 | 32 |
| *E. coli* Δ*acrB* | 64 | 64 | >64 | 8 |
| *E. coli* Δ*tolC* | 64 | >64 | >64 | 16 |
| *P. aeruginosa* PA14 | >64 | >64 | >64 | >64 |

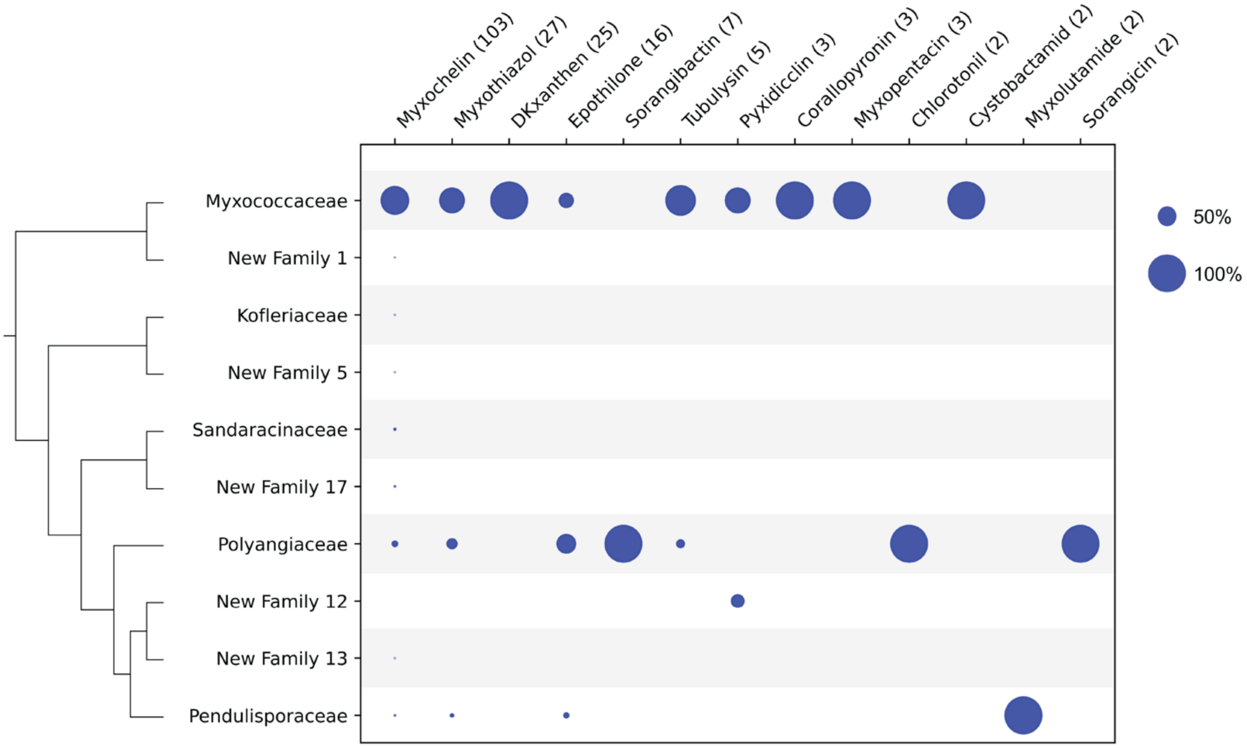

**Supplementary Figure 44** Distribution of characterized *Myxococcota* biosynthetic gene clusters (BGCs) across taxonomic groups. Dot size indicates the relative abundance of each BGC, with numerical values denoting absolute occurrences in previously reported *Myxococcota* genomes and in those presented in this study. The accompanying phylogenetic tree is resolved at the family level.

**Supplementary Note 5** Selecting the clustering thresholds for BiG-SCAPE[13] and BiG-SLiCE[14]

We evaluated the effect of different BiG-SCAPE similarity thresholds on clustering quality by comparing the resulting GCFs against compound-level similarity. First, we downloaded all BGCs from MIBiG (v3.0)[15] together with the SMILES strings of their known products. Using RDKit[16], we computed Extended Connectivity Fingerprints for all compounds and calculated pairwise Tanimoto similarities to generate a compound similarity matrix. K-means clustering was then applied to group compounds, with the optimal number of clusters (k) selected by maximizing the silhouette score. These compounds were then clustered at the optimal 'k' value and subsequently mapped back to their corresponding BGCs, forming a ground-truth clustering in BGC space. We then clustered all MIBiG BGCs at different BiG-SCAPE thresholds and compared them to the ground truth using the adjusted_rand_score and adjusted_mutual_ info_score (3 different methods) (Python library scikit-learn[17]). Both metrics measure agreement between two clustering solutions, with values close to 1 indicating high concordance, while 0 suggests no agreement. As shown in Supplementary Table 13, all metrics achieved their highest values at a BiG-SCAPE threshold of 0.5. Therefore, we adopted a threshold of 0.5 in our study for BiG-SCAPE-based clustering to identify the correct BiG-SLiCE threshold in the subsequent section

**Supplementary Table 13** Comparison of clusterings produced at different threshold values for BiG-SCAPE with the ground truth based on compound similarity for data from MIBiG.

| BiG-SCAPE threshold | adjusted_rand_score | adjusted_mutual_info_score (geometric) | adjusted_mutual_info_score (arithmetic) | adjusted_mutual_info_score (max) |
|---|---|---|---|---|
| 0.1 | 0.111 | 0.15 | 0.147 | 0.081 |
| 0.2 | 0.231 | 0.272 | 0.268 | 0.161 |
| 0.3 | 0.317 | 0.371 | 0.368 | 0.248 |
| 0.4 | 0.375 | 0.439 | 0.437 | 0.327 |
| **0.5** | **0.375** | **0.459** | **0.458** | **0.397** |
| 0.6 | 0.292 | 0.424 | 0.424 | 0.39 |
| 0.7 | 0.263 | 0.396 | 0.394 | 0.321 |
| 0.8 | 0.244 | 0.378 | 0.375 | 0.286 |
| 0.9 | 0.229 | 0.358 | 0.354 | 0.263 |
| 1 | 0.226 | 0.355 | 0.351 | 0.259 |

We compared the BGC clustering results produced by BiG-SCAPE (ground truth) to identify which BiG-SLiCE threshold yields comparable GCF assignments. For this analysis, we used BGCs from our 153 newly sequenced *Myxococcota* genomes, publicly available *Myxococcota* BGCs from the antiSMASH database, and all MIBiG BGCs. These BGCs were clustered using BiG-SCAPE (threshold 0.5; ground truth; see above paragraph) and BiG-SLiCE across thresholds ranging from 0.1 to 0.9 (step size 0.1). Clustering similarity was quantified using the adjusted_rand_score and adjusted_mutual_info score as described above. As shown in Supplementary Table 14, BiG-SLiCE at a threshold of 0.5 produced the highest agreement with the BiG-SCAPE clustering (threshold 0.5). However, the difference between thresholds 0.4 and 0.5 was minimal, and because 0.4 is the default BiG-SLiCE threshold, we adopted 0.4 for clustering in the main analyses.

**Supplementary Table 14** Comparison of clusterings produced at different threshold values for BiG-SLICE with the ground truth based on BiG-SCAPE clustering at threshold 0.5 for the same dataset.

| BiG-SLICE threshold | adjusted_rand_score | adjusted_mutual_info_score (geometric) | adjusted_mutual_info_score (arithmetic) | adjusted_mutual_info_score (max) |
|---|---|---|---|---|
| 0.1 | 0.116 | 0.202 | 0.195 | 0.110 |
| 0.2 | 0.196 | 0.362 | 0.355 | 0.223 |
| 0.3 | 0.298 | 0.526 | 0.520 | 0.378 |
| 0.4 | 0.366 | 0.600 | 0.598 | 0.501 |
| **0.5** | **0.386** | **0.619** | **0.619** | **0.592** |
| 0.6 | 0.344 | 0.602 | 0.602 | 0.562 |
| 0.7 | 0.085 | 0.490 | 0.482 | 0.370 |
| 0.8 | 0.017 | 0.374 | 0.350 | 0.232 |
| 0.9 | 0.017 | 0.339 | 0.289 | 0.176 |

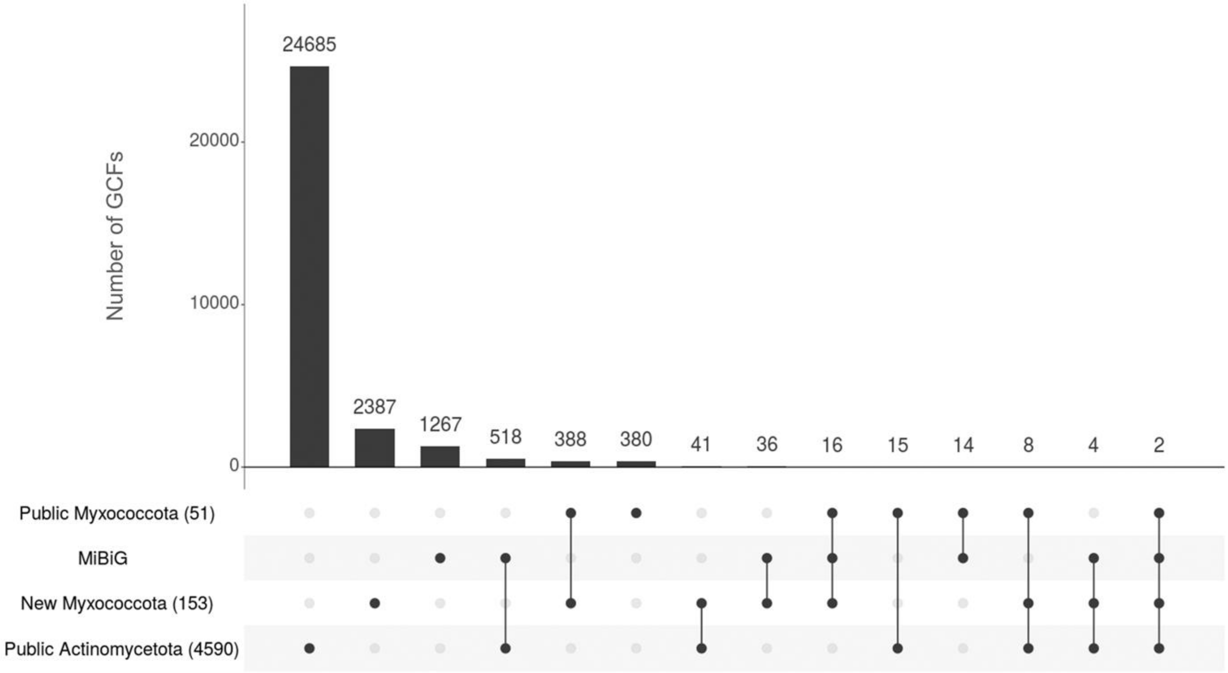

**Supplementary Figure 45** Upset plot showing the number of unique and shared GCFs between different taxa and databases considered in this study.

**Supplementary Note 6** Selection of *Actinomycetota* strains comparable to the in-house *Myxococcota* based on 16S DNA sequence.

The 16S rRNA sequences belonging to the phylum *Actinomycetota* were retrieved using NCBI BLAST[18], with three type strain sequences used as queries: Micromonospora (NR_026282), Streptomyces (NR_025663), and Actinomadura (NR_025344). This resulted in the retrieval of 16S rRNA sequences for 779 strains (having complete genomes) out of the 4,590 *Actinomycetota* strains with biosynthetic gene cluster (BGC) data available in the antiSMASH database[19]. In parallel, 16S sequences of 204 *Myxococcota* genomes were extracted from a local BLAST using Geneious Prime[20] (v2024.0.3).

The following tasks were performed using Galaxy's mothur toolsuite[21]. Both datasets were separately aligned based on the Silva SSU[22,23] (v138.1) database reference alignment and then trimmed. This resulted in 16S sequence lengths between 1,405 and 1,481 nt equivalent to the *E. coli* K-12 reference 16S sequence variable regions V1 to V9. With the processed sequences, we built separate *Myxococcota* and *Actinomycetota* distance matrices to establish sequence identities between the strains. A mothur clustering analysis (OptiClust, precision=0.001, metric=mcc, initial randomization=singleton, max number of iterations=100) was then performed using the distance matrices, which grouped each dataset's strains into families (94.1 % 16S RNA sequence identity), genera (96.3 % sequence identity), and species (98.6 % sequence identity) at 16S phylogenetic resolution.

Based on the phylogenetic diversity of 204 *Myxococcota* strains dataset, 204 out of the 779 *Actinomycetota* strains were selected to produce a dataset with matched diversity. Given its historical value in drug discovery, strains from the family *Streptomyteceae* were prioritized to match with the most represented *Myxococcota* family (*Myxococcaceae*). The remaining taxa were impartially selected in order to replicate the *Myxococcota* dataset's phylogenetic diversity.

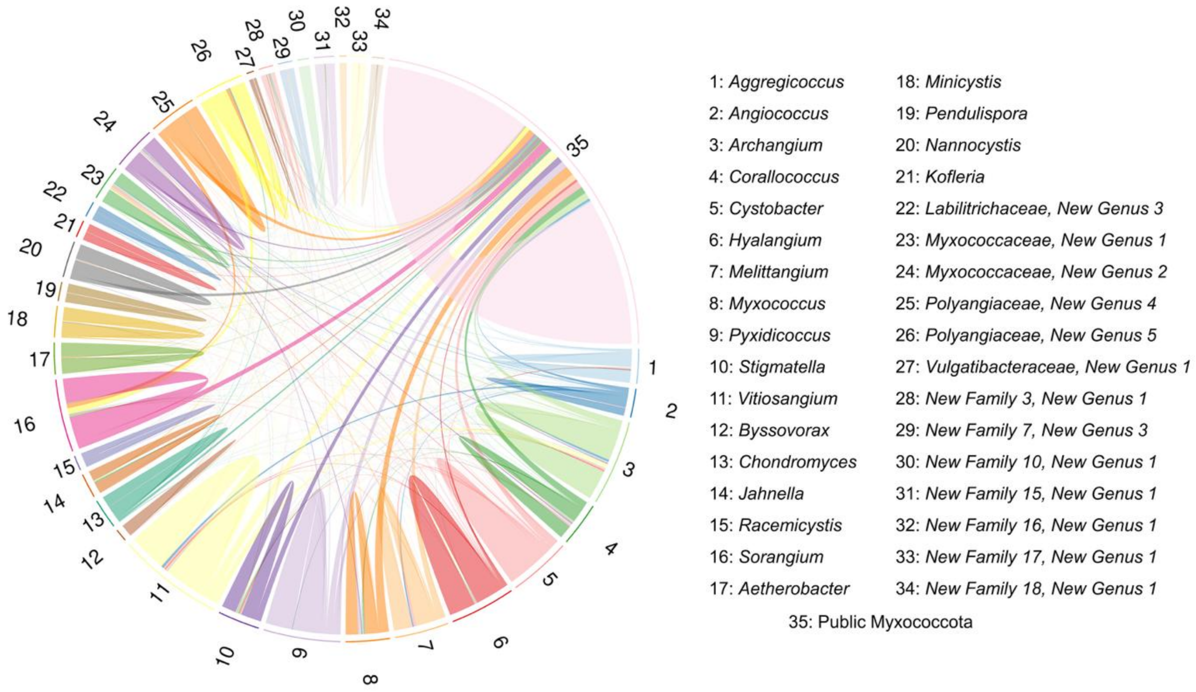

**Supplementary Figure 46** Ribbon plot showing the overlap of GCFs between different genera (comprising at least two strains) in *Myxococcota* and public *Myxococcota* genomes.

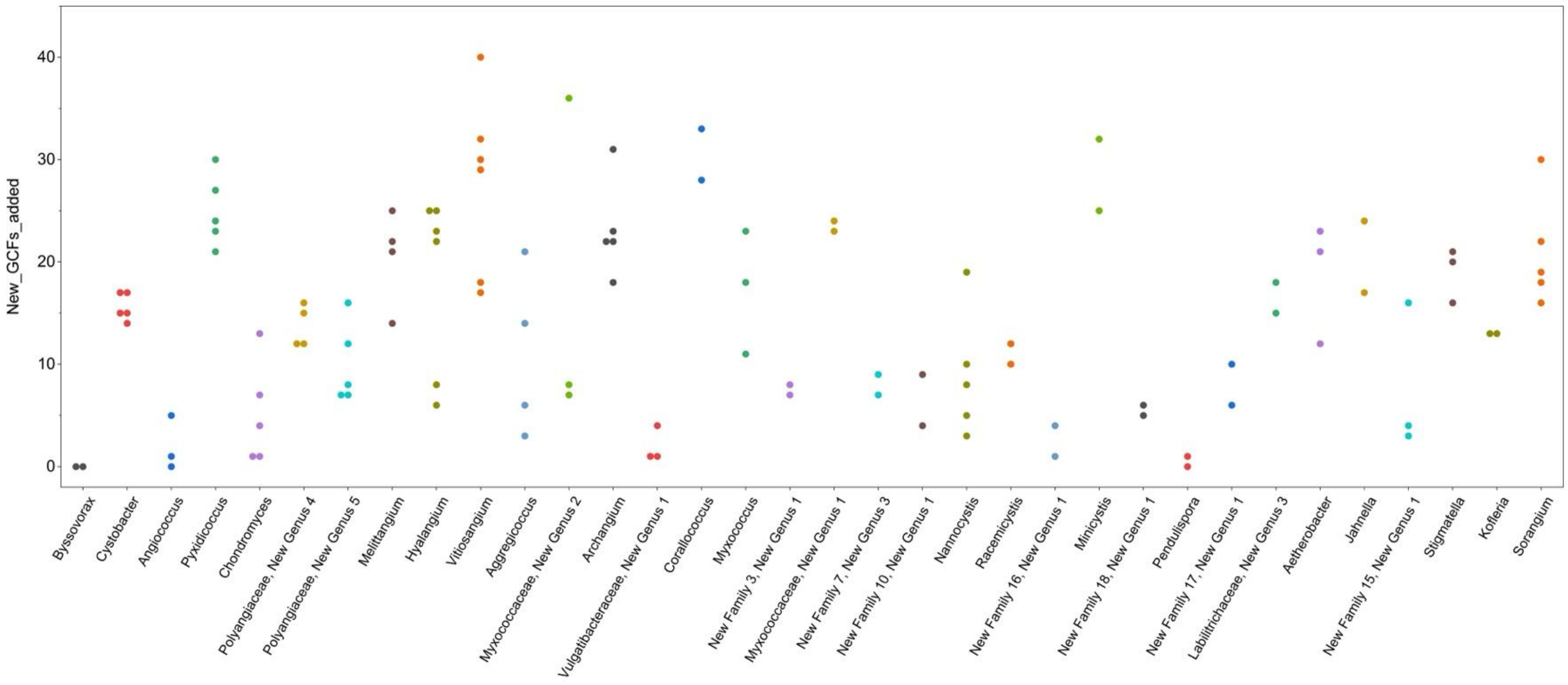

**Supplementary Figure 47** Swarm plot showing the number of new GCFs added by individual strains in each genus. Only genera with at least two strains considered.

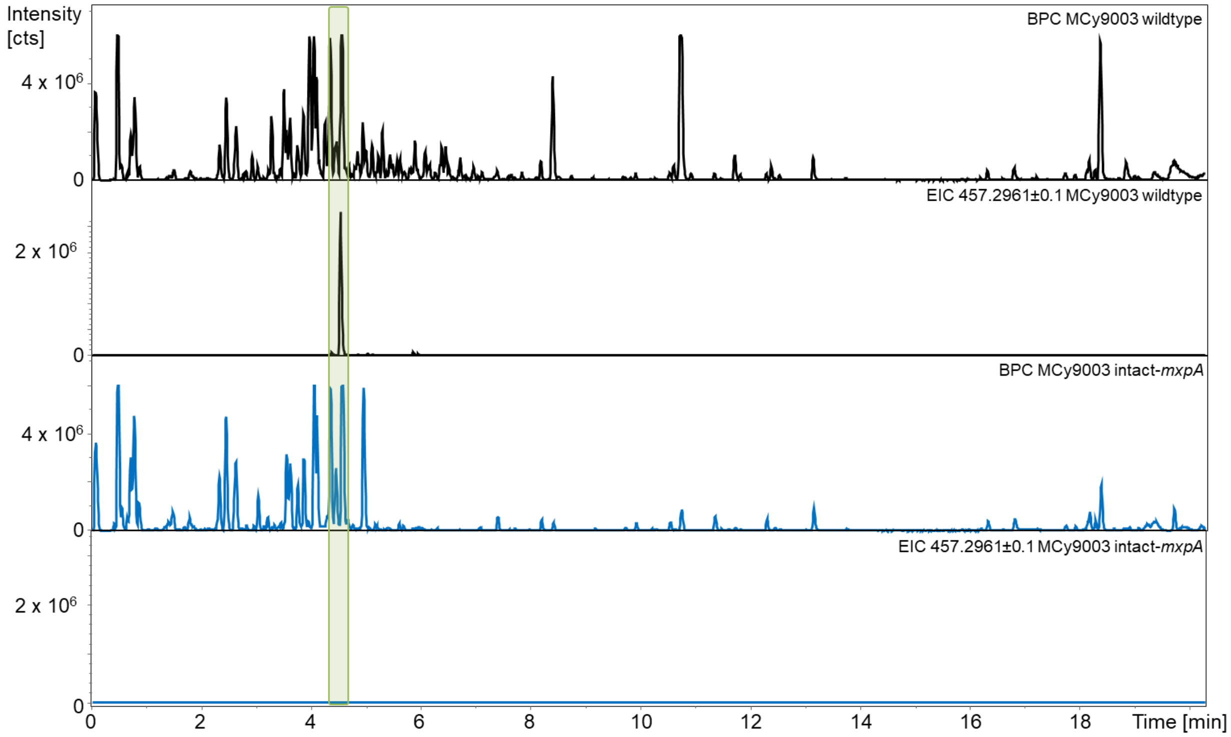

**Supplementary Figure 48** Inactivation mutant of *mxpA*. Black chromatograms depict the base peak chromatogram (BPC) from MCy9003 wildtype extracts and the extracted ion chromatogram (EIC) at 457.2961 for highlighting the myxopentacin peak observed in wildtype extracts, blue chromatograms show BPC and EIC in the inactivation MCy9003 intact-*mxpA* mutant.

**Supplementary Note 7** Structure elucidation of myxopentacin A.

HRESIMS data of myxopentacin A showed an intense signal for the double charged ion $[M+2H]^{2+}$ at *m/z* 457.297 (Supplementary Figure 54) consistent with a sum formula of $C_{46}H_{78}N_{10}O_9$ (calc. 457.29711 measured 457.29713 $\Delta$ = 0.04 ppm). Analysis of the proton spectrum showed signals characteristic for a mostly peptidic structure (Supplementary Table 15, Supplementary Figure 50). Examination of the HSQC spectrum (Supplementary Figure 52) revealed a deshielded signal at $\delta_H$ 4.09 ppm, $\delta_c$ 61.3 ppm which could be assigned as $\alpha$-position of valine using COSY and HMBC correlations (Supplementary Figure 51, 53) forming the C-terminal end of the peptide with the respective shift of the carbonyl at, $\delta_c$ 178.1 ppm indicating a free carboxylic acid. Neighboring to the assigned valine, we assigned a 5-amino-2-ethyl-8-guanidino-4-oxooctanoic acid (AGOOA) building block using characteristic chemical shifts and COSY as well as HMBC correlations: the deshielded signal at $\delta_H$ 4.43 ppm ($H\text{-}7_{AGOOA}$) indicative for an $\alpha$-position, neighbors a ketone as indicated by HMBC correlations to a carbon resonance at $\delta_C$ 208.3 ppm ($C\text{-}6_{AGOOA}$) To the same carbon resonance, HMBC correlations can be observed from two methylene protons at $\delta_H$ 3.07 and 2.58 ppm ($H\text{-}5_{AGOOA}$). These two methylene protons are part of a spin system consisting of a methine group at $\delta_H$ 2.77 ppm ($H\text{-}2_{AGOOA}$) substituted by an ethyl function and connected to the C-terminal valine via a peptide bond. Further five deshielded proton signals at $\delta_H$ 4.36, 4.28, 4.30, 4.42 and 3.67 ppm could be assigned as $\beta$-protons of five cispentacin (APCP) moieties, adjacent to the AGOOA building block. $MS^2$ fragmentation data was used to support the structure determined via 2D NMR spectroscopy (Supplementary Figure 55), showing a repetitive chain of five ACPC units, followed by the AGOOA moiety and the C-terminal valine. The configuration of the amino acids was determined by Marfey´s analysis to be L-valine and 1*(R)*,2*(S)*-amino-cyclopentylcarboxylic acid (Supplementary Figures 56 and 57).

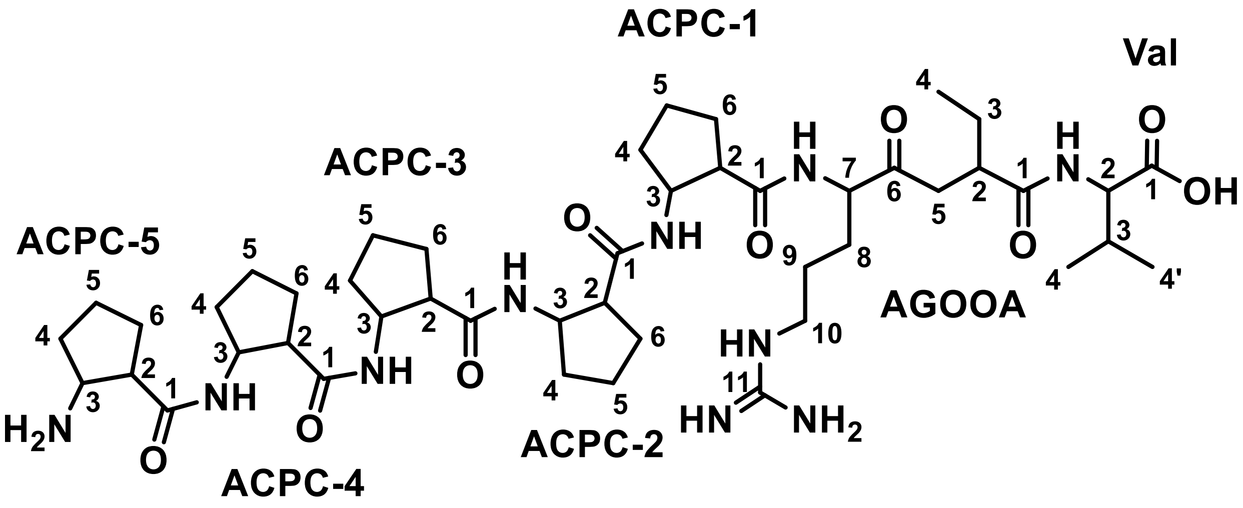

**Supplementary Figure 49** Chemical structure of myxopentacin A with atom numbering used for NMR-based structure elucidation.

**Supplementary Table 15** NMR spectroscopic data of myxopentacin A in methanol-$d_4$ at 500/125 MHz.

| # | δ $^{13}$C [ppm] | δ $^{1}$H [ppm], mult (*J* [Hz]) | COSY | HMBC |
|---|---|---|---|---|
| **Val** | | | | |
| 1 | 178.1 | - | - | - |
| 2 | 61.3 | 4.09, bd (5.6) | 3 | 1, 3, 4, 4’, AGOOA-1 |
| 3 | 32.2 | 2.08, m | 2, 4, 4’ | 2, 4, 4’ |
| 4 | 19.8 | 0.93* | 3, 4’ | 2, 3, 4’ |
| 4’ | 18.3 | 0.92* | 3, 4 | 2, 3, 4 |
| **AGOOA** | | | | |
| 1 | 176.3 | - | - | - |
| 2 | 44.1 | 2.77* | 3, 5 | 3, 4 |
| 3 | 26.7 | 1.64, 1.51* | 2, 4 | 1, 2, 3, 4, 5 |
| 4 | 11.7 | 0.96, t (7.5) | 3 | 2, 3 |
| 5 | 42.6 | 3.07, dd (18.3, 10.9), 2.58, dd (18.3, 3.6) | 2 | 1, 2, 3, 6 |
| 6 | 208.3 | - | - | - |
| 7 | 58.4 | 4.43* | 8 | 6, 8, 9, ACPC-1 |
| 8 | 27.6 | 1.58, 1.85* | 7, 9 | 7, 9 |
| 9 | 25.1 | 1.49* | 8, 10 | - |
| 10 | 41.6 | 3.17, bt (6.5) | 9 | 7, 8, 11 |
| 11 | 158.3 | - | - | - |
| **ACPC-1** | | | | |
| 1 | 175.2 | - | - | - |
| 2 | 49.1 | 2.91, bdd (15.4, 8.0) | 3, 6 | 1, 3, 6 |

| | | | | |
|---|---|---|---|---|
| 3 | 53.6 | 4.36, bdd (13.4, 6.5) | 2, 4 | 2, 4, 5, 6, ACPC-2 |
| 4 | 32.9 | 1.65, 1.88* | 3, 5 | 3, 5, 6 |
| 5 | 23.4 | 1.56, 1.86* | 4, 6 | 2, 4, 6 |
| 6 | 28.5 | 1.82, 1.97* | 5, 2 | 2, 3, 5 |
| **ACPC-2** | | | | |
| 1 | 174.7 | - | - | - |
| 2 | 49.1 | 2.83* | 3, 6 | 1, 3, 4, 5, 6 |
| 3 | 53.6 | 4.28* | 2, 4 | 1, 2, 6, ACPC-3 |
| 4 | 32.9 | 1.65, 1.92* | 3, 5 | 3, 5 |
| 5 | 23.4 | 1.56, .1.86* | 4, 6 | 2, 4, 6 |
| 6 | 28.5 | 1.82, 1.97* | 5, 2 | 1, 2, 3, 5 |
| **ACPC-3** | | | | |
| 1 | 174.5 | - | - | - |
| 2 | 49.1 | 2.83* | 3, 6 | 1, 3, 4, 5, 6 |
| 3 | 53.6 | 4.30* | 2, 4 | 1, 2, 4, ACPC-4 |
| 4 | 32.9 | 1.65, 1.92* | 3, 5 | 3, 5 |
| 5 | 23.4 | 1.56, .1.86* | 4, 6 | 2, 4, 6 |
| 6 | 28.5 | 1.82, 1.97* | 5, 2 | 1, 2, 3, 5 |
| **ACPC-4** | | | | |
| 1 | 174.5 | - | - | - |
| 2 | 49.1 | 2.83* | 3, 6 | 1, 3, 4, 5, 6 |
| 3 | 53.5 | 4.42, bdd (13.4, 7.4) | 2, 4 | 2, 4, ACPC-5 |
| 4 | 32.9 | 1.65, 1.92* | 3, 5 | 3, 5 |
| 5 | 23.4 | 1.56, .1.86* | 4, 6 | 2, 4, 6 |
| 6 | 28.5 | 1.82, 1.97* | 5, 2 | 2, 3, 5 |
| **ACPC-5** | | | | |
| 1 | 174.7 | - | - | - |
| 2 | 46.5 | 2.84* | 3, 6 | 1, 2, 6 |
| 3 | 54.4 | 3.67, bdd (11.0, 6.6) | 2, 4 | 2, 4, 5, 6 |
| 4 | 31.8 | 1.80, 2.10* | 3, 5 | 1, 3, 5 |
| 5 | 22.6 | 1.74, 1.91* | 4, 6 | 4, 6 |
| 6 | 29.6 | 1.87, 2.04* | | 2, 5 |

* no assignment possible due to overlapping signals

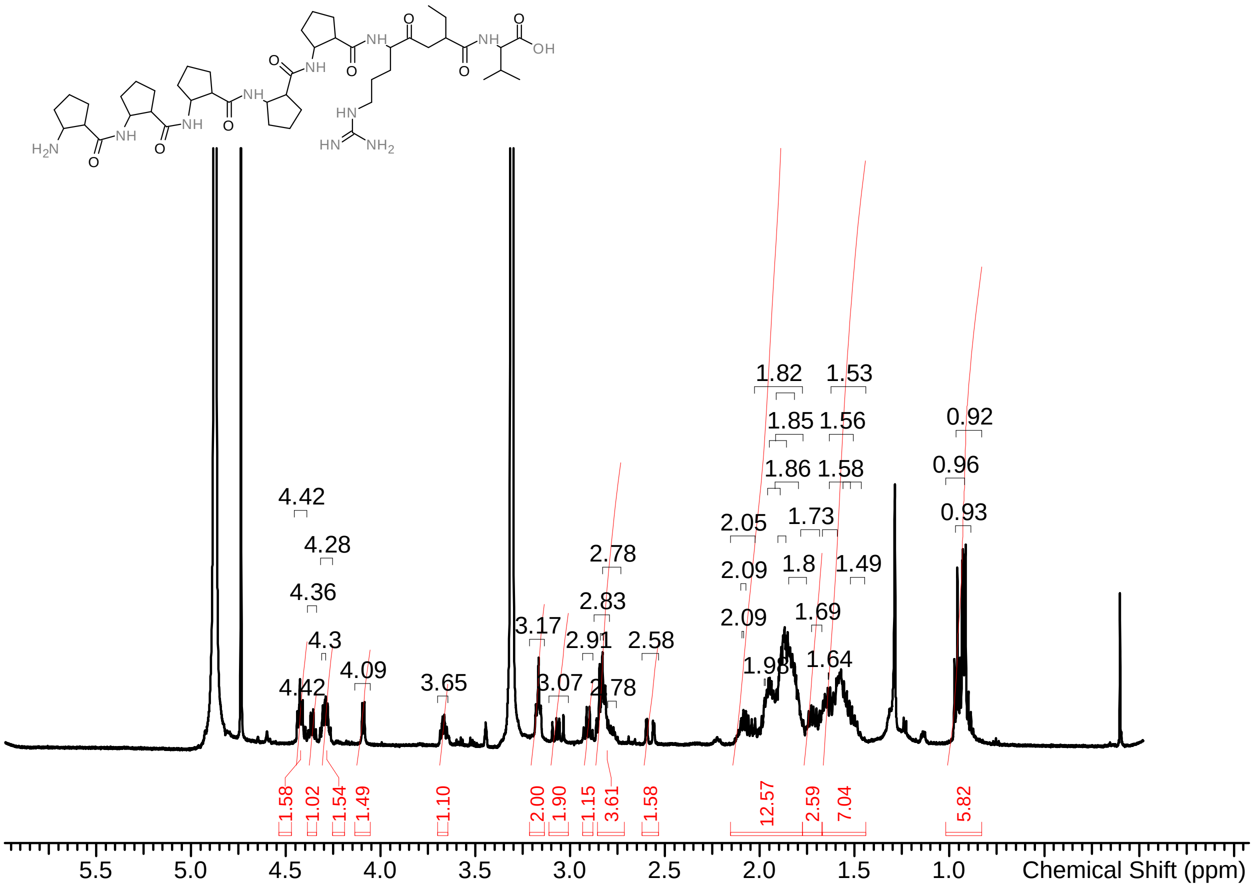

**Supplementary Figure 50** $^{1}$H-spectrum of myxopentacin A in MeOD-$d_4$ at 500 MHz.

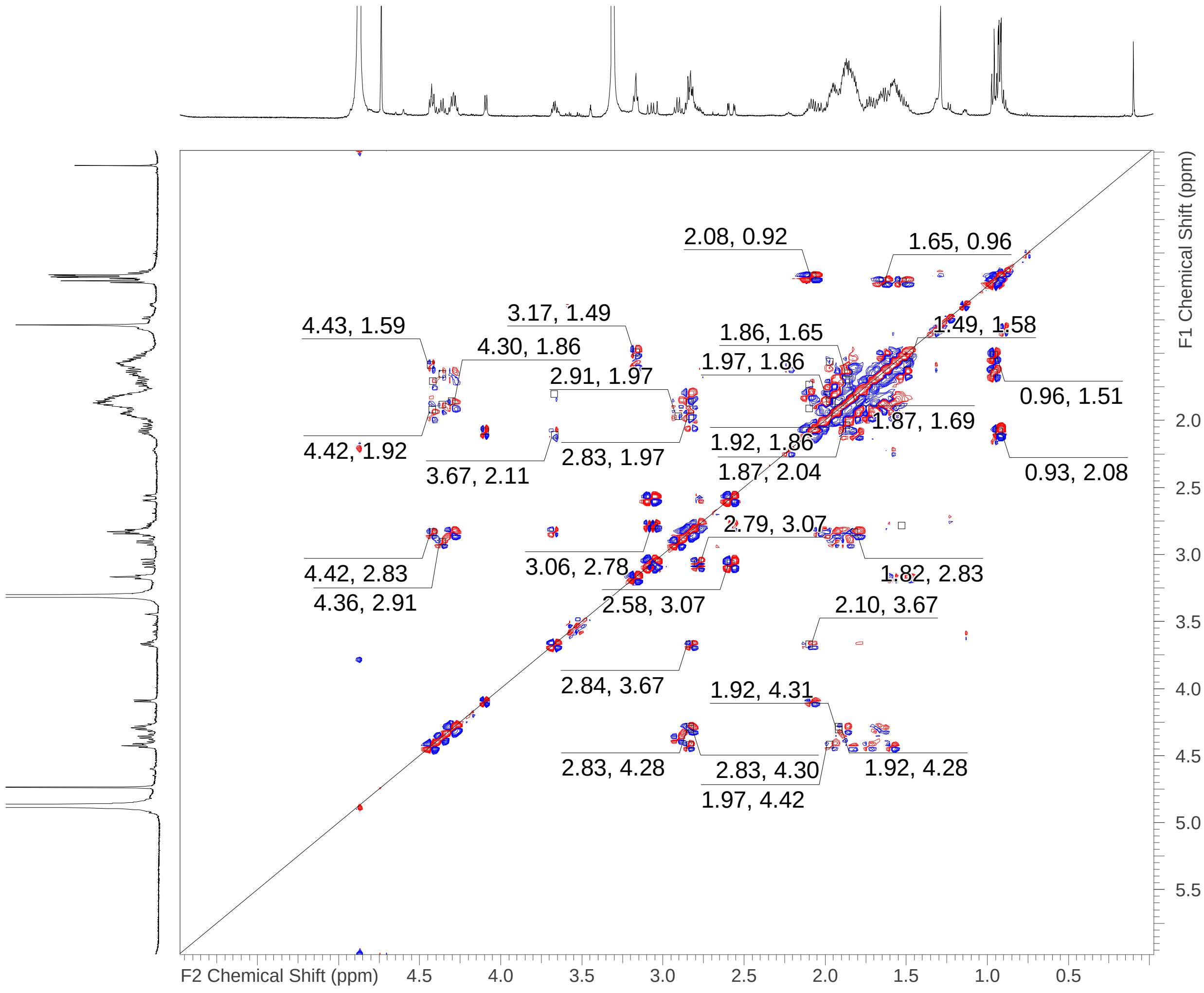

**Supplementary Figure 51** COSY-spectrum of myxopentacin A in MeOD-$d_4$ at 500 MHz.

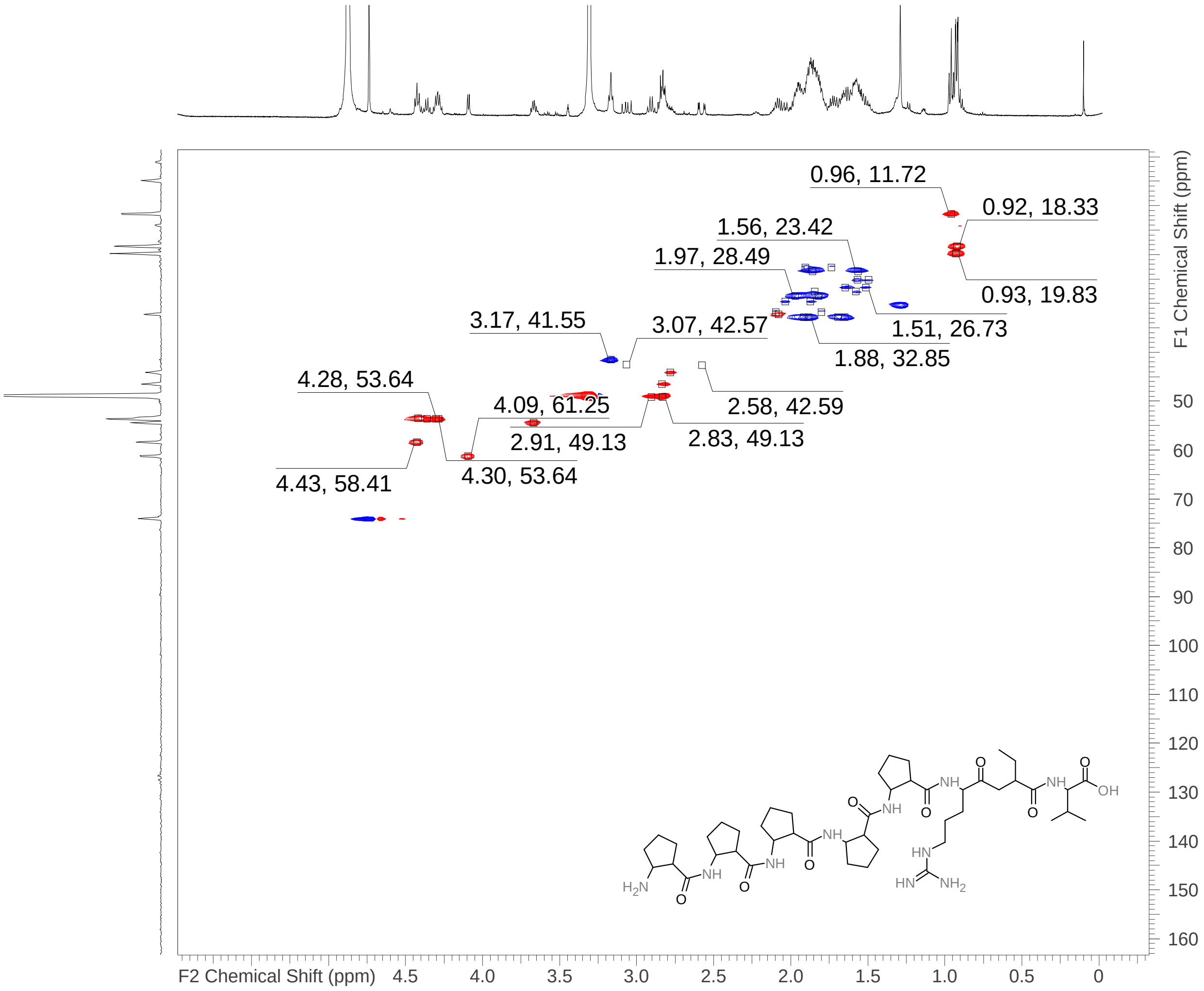

**Supplementary Figure 52** HSQC-spectrum of myxopentacin A in MeOD-$d_4$ at 500 MHz/125 MHz.

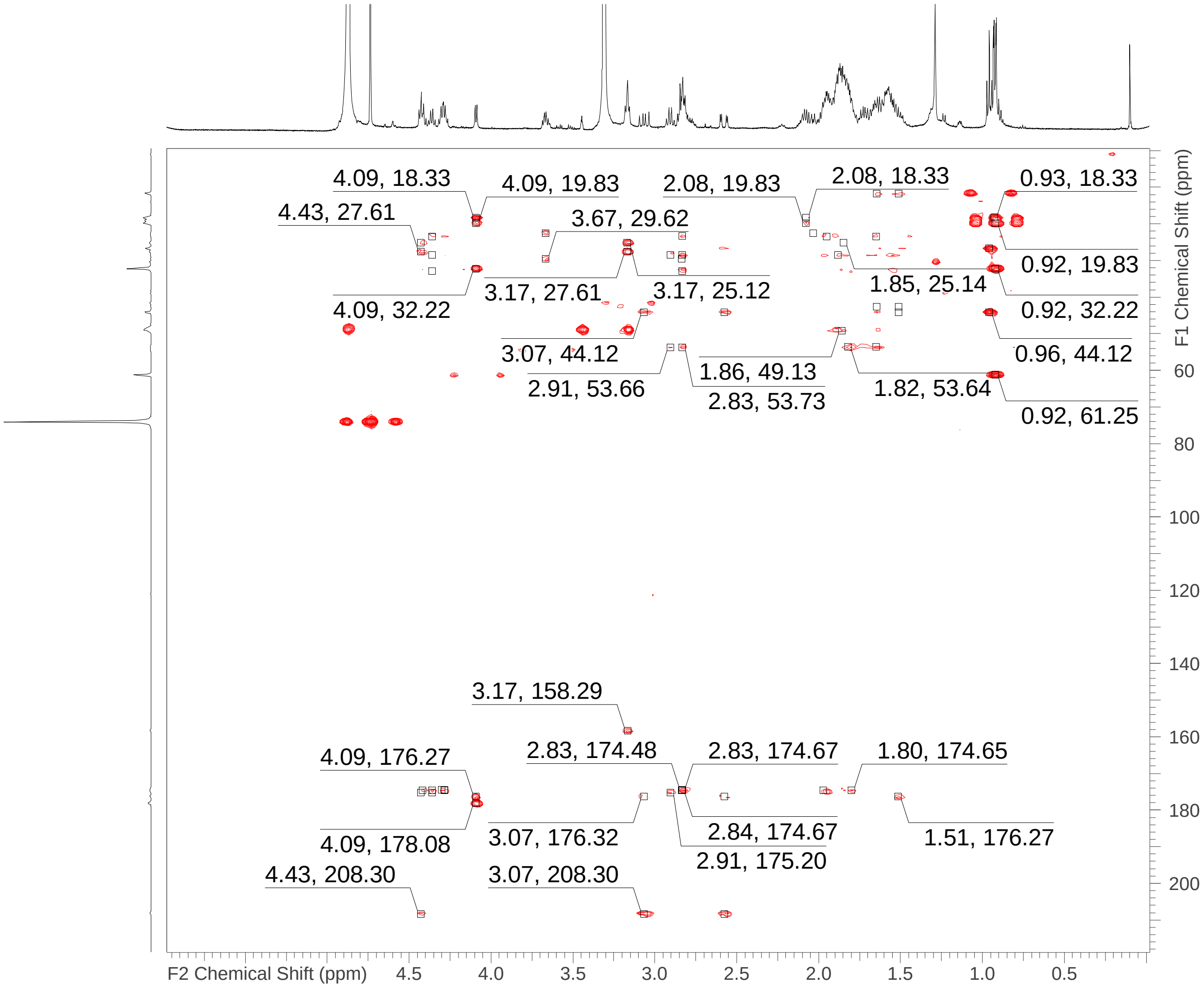

**Supplementary Figure 53** HMBC-spectrum of myxopentacin A in MeOD-$d_4$ at 500 MHz/125 MHz.

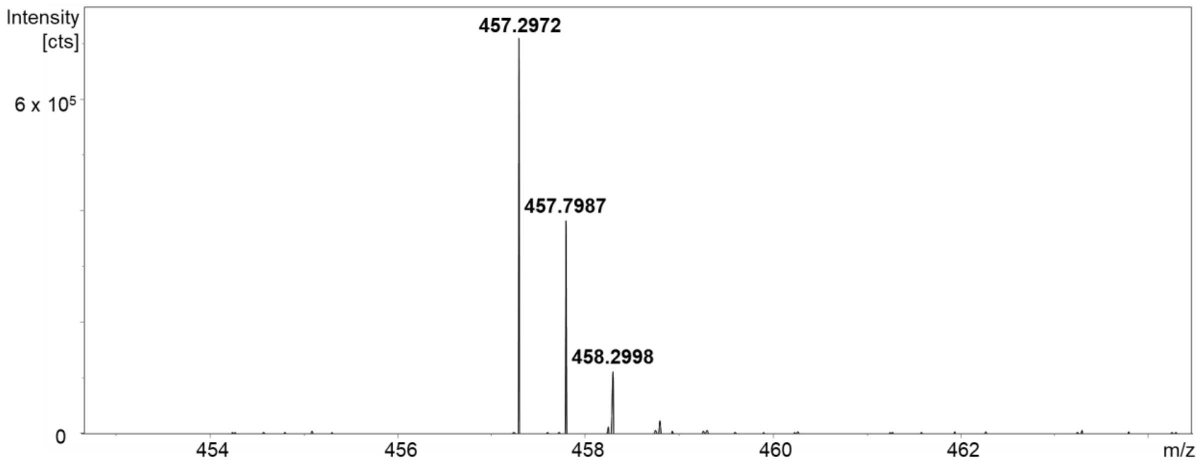

**Supplementary Figure 54** MS spectrum for myxopentacin A.

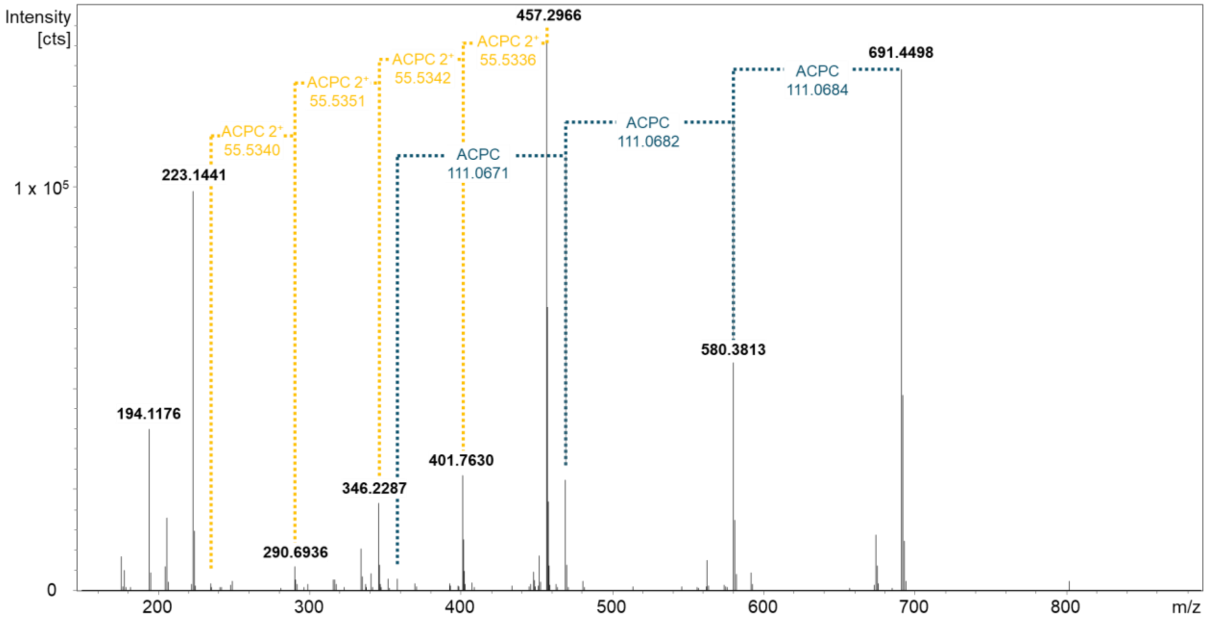

**Supplementary Figure 55** $MS^2$ fragment spectrum for myxopentacin A showing characteristic ACPC losses in both the $[M+H]^+$ (blue) and $[M+2H]^{2+}$ (yellow) series supporting the NMR-assigned structure.

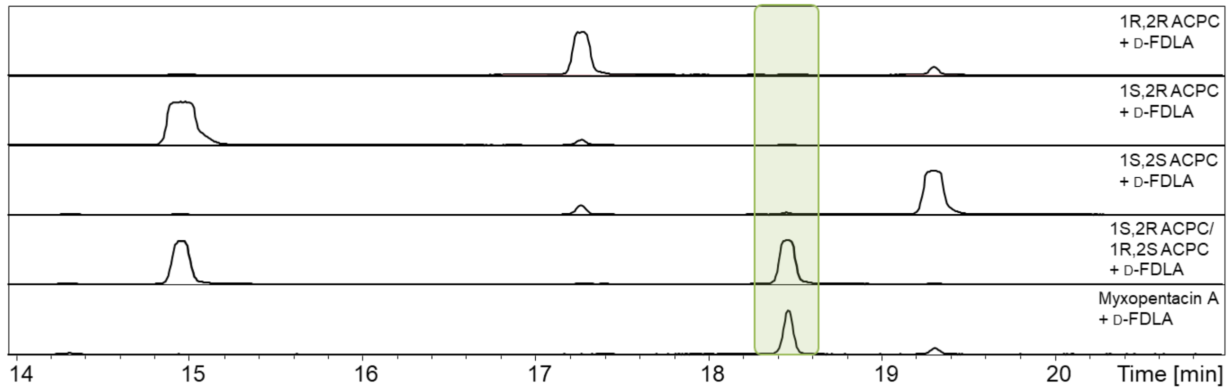

**Supplementary Figure 56** Marfey's derivatization of reference ACPC (cispentacin) with D-FDLA and L-FDLA and myxopentacin A with D-FDLA for retention time comparison resulted in assignment of 1R, 2S ACPC incorporation in myxopentacin A.

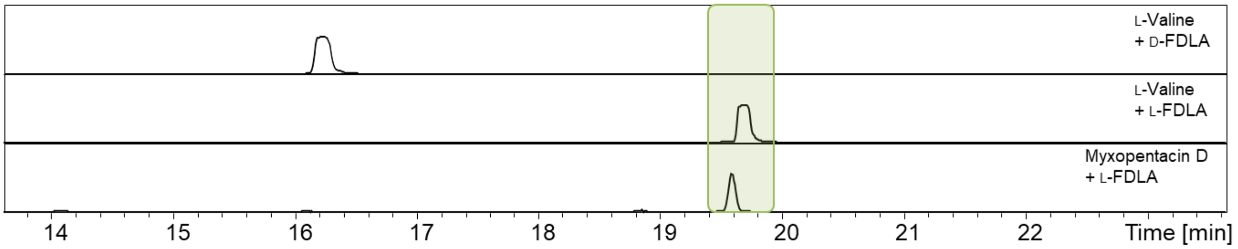

**Supplementary Figure 57** Marfey's derivatization of reference L-valine with D-FDLA and L-FDLA and myxopentacin A with with L-FDLA for retention time comparison resulted in assignment of L-valine incorporation in myxopentacin A.

**Supplementary Note 8** Biosynthesis of myxopentacin.

Making use of the NMR-based myxopentacin A structure, the minimal boundaries of the *mxp*BGC in MCy9003 were defined to span 52.2 kb that encode 21 genes named *mxpA-mxpU* (Supplementary Figure 58A, Supplementary Table 16). Beyond *mxpA-H*, the cispentacin cassette, several additional assignments can be made (Supplementary Figure 58, Supplementary Table 16). MxpA encodes a two modular NRPS with the NRPS A domain code (DPWFLGAVFK) of M01 aligning well with the observed valine incorporation (best match = valine; PARASECT[24] prediction 0.861). The NRPS A domain code of M01 (DPWFVGGVFK) suggests incorporation of an aliphatic amino acid as well with the three best hits valine, leucine and isoleucine (PARASECT prediction 0.845, 0.825, 0.787, respectively) aligning with the observed aminobutyric acid incorporation (Supplementary Figure 58B). Adjacent to *mxp*B-H, the cispentacin cassette, are *mxpJ*, *mxpK* and *mxpL*, which show significant homologies to genes in the ketomemicin pathway[25]. MxpJ shares homology (50% aa identity) with KtmA, the aldolase catalyzing an aldol reaction concomitant with decarboxylation starting from malonyl-CoA and phenylpyruvate during ketomemicin formation. We therefore expect MxpJ to catalyze a similar reaction on the dipeptide generated by MxpA after conversion of the terminal amine to a keto-function (Supplementary Figure 58B), but were unable to pinpoint the enzyme performing the required transamination reaction. MxpL shares homology (62% aa identity) with KtmC, the dehydratase reducing -OH group generated by the aldolase reaction and is therefore expected to perform a similar reaction on the building block generated by MxpJ in myxopentacin biosynthesis. *MxpB* encodes six NRPS modules aligning well with the observed incorporation of five ACPC and one arginine building block in myxopentacin A (Supplementary Figure 58C). However, the function of the encoded PKS module (M03) remains elusive. The NRPS A domain code of M04 and M06 (DVWHVSLVDK) both have serine as a best matching hit in PARASECT (1.000) aligning with the fact that serine and ACPC both show hydrophilic substituents in β-position. Interestingly, the NRPS A domain codes in M05, M07 and M08 differ from M04 and M06 (DILQLGLVWK, DAWGQAFIDK and DAFFLGFTYK, respectively) without any good matches of their predicted specificities to the observed ACPC incorporation. The NRPS A domain code of M09 (DAEDIGTVVK) suggests incorporation of an alkaline amino acid with best hits being lysine, ornithine or arginine (PARASECT prediction 0.938, 0.827, 0.797, respectively) aligning well with the observed arginine incorporation. Lastly, MxpK shares homology (59% aa identity) with KtmB, a PLP-dependent amino acid C-acyltransferase catalyzing the Claisen-type condensation between phenylalanine and benzylfumaryl-CoA in ketomemicin and expected to perform a similar reaction on the two building blocks generated from MxpA + MxpJ + MxpL and MxpU + MxpK to form the final myxopentacin A.

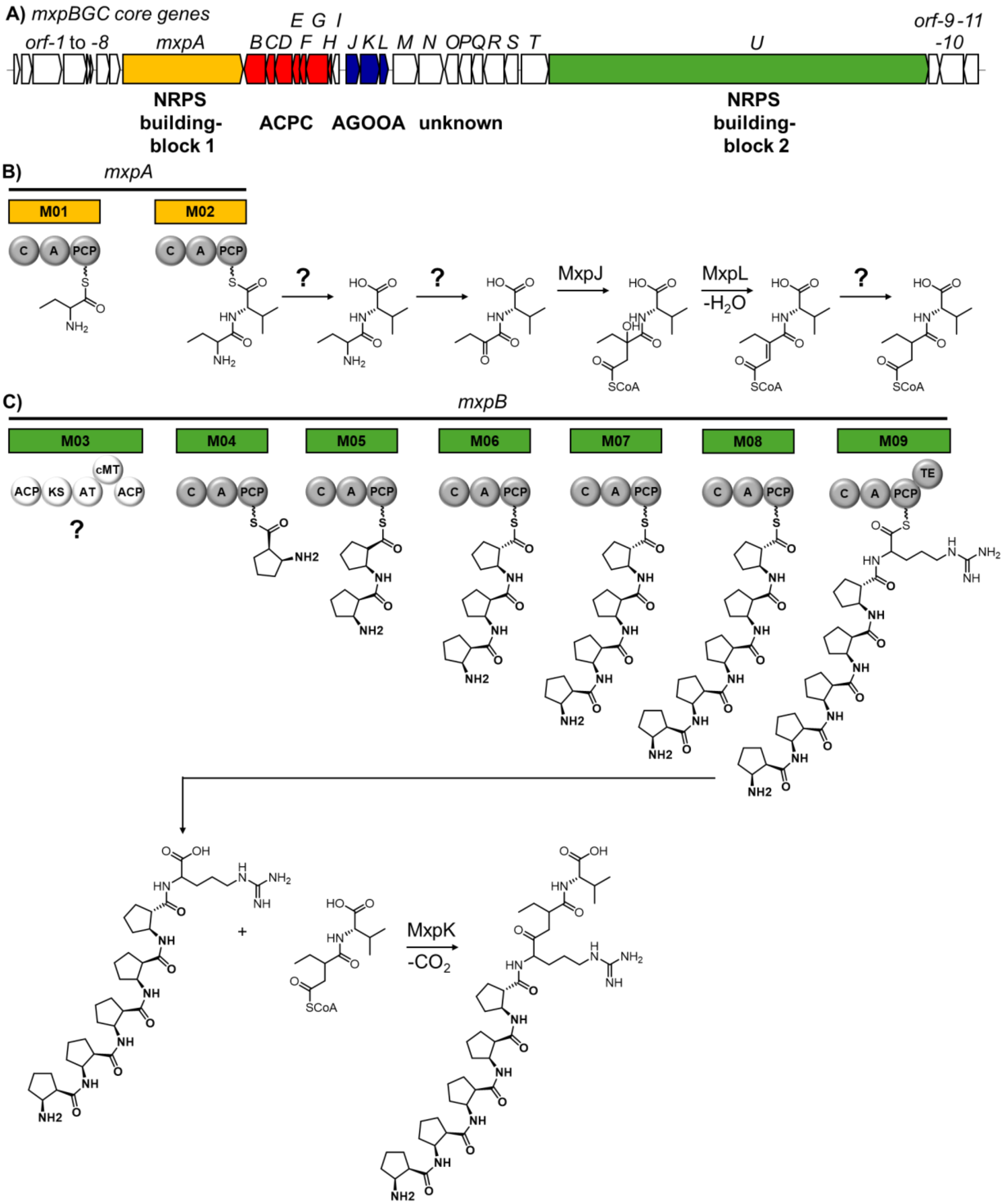

**Supplementary Figure 58** *mxp* BGC from *Myxococcus* MCy9003 with proposed functions of the enzymes encoded by the NRPS core and some of the accompanying genes. **A** Besides the cispentacin cassette used for genome mining (*mxpB-H*), the minimal *mxp*BGC includes two NRPS cassettes (*mxpA*, yellow and *mxpU* green) as well as a three gene cassette (*mxpJ-L*) likely involved in formation of the AGOOA building block observed in myxopentacin A. White = Genes encoding proteins with unknown function. **B** The C-terminal end of myxopentacin A is likely initiated by the two modules of *mxpA*, with subsequent elongation by MxpJ and L. **C** The six modules of *mxpU* are predicted to form the cispentacin chain with a final arginine elongation before release from the assembly line and coupling to the building block shown in **B**.

**Supplementary Table 16** Genes detected in the *mxp*BGC from *Myxococcus* MCy9003 with their closest protein homologues.

| Name | Size [AA] | Proposed function/ homologue | Organism | Accession number of closest protein homologue | Sequence identity [%] |
|---|---|---|---|---|---|
| *mxpA* | 2593 | NRPS, Gramicidin S synthase 2 | *Brevibacillus brevis* | P0C064.2 | 37% |
| *mxpB* | 469 | 2-succinyl-benzoate-CoA ligase | *Bacillus licheniformis* | Q65FT5.1 | 24% |
| *mxpC* | 194 | Phosphoribosylformimino-5-aminoimidazole carboxamide ribotide isomerase | *Rippkaea orientalis* | B7K3A9.1 | 26% |
| *mxpD* | 374 | Beta-ketoacyl-ACP synthase | *Mycobacterium tuberculosi* | H8ESN0.2 | 42% |
| *mxpE* | 174 | Coronafacic acid dehydratase | *Pseudomonas savastanoi* | P72238.1 | 53% |
| *mxpF* | 148 | Acyl-CoA thioesterase YbgC | *Haemophilus influenzae* | P44679.1 | 29% |
| *mxpG* | 465 | Ornithine aminotransferase | *Brevibacillus brevis* | C0ZBR4.1 | 29% |
| *mxpH* | 89 | D-alanyl carrier protein | *Streptococcus sanguinis* | A3CR85.1 | 40% |
| *mxpI* | 153 | Cys-tRNA(Pro)/Cys-tRNA(Cys) deacylase YbaK | *Haemophilus influenzae* | P45202.2 | 31% |
| *mxpJ* | 282 | Citrate lyase subunit beta | *Klebsiella pneumoniae* | P17725.2 | 32% |
| *mxpK* | 410 | 8-amino-7-oxononanoate synthase | *Marinomonas sp.* | A6W0Y0.1 | 27% |
| *mxpL* | 178 | Mesaconyl-CoA hydratase | *Haloarcula marismortui* | Q5V464.2 | 30% |
| *mxpM* | 529 | 4-amino-L-phenylalanyl-[CmlP-peptidyl-carrier-protein] 3-hydroxylase | *Streptomyces venezuela* | F2RB80.1 | 38% |
| *mxpN* | 534 | 4-amino-L-phenylalanyl-[CmlP-peptidyl-carrier-protein] 3-hydroxylase | *Streptomyces venezuelae* | F2RB80.1 | 32% |
| *mxpO* | 293 | CAAX prenyl protease 2 | *Methanococcus maripaludis* | Q6LZY8.1 | 27% |

| | | | | | |
|---|---|---|---|---|---|
| *mxpP* | 272 | Aspartyl/asparaginyl beta-hydroxylase | *Homo sapiens* | Q12797.3 | 40% |
| *mxpQ* | 243 | Methyltransferase sdnD | *Sordaria araneosa* | A0A1B4XBG9.1 | 26% |
| *mxpR* | 435 | Macrolide efflux protein A | *Streptococcus pyogenes* | P95827.1 | 26% |
| *mxpS* | 305 | 2-oxoglutarate and iron-dependent oxygenase JMJD | *Gallus gallus* | Q5ZHV5.1 | 24% |
| *mxpT* | 592 | Medium-chain-fatty-acid-[acyl-carrier-protein] ligase JamA | *Moorena producens* | Q6E7K9.1 | 50% |
| *mxpU* | 8206 | NRPS, Tyrocidine synthase 3 | *Brevibacillus parabrevis* | O30409.1 | 37% |

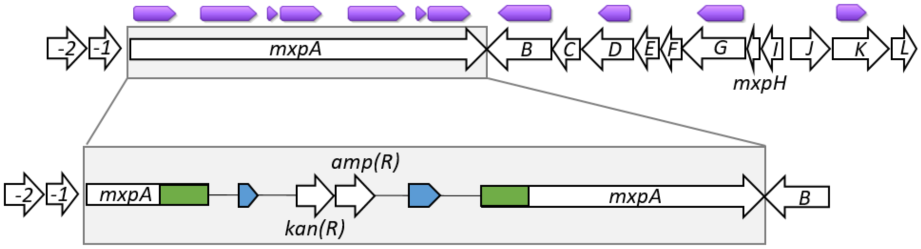

**Supplementary Figure 59** Inactivation of mxpA. The plasmid pCR2.1-001inact39Ao1 was inserted into mxpA via homologous recombination using a PCR amplified fragment (001inact39A) from MCy9003 genomic DNA using 001-inact-nrps39A-F and 001-inact-nrps39A-R primers (see Supplementary Table 17). The orientation of the homology region in a plasmid was determined using PCR21R in combination with 001-inact-nrps39A-F and 001-inact-nrps39A-R primer (Supplementary Table 17). For confirmation of the homology region in the plasmid, the primer G001-inact39Ao1 (Supplementary Table 17) aligning in the vicinity of the homology region was used. Homology arms shown in green, origins of replication in blue and selection markers for ampicillin and kanamycin labelled as *kan(R)* and *amp(R).* Catalytic domains for the genes mxpA-mxgL are shown in purple.

**Supplementary Table 17** Plasmids and oligonucleotides used in this study.

| Plasmid name | Details | Reference |
|---|---|---|
| pCR2.1-TOPO | ori(pUC), ori(f1), kanamycin resistance (kanR), ampicilin resistance (ampR) | TOPO cloning kit |
| pCR2.1 001inact39Ao1 | 1 kb homology from the first A domain of mxpA gene cloned into pCR2.1 in forward orientation | This study |
| **Primer** | **5'-Sequence-3'** | **Construct/Usage** |
| 001-inact-nrps39A-F | GTGTCGGGCGCTCGGTC | pCR2.1 001inact39Ao1 |
| 001-inact-nrps39A-R | CACGGATCTTCACCTGGAGGTC | |
| PCR21R | ATAGGGCGAATTGGGCCCT | Reverse primer on the backbone of pCR2.1 vector |
| G001-inact39Ao1 | CCATCTCCAGCGCGATGAC | Aligning on the genome located in front of pCR2.1-000inact39Ao1 plasmid |

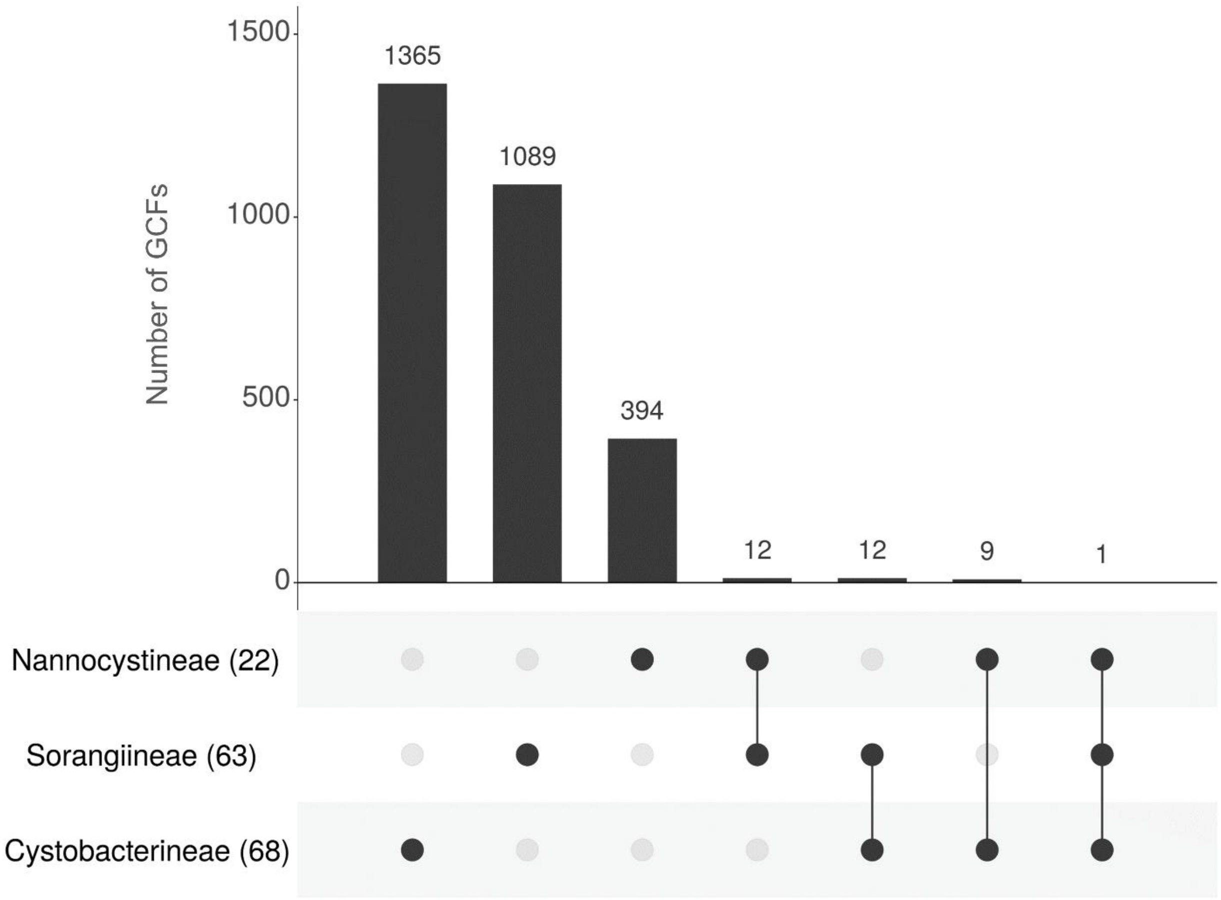

**Supplementary Figure 60** Upset plot showing the number of unique and shared GCFs between different sub-orders belonging to *Myxococcota* phylum.
